# Trait Anxiety Effects on Late Phase Threatening Speech Processing: Evidence from Electroencephalography

**DOI:** 10.1101/2020.10.02.323642

**Authors:** Simon Busch-Moreno, Jyrki Tuomainen, David Vinson

**Affiliations:** Division of Psychology and Language Sciences, University College London, London, United Kingdom

**Keywords:** prosody, semantics, anxiety, emotion, cognition, electrophysiology

## Abstract

The effects of threatening stimuli, including threatening language, on trait anxiety have been widely studied. However, whether anxiety levels have a direct effect on language processing has not been so consistently explored. The present study focuses on eventrelated potential (ERP) patterns resulting from electroencephalographic (EEG) measurement of participants’ (n = 36) brain activity while they perform a dichotic listening task. Participants’ anxiety level was measured via a behavioural inhibition system scale (BIS). Later, participants listened to dichotically paired sentences, one neutral and the other threatening, and indicated at which ear they heard the threatening stimulus. Threatening sentences expressed threat semantically-only, prosodically-only, or both combined (congruent threat). ERPs showed a late positivity, interpreted as a late positive complex (LPC). Results from Bayesian hierarchical models provided strong support for an association between LPC and BIS score. This was interpreted as an effect of trait anxiety on deliberation processes. We discuss two possible interpretations. On the one hand, verbal repetitive thinking, as associated with anxious rumination and worry, can be the mechanism disrupting late phase deliberation processes. Instantiated by inner speech, verbal repetitive thinking might be the vehicle of anxiety-related reappraisal and/or rehearsal. On the other hand, increased BIS could be simply affecting an extended evaluation stage as proposed by multistep models, maybe due to over-engagement with threat or to task-related effects.

**Graphical Abstract:** 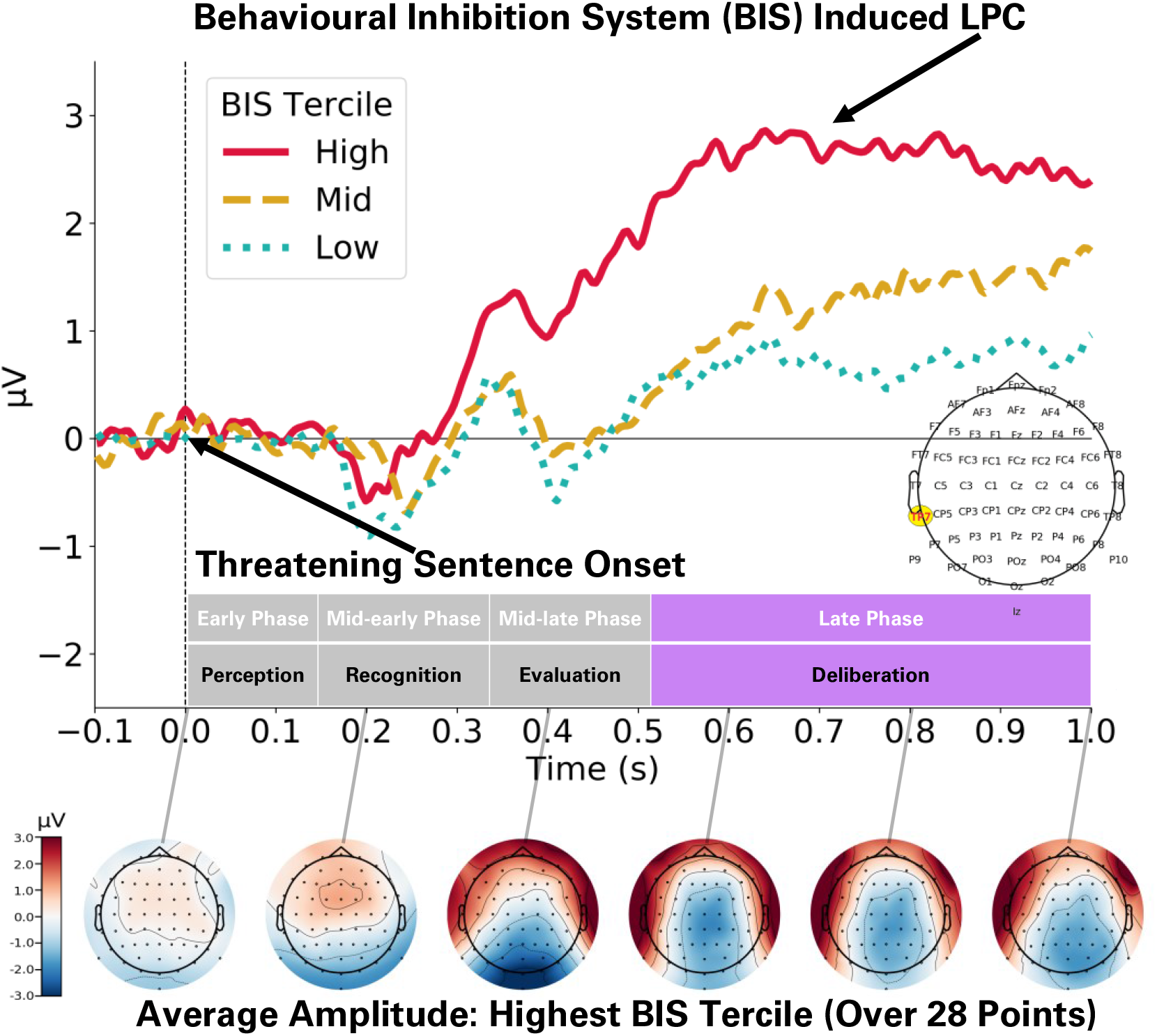

---

Both anxiety and language processing are characterized by well-defined lateralization patterns. Anxiety has been widely associated to a dual processing pattern: attention-related arousal and fear-related responses tend to involve greater right hemisphere (RH) processing, while instead evaluation-related apprehension and inhibition-related responses tend to involve greater left hemisphere (LH) processing (Heller et al., 1997; Nietschke et al., 2000; Spielberg et al, 2013). According to current models of speech processing (i.e. dual stream model), speech comprehension recruits both hemispheres (Kemmerer, 2015). While more RH involvement is required for slow rate suprasegmental processing of prosody and/or affect recognition, more LH involvement is required for fast rate segmental processing and/or lexical categorization (Belin et al., 2004; Liebenthal et al., 2010; Poeppel et al., 2008; Zatorre et al., 2002). These observations speak not only of general hemispheric activity, but also of very specific activity patterns and brain anatomical structures that, in many cases, are shared by language and anxiety processing. This opens the question of whether these processes simply co-occur, showing superficial similarities, or actually interact with each other.

An additional element of complexity is that anatomical differences are not necessarily absolute or static, but they may vary over time, as evidenced by temporal changes in lateralization of emotional prosody and semantics lateralization patterns (Kotz and Paulmann, 2011). Similarly, anxiety may modulate processing of threat differently at early and late processing phases, with early over-attention to threat and later over-engagement with threat (Bar-Haim et al., 2007). Here, over-attention implies facilitated attention to threatening stimuli, and over-engagement implies difficulty in disengaging from threatening stimuli (Cisler and Koster, 2010). Therefore, understanding these anxiety-related attention processes as multistage processes might be crucial. A two-stage model proposes that trait anxious personality is characterized by an over-active behavioural inhibition system (BIS), when attempting to resolve approach/avoidance conflicts. This would develop in two stages: 1) valuation of stimuli immediately after stimulus input, and 2) motivation for behaviour as approach or withdrawal (Corr and McNaughton, 2012). However, this model does not give an account of processes that might occur between valuation and motivation, as proposed in a multistage model of anxiety (Bar-Haim et al, 2007). In this model, anxiety develops through four processing stages: 1) pre-attentive threat evaluation, 2) resource allocation, 3) guided threat evaluation, 4) goal engagement system. Here, a valuation input can affect early pre-attentive stages, which can account for fear responses inducing rapid withdrawal (output), or fight-flight-freeze system (FFFS) responses (Corr and McNaughton, 2012). Later stages can involve resource allocation (output), which can lead to behavioural inhibition (or BIS engagement) in a threat evaluation stage. Likewise, a goal engagement system can account for a motivation output that occurs after BIS activation, for either approaching or withdrawing.

An interesting parallel can be seen for emotional language processing where a multistep model has also been proposed (Kotz and Paulmann, 2011). This model proposes three main stages: 1) early stage perceptual processing, 2) mid stage recognition processing, 3) late stage evaluation processing. An important aspect of this model is that it takes into account hemispheric differences of language and emotional processing. The processing of acoustic properties would involve activity at early stages, where different types of information would be associated with hemispheric differences. In particular, greater RH engagement would be associated to prosodic slower rate spectral information and greater LH engagement would be associated to phonological (and by extension semantic) faster rate temporal information (Poeppel et al., 2008; Zatorre et al., 2002). Mid stages might involve the emotional recognition of stimuli, implying hemispheric differences varying with stimulus type and/or conveyed emotion (Schirmer and Kotz, 2006). Late stages would be associated to informational integration and evaluation of emotional stimuli (Kotz and Paulmann, 2011). If indeed the similarities between the multistage model of anxiety and the multistep model of language are more than coincidental, the latter should also have a fourth stage, which can account for motivation/goal-orientation processes associated to behavioural outputs in parallel to the multistage model of anxiety.

Nevertheless, to develop such a model, more evidence is necessary about how informational properties of language interact with intrinsic affect. Given that lexical items and prosody can convey emotional meaning at the same time (Nygaard et al., 2009; Schirmer and Kotz, 2003), the simultaneous recognition and integration of diverse informational properties is required, including the processing of segmental and spectral information, tonal and temporal information, and identification and categorization processes. As mentioned before, these aspects of language processing show different hemispheric asymmetries, which might be linked to anxiety related lateralisation in terms of how this information is recognized, integrated, and evaluated. Thus, a model of anxious lateralization (Heller et al., 1997) could be extended into a multistage model of anxiety (Bar-Haim et al., 2007), as early over-attention to threat and later (sustained) over-engagement with threat could present different lateralisation patterns (Spielberg et al., 2013).

Recent electroencephalography (EEG) evidence indicates that a hyperactive BIS, signalled by higher scores in BIS psychometric scales, is characterised by a right frontal hemispheric pattern (Gable et al., 2017; Neal and Gable, 2017, but see: De Pascalis et al., 2013). However, some key theoretical notions cannot be overlooked. On the theoretical side, the proposed anatomic-physiological basis of BIS/anxiety is the amygdala-hippocampal-septal route (Gray and McNaughton, 2000), which can modulate diverse cortical patterns depending on environmental conditions. Indeed, it has been proposed that the BIS limbic system might modulate higher processes such as worry and rumination through language systems (in particular left-lateralized production/rehearsal), which could induce additional inhibitory control (Gray and McNaughton, 2000). Hence, while RH may induce initial inhibitory control (McNaughton et al., 2013; Robinson et al., 2019, Neal and Gable et al., 2017), there might be later LH influence involving evaluation or deliberation processes. Previous EEG studies have observed left or bilateral frontal alpha activity associated with anxious apprehension as measured by worry (Heller et al., 1997; Nitschke et al., 1999), and bilateral alpha for rumination-correlated BIS (Keune et al., 2012).

Given these findings, manipulating hemispheric input could reveal some of the nuances of how anxiety affects threatening speech processing at different processing phases. In other words, stimuli that are processed first by LH or RH might be affected differently depending on: 1) their language informational properties (i.e. whether threat is expressed via semantics or prosody) and/or 2) participants’ intrinsic lateralization differences when processing threatening stimuli (i.e. anxiety). Considering this, dichotic listening (DL) stands out as an ideal behavioural approach to observe how anxiety-related hemispheric asymmetries might influence emotional language processing hemispheric asymmetries. This relates to the frequently observed phenomenon of a right ear advantage (REA) for non-prosodic language stimuli (Hugdahl, 2011); this implies participants answering faster and/or more accurately to stimuli presented at their right ear when compared to stimuli presented at their left ear. On the other hand, prosody, in particular emotional prosody, leads to either a left ear advantage (LEA) or at least a diminished REA (Godfrey and Grimshaw, 2015; Grimshaw et al., 2003). Few DL experiments have researched the effects of anxiety on emotional speech processing (Gadea et al., 2011). Instead, they either use speech/prosody as an emotion-eliciting stimulus or use DL mainly as an attentional manipulation technique (Bruder et al., 2004; Leshem, 2018; Peschard et al., 2016; Sander et al., 2005).

However, these behavioural tasks cannot provide evidence of effects on specific time-windows, as required by multistep models (i.e. Kotz and Paulmann, 2011). EEG measurements, via the event-related potential (ERP) technique, could help to elucidate the nuances of anxiety effects (e.g. Moser et al., 2014; Wabnitz et al., 2015). In the present study’s case, observations could reveal whether early and/or late ERPs differ given ear, sentence-type and task; and how this may relate to over-attention or over-engagement processes in anxiety (increased BIS). To our knowledge, no EEG experiment has integrated trait anxiety measures with DL to investigate the effects of anxiety on threatening semantics and prosody. In the present study we implement a dichotic listening experiment where participants answer to threatening sentences containing only semantic threat, only prosodic threat, or both (semantic-prosodic congruency). Participants were required to answer only after the end of each sentence, to avoid motor response contamination of the EEG signal during listening. The experiment was comprised of direct- and indirect-threat tasks; where the latter implies answering to the neutral stimulus of the dichotic pair (intended as a control). As a proxy of trait anxiety we used a BIS scale from the Reinforcement Sensitivity Theory Personality Questionnaire (RST-PQ) (Corr and Cooper, 2016). This psychometric measure, well grounded in physiological research, is well suited to to our present theoretical framework and experimental aims. Finally, via EEG measurement, we can provide information on relevant ERPs at different processing time-windows, crucial for testing our theoretical model.

To broadly summarise our hypotheses, we expect that higher levels of anxiety (higher BIS scores) will induce early over-attention to threat but mid-late over-engagement with threat. The first should be associated with anxiety induced arousal, which has been observed to be right lateralized (Heller et al., 1997; Neo et al., 2011). The second should be associated to a left lateralized or bilateral pattern (Nitschke et al, 1999), which might be due to mid-stage LH thought-induced (e.g. worry or rumination) evaluation and inhibition (Spielberg et al., 2013), but a later RH involvement associated to arousal induced by sustained state anxiety (McNaughton and Gray, 2000). To test these, we established four time-windows where differences may be present: 1) pre-attentive (50-150ms), 2) attentive (150-250ms), 3) evaluative (250-500ms), 4) orientative (500-750ms). These were selected *a priori* as non-overlapping intervals that span over relevant amplitude peaks. Although previous literature does not directly specify these intervals, the emotional language literature does specify a number of ERPs peaking at these relevant time-points: 100ms (sensorial perceptual stage), 200ms (recognition stage), ∼300-600ms (evaluation stage) (Kotz and Paulmann, 2011).

Our first two time-windows map closely onto these first two stages, with the third and fourth designed in order to tap into the proposed fourth step of emotional language processing. We propose that anxiety-associated activity around 600ms would not reflect evaluation processes, but deliberation/orientation (not included in the multistep model of emotional language processing). For instance, late positive potentials (LPP) have been proposed as signaling rehearsal processes during emotional processing in anxiety (Hajcak et al., 2010). Time-windows for these ERPs vary widely, usually going from around 400ms to 800ms to 1s, and they tend to overlap with P300, also defined with wide variability in anxiety-related studies (Hajcak et al., 2010). Although they may differ in topology, these ERPs have been proposed as signaling the same or similar processes (e.g. Sassenhagen et al., 2014). Nevertheless, this overlap and wide variability may also be a problem associated with the selection of analysis time-windows by eye-balling the ERP grand averages, rendering systematic differences in reported averaged time-windows for analysis. For this reason, the use of automatic window selection or the a priori selection of time-windows is preferred, and in the case of no clear previous information about time-windows (as in our present case) the a priori segmentation of the epoch in sequential and narrow time-windows is recommended (Luck et al., 2014).

Given that we are not aware of sufficient previous studies investigating dichotic listening, threatening prosody/semantics and anxiety (BIS) in simultaneity, we opted out for establishing sequential a priori time-windows for analysis (as described above). Also, as present research has no direct precedents in the literature, it is not clear whether observed ERPs and their amplitude differences by condition in such time-windows will be coincidental with those observed in previous research, which regards DL, emotional speech, or anxiety, but not all three combined. Hence, we prefer to refrain from predicting effects on specific ERPs, and instead we propose the following specific predictions: In windows 1 and 2 anxiety should affect speech processing (Pell et al., 2015), due to anxious over-attention to threat. This would increase RH involvement for prosody, facilitating detection of stimuli presented contralaterally (left ear). In window 3 the effects of anxiety should result from over-engagement with threat (Spielberg et al., 2013); slowing down LH processing of semantic stimuli (right ear presentation). In window 4, anxiety should affect goal orientation (Bar-Haim, 2007), where both hemispheres should be comparably affected (no ear differences). This could be the result of a worry-arousal loop (McNaughton and Gray, 2000) due to continued exposure to threatening stimuli, or could result from delayed disengagement from threat (Cisler and Koster, 2010) or sustained attention to emotional stimuli (Hajcak et al., 2010), which can be also understood as anxiety-related over-engagement with threat.

## Methods

### Participants

Participants (n = 36, mean age = 28.6, age range = [19, 54], 19 females) were recruited using Sona Systems (sona-systems.com). Only participants reporting being right-handed, having English as first language, without hearing problems and with no history of neurological/psychiatric disorders were recruited. Participants were remunerated at a £7.5/hour rate. All participants were informed about their rights, including those regarding their data (protected under GDPR protocols), and gave their informed consent before participating. They received a debriefing statement at the end of the experiment. The study was carried out under approval from the UCL Research Ethics Committee in accord with the Declaration of Helsinki.

### Materials

Four types of sentences were recorded: Prosody (neutral-semantics and threatening-prosody), Semantic (threatening-semantics and neutral-prosody), Congruent (threatening-semantics and threatening-prosody), and Neutral (neutral-semantics and neutral-prosody). Sentences were extracted from movie subtitles by matching them with a list of normed threatening words from the extended Affective Norms for English Words (ANEW) (Warriner et al., 2013). For the present study, any word over 5 points in the arousal scale, and below 5 points in the valence and dominance scales was considered semantically threatening. Every word with less than 5 arousal points and between 4 and 6 valence points was considered semantically neutral. Words’ frequencies were extracted from SUBTLEX-UK (van Heuven et al., 2014), only sentences containing words with Zipf log frequencies over 3 were included.

Sentences were recorded in an acoustically isolated chamber with a RODE NT1-A1 microphone. The speaker was instructed to speak in what he considered his own threatening/angry or neutral voice for recording Prosody/Congruent and Semantic/Neutral sentences respectively. Table 1 summarises sentences’ word length and duration. Figure 1 shows oscillograms and spectrograms of four sentences used as stimuli. All threatening sentences were paired with a different Neutral sentence of similar length; minor increases of silences (up to 40ms) were applied to make paired sentences’ durations match perfectly. This resulted in 54 recorded sentences per threatening category (Prosody, Semantic, Congruent), each one paired with a different sentence from the Neutral category. This provided 162 pairings of two sentences (one threatening and one neutral). We created two versions of each pair: one with threat at the right ear, and one with threat at the left ear: 324 distinct stimuli in total.

**Table 1.**
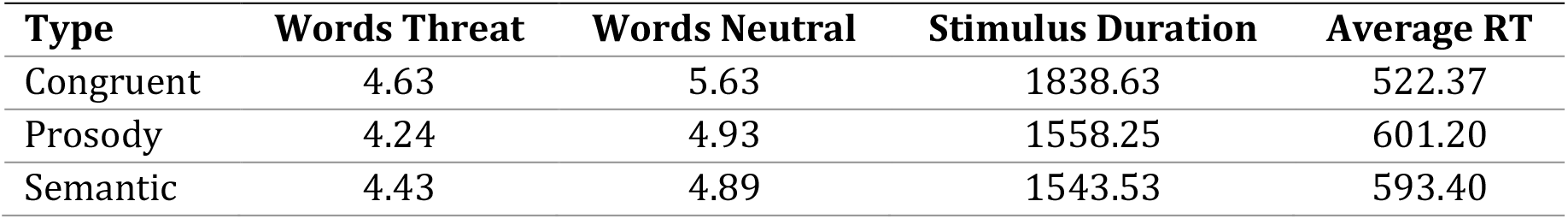
Average number of words, duration and reaction time per stimulus type

| Type | Words Threat | Words Neutral | Stimulus Duration | Average RT |
| --- | --- | --- | --- | --- |
| Congruent | 4.63 | 5.63 | 1838.63 | 522.37 |
| Prosody | 4.24 | 4.93 | 1558.25 | 601.20 |
| Semantic | 4.43 | 4.89 | 1543.53 | 593.40 |

**Figure 1.**
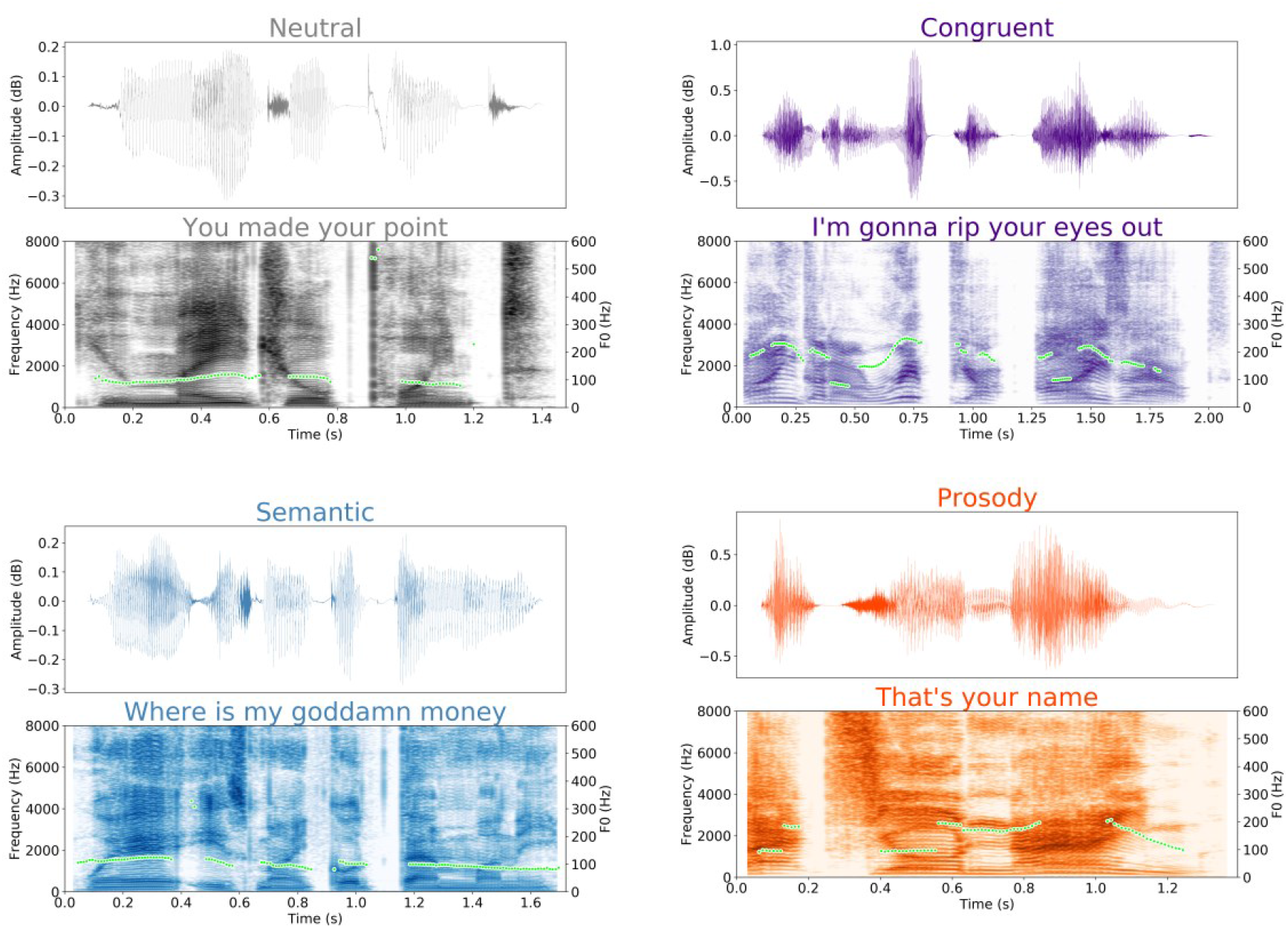
Examples of sentences with different semantic content and prosody. Top of each image: oscillogram. Bottom of each image: spectrograms. Top left: neutral prosody and neutral semantics (Neutral). Top right: threatening prosody and threatening semantics (Congruent). Bottom left: neutral prosody and threatening semantics (Semantic). Bottom right: threatening prosody and neutral semantics (Prosody). Green dots indicate fundamental frequency (F0) contours.

In addition, we assessed the prosodic properties of threatening sentences as compared to neutral sentences. Threatening prosody has been observed to align with hot anger (Banse and Scherer, 1996; Hammerschmidt and Jürgens, 2007) and can be defined mainly by pitch and roughness (voice quality). Thus, we focused only on Median Pitch (median fundamental frequency of each sentence) and Hammarberg Index (difference between 0Hz to 2000Hz and 2000Hz to 5000Hz frequency ranges), as these acoustic measures can capture pitch (fundamental frequency) and voice quality variation respectively within the spectral domain. Measures were extracted from sentences with Python’s Parselmouth interface to Praat (Jadoul et al., 2018). Acoustic measures were compared by using BEST (Kruschke, 2013), but using *SD* ∼ *Exponential* (*λ*=2) instead of uniform distributions. All models were sampled with MCM and used 1000 tuning samples and 1000 samples, convergence and sampling were excellent for both models (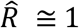, ESS > 400, BFMI > 0.8). As very clear in Table 2, sentences containing threatening prosody have higher median pitch and lower voice quality (Hammarberg index, i.e. more roughness) than sentences without threatening prosody. For both measures, Prosody and Congruent distributions greatly overlap and Semantic and Neutral sentences almost completely overlap.

**Table 2.**
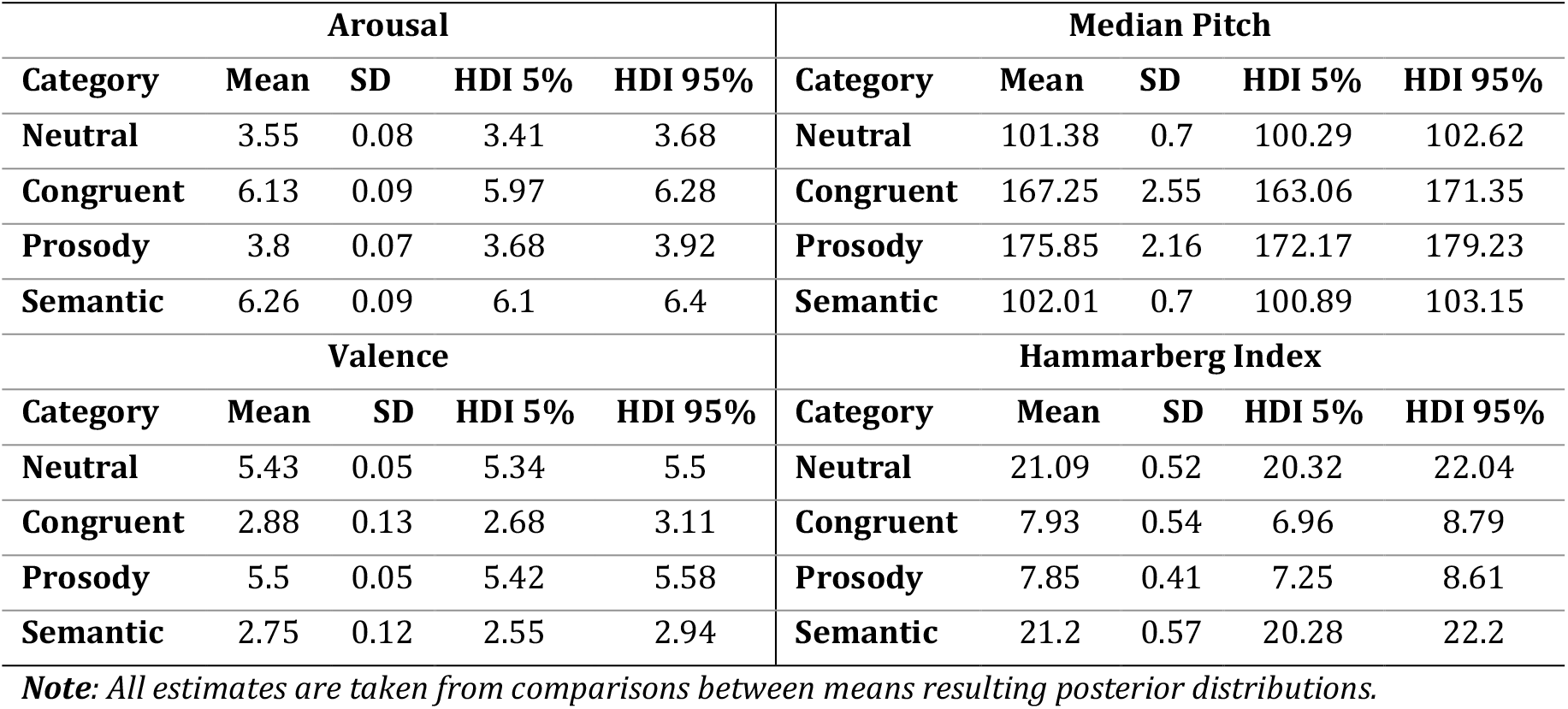
Estimates from BEST Model

Finally, we collected sentences’ threat ratings post-experimentally, for further confirming effects of both acoustic measures and ANEW norms. Ratings were given by 10 participants for 54 Prosody and 54 Neutral sentences (total number of Prosody and Neutral) in a 9-point Likert scale (0 = not threatening at all, 8 = very threatening). The same procedure was followed for the 54 Semantic sentences together with the same 54 Neutral sentences in a second rating session by 10 participants. Ratings were analysed by using a hierarchical Ordered-logistic model, with subjects as a varying intercept, eight scale cutpoints and normal priors (equivalent to L2 regularisation) for non-varying slopes for Pitch, Roughness, Arousal and Valence. The full model and plots can be found in our OSF repository (https://osf.io/n5b6h/). Figure 2 summarises posterior distributions as probabilities derived from the ordered-logistic model. For the Prosody task, these indicate that when Median Pitch (MP) is high and Hammarberg Index (HI) low, the probability of rating sentences over 6 on the threat scale increases by over 40%. For the Semantic task, when Arousal is high and Valence is low the probability of rating sentences over 7 on the threat scale increases over 60%. In both tasks when MP/Arousal are low and HI/Valence are high the probability of giving 1 or less on the threat scale increases by around 60%. The models also indicate that Arousal/Valence had no relevant effects on Prosody and MP/HI had no relevant effects on Semantic. See supplementary materials for tables with more details (Supplement 1, supplementary tables 1.1 and 1.2). All relevant materials used and their measures can also be found in supplementary materials (Supplement 1, tables 1.3 and .14).

**Figure 2.**
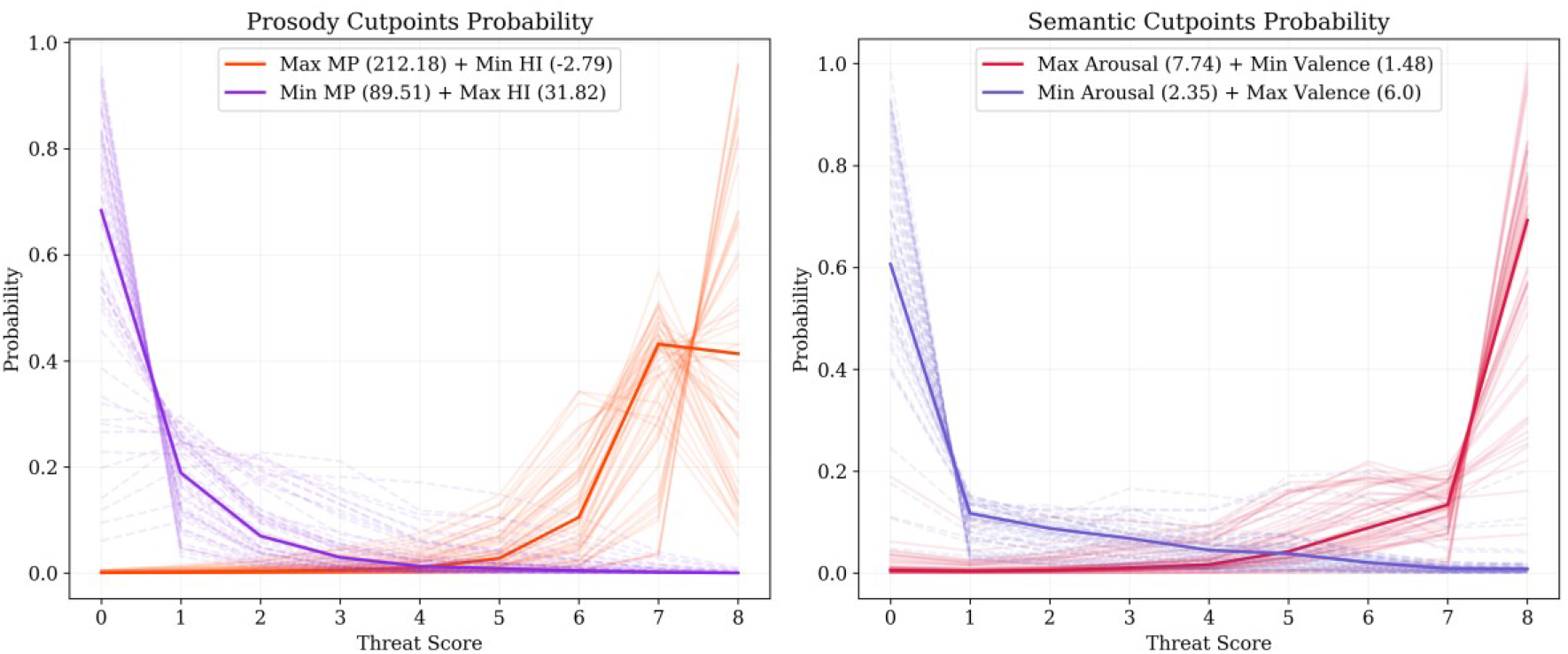
Probability of threat ratings. Left panel: probability of rating a Prosody sentence as threatening on a 0-8 Likert scale. Orange solid line: probability increase when a sentence has the Max Median Pitch (MP) and Min Hammarberg Index (HI). Violet dashed line: probability decrease when a sentence has the Min MP and Max HI. Right panel: probability of rating a Semantic sentence as threatening in a 0-8 Likert scale. Red solid line: probability increase when a sentence has the Max Arousal and Min Valence. Blue dashed line: probability decrease when a sentence has the Min Arousal and Max Valence. Faded lines are random samples from the posterior indicating uncertainty.

### Procedure

Participants were introduced to the recording room (electrically and sound isolated chamber), signed consent, and sat at 1m distance from a 20” screen used to display all tasks via PsychoPy2 (Peirce et al., 2019). Next, participants provided their demographic information (age and sex) and completed the BIS scale (m = 23, SD = 13) and FFFS scale (m = 29, SD = 9.5) (for more details see Corr and Cooper, 2016). Participants’ head dimensions were measured and EEG caps of appropriate size were placed and centered, conductivity gel was placed and the Biosemi 64 Ag/Cl electrode system (biosemi.com) was connected. Impedance levels were kept below 20Ω and electrodes were checked to be working properly. While EEG recording, participants completed a direct-threat task where they answered at which ear (left or right) they heard a threatening sentence (always paired with a neutral sentence) by pressing the left or right arrows on a keyboard (as fast and accurately as possible) only after the stimulus finished playing (sentence offset) and a target symbol appeared on screen. Trials (324 per task) started at sentence onset and were separated by a 1.5s inter-stimulus interval (ISI). In the indirect-threat task they answered to the neutral sentence of the pair in the same manner as in direct-threat. Order of task was counterbalanced across participants. Participants heard sentences twice such that a threatening sentence was presented once at each ear. Participants were requested to swap response hand every other block. Each block consisted of 18 sentences (trials), after which participants were allowed to rest. Starting hand and starting ear were also counterbalanced across participants. Due to a coding issue, in the indirect-threat condition for half of the participants (list B: starting with right ear only) stimuli were not exactly balanced for ear presentation. This issue was not noticed by participants during debriefing, it did not affect more than 4 sentences per condition per subject, and did not systematically affect the results. See supplementary materials for follow-up analyses to address this issue (Supplement 3).

### EEG Data Processing

EEG recordings were pre-processed using Python’s MNE package (Gramfort et al., 2014). A completely automated pre-processing pipeline was implemented, based on Jas et al.’s (2018) method. This consisted in the following steps:

1) Importing data, previously down-sampled to 512Hz using Biosemi decimation tools, checking that all events were correctly placed and fixing if necessary.
2) Preparing data for independent component analysis (ICA): set data to average reference (Dien, 1998; Lei and Liao, 2017), low-pass filter at 40hz to avoid aliasing artifacts, down sample to 256hz, high pass filter at 1hz for better artifact detection (both FIR filters, -6dB cut-off), and automatic rejection of noisy segments based on the Autoreject MNE package (Jas et al., 2017; Winkler et al., 2015).
3) Compute ICA components by using python Picard package (Ablin et al., 2017) for EOG artifact detection. ICA components that are highly correlated with noise (> 0.35, using MNE component correlation method) at Fpz channel were marked for later rejection.
5) Reload raw data to apply average reference to raw data (Dien, 1998; Lei & Liao, 2017); to use more relaxed filters: first a high pass (0.1Hz, FIR, -6dB cut-off) and later a low pass (100Hz, FIR, -6dB cut-off) filter (Luck, 2014; VanRullen, 2011; Widmann et al., 2015); and to exclude EOG artifacts detected in previous step (1 to 2 components were removed from each participant recording.).
6) Create epochs from 0ms (sentence onset) to 1000ms using a 100ms pre-onset period (10% of epoch). This 1s epoch allows sufficient time for observing the proposed 4 time-windows and safely avoids possible contamination from later behavioural responses (analysis of later response preparation and/or response is out of present scope and would require dynamic epoching with respect to RTs).
7) Applying automatic detection, repairing and rejection of noisy epochs by using Autoreject’s Bayesian optimization method (Jas et al., 2017). 4 to 8 channels were interpolated by participant. Between 0 and 206 trials were dropped per participant (32.75 on average), and between 29 and 54 trials were retained per condition. 54 being the total number of trials per condition (but see minor technical issue at the end of Procedure section, see also Supplement 4). So, trials are generally balanced.
8) Applying baseline correction (baseline subtraction) (Luck, 2014; Tanner et al., 015) by using the mean of the baseline period established in the previous step.
9) Extracting trial by trial mean amplitudes at 4 *a priori* defined time-windows: 50-150ms, 150-250ms, 250-500ms, 500-750ms; these time-windows cover the time-windows proposed by the multistep model (Kotz and Paulmann, 2011), plus our proposed fourth time-window. ERP and scalp plots of processed data were produced using the MNE package.

### Data Analysis

Figure 3 shows the model used for EEG data. The diagram is based on Kruschke’s (2015) and Martin’s (2018) model specification and presentation, and their guidelines on robust regression. The model was sampled using Markov Chain Monte Carlo (MCMC) No U-turn Sampling (NUTS) as provided by PyMC3 (Salvatier et al., 2016). All models were sampled with two chains of 2000 tuning steps and 2000 samples, and initialised using automatic differentiation variational inference (ADVI) as provided by PyMC3. Plots of results were produced using Arviz (Kumar et al., 2019) and Matplotlib (Hunter, 2007). Results were assessed using a region of practical equivalence (ROPE) method (Kruschke, 2015; Martin, 2018), where high posterior density (HDI) intervals were considered as presenting a considerable difference when far away from ROPEs defined as 1SD to 2SDs around zero.

**Figure 3.**
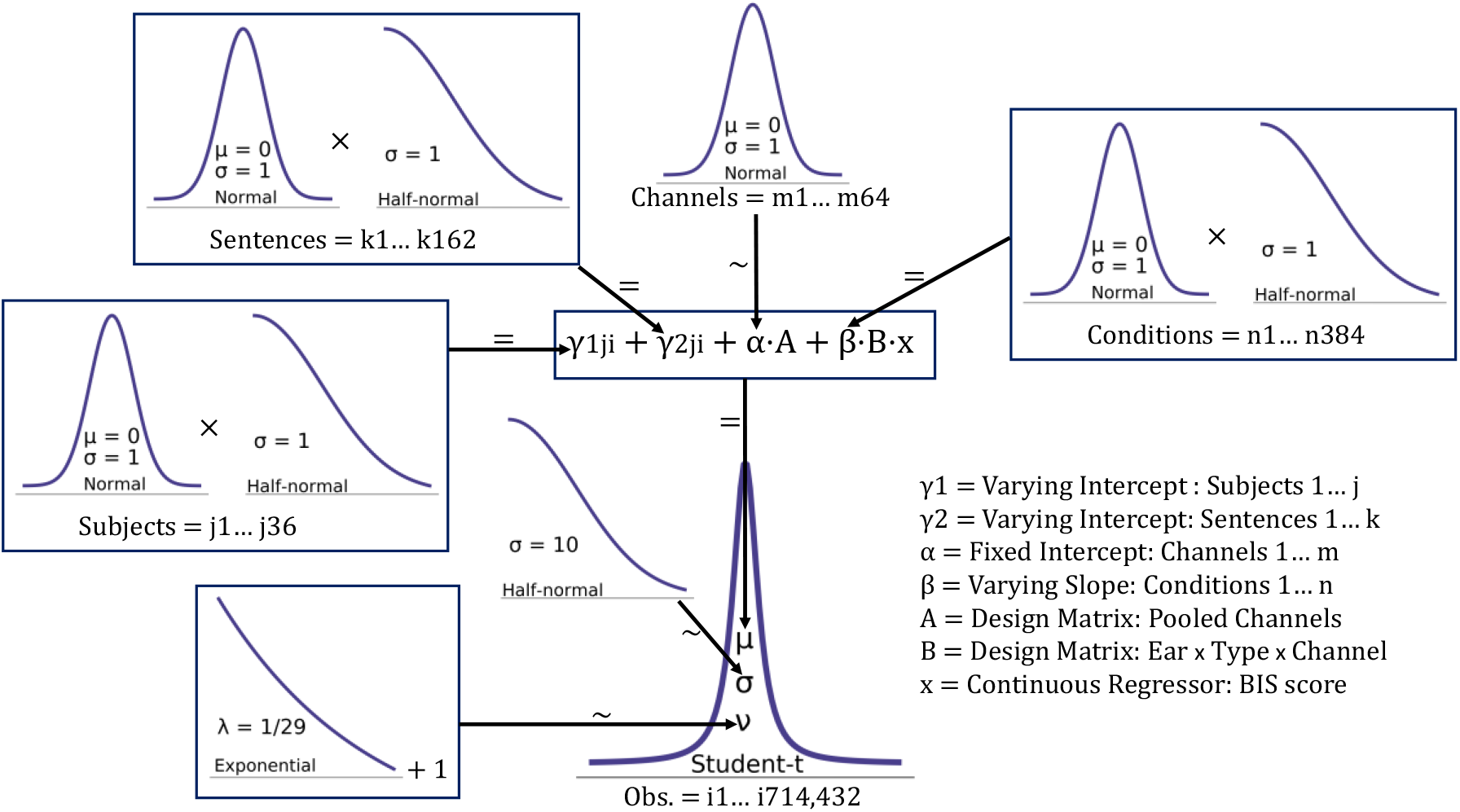
The diagram shows the prior distributions used for robust regression on amplitude. Arrows indicate the relationship between a parameter and priors or hyperpriors, where tilde (∼) indicates a stochastic relationship and equal (=) indicates a deterministic relationship. Observations (Obs.) in the likelihood distribution are equivalent to mean amplitude at each single trial. Channels: 64 electrodes. Ear: 2 ears (left and right). Type: 3 sentence-type (Congruent, Prosody, Semantic).

The core idea of the model is to un-pool data from the individual level of subjects and items, so we can focus on group level slopes at each single interaction point, trial-by-trial. Given this, pooling at the electrode (Channel in Figure 3) level through a normal non-varying prior helped us to define an offset for the other reparametrized priors (e.g. McElreath, 2020), which do not contain individual location parameters. In this way we could obtain intercepts from each electrode and improve sampling and convergence in a substantial manner. To be consistent with previous research, we did not include FFFS rating in the analyses, as our main focus is on trait anxiety and not on trait fear. Fear measures (FFFS) were collected as they could be required for comparison in future analyses. See our OSF repository for details on behavioural models’ structures (https://osf.io/n5b6h/).

In short, the model (Figure 3) comprises one non-varying intercept Channel (64 electrodes matrix); two varying intercept Subjects (36 participants) and Sentences (162 stimuli); and one varying slope (matrix: 2 ears by 3 sentence-types by 64 electrodes), multiplied by the BIS (x in Figure 3) continuous variable. All varying parameters are re-parametrised by using a scale (half-normal) by the main parameter (normal distribution). The likelihood is a Student-t distribution (see parameters in Figure 3), ideal for mean amplitude data, which tends to cluster around zero but leave long negative and positive tails; in this way possible outliers are included into the model instead of being discarded (Kruschke, 2015; Martin, 2018). The present model will be able to tell whether BIS varies differently given ear, sentence-type and electrodes parameters. In this way, we will obtain a posterior distribution with a respective HDI indicating the slope of BIS, namely how much amplitude increase/decreases as a function of BIS per electrode at each ear (left or right) per type (Congruent, Prosody, Semantic). This model is implemented separately at each time-window, thus allowing to observe whether BIS has an effect at early (50-150ms), mid-early (150-250ms), mid-late (250-500ms) or late (500-750ms) time-windows according to the present hypotheses. This implies that when there is an important increase/decrease in amplitude at a given condition (e.g. Congruent at Left Ear at Cz Channel), HDIs of BIS slopes should fall away from zero and ROPE and the specific amplitude change at a given electrode could be predicted simply by obtaining the product of any BIS score and the estimated posterior distribution of the slope.

## Results

### Behavioural Results

All accuracy models sampled well, showing excellent convergence (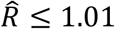, ESS > 900, BFMIs > 0.6); full summaries and plots can be found in the present study’s Open Science Framework (OSF) repository (https://osf.io/n5b6h/). Accuracy results indicate very high overall accuracy (over 90%) and no change as a function of ear, sentence type, nor BIS. Table 4 and Table 5 summarise accuracy results from direct- and indirect-threat conditions.

**Table 4.**
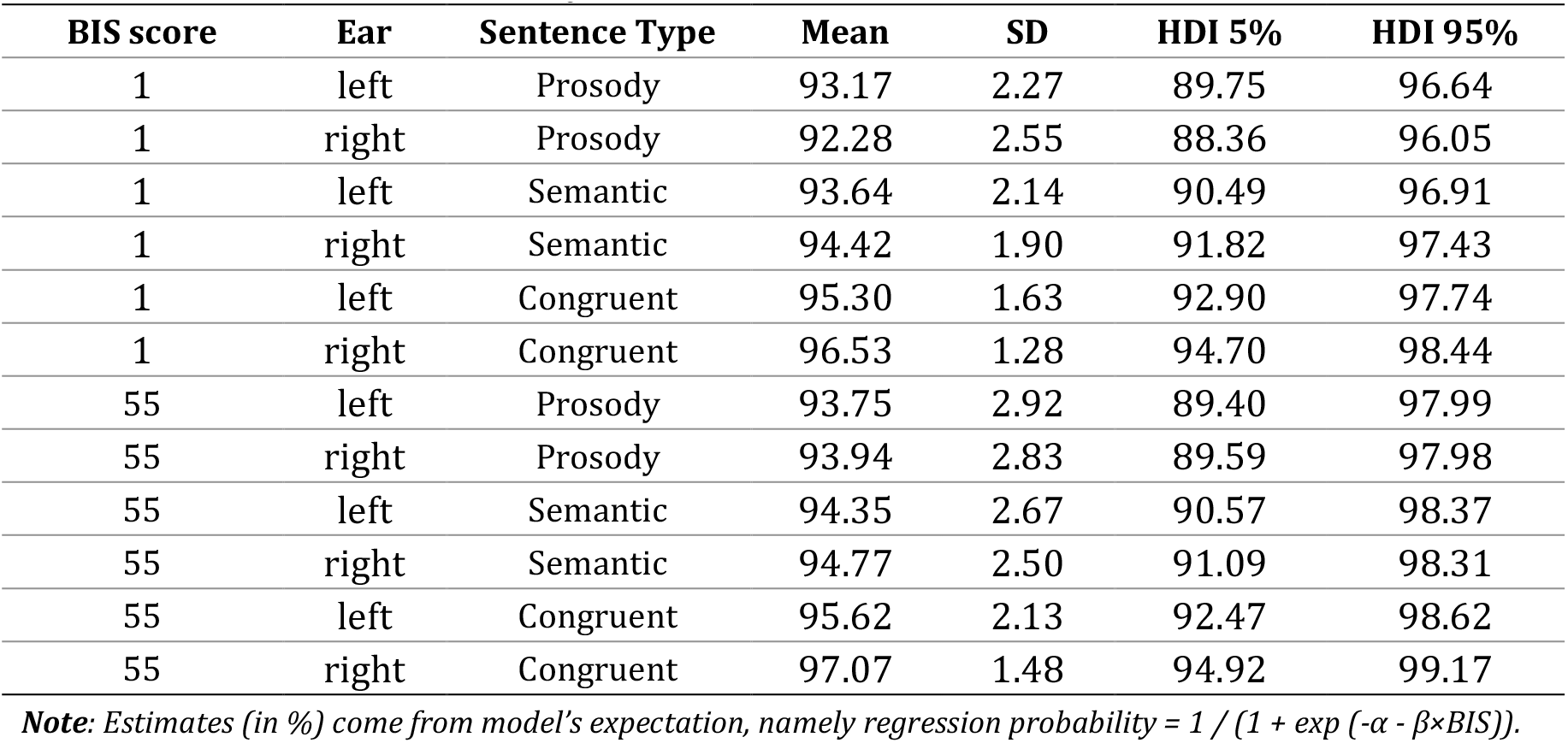
Direct-threat Accuracy Estimates

| BIS score | Ear | Sentence Type | Mean | SD | HDI 5% | HDI 95% |
| --- | --- | --- | --- | --- | --- | --- |
| 1 | left | Prosody | 93.17 | 2.27 | 89.75 | 96.64 |
| 1 | right | Prosody | 92.28 | 2.55 | 88.36 | 96.05 |
| 1 | left | Semantic | 93.64 | 2.14 | 90.49 | 96.91 |
| 1 | right | Semantic | 94.42 | 1.90 | 91.82 | 97.43 |
| 1 | left | Congruent | 95.30 | 1.63 | 92.90 | 97.74 |
| 1 | right | Congruent | 96.53 | 1.28 | 94.70 | 98.44 |
| 55 | left | Prosody | 93.75 | 2.92 | 89.40 | 97.99 |
| 55 | right | Prosody | 93.94 | 2.83 | 89.59 | 97.98 |
| 55 | left | Semantic | 94.35 | 2.67 | 90.57 | 98.37 |
| 55 | right | Semantic | 94.77 | 2.50 | 91.09 | 98.31 |
| 55 | left | Congruent | 95.62 | 2.13 | 92.47 | 98.62 |
| 55 | right | Congruent | 97.07 | 1.48 | 94.92 | 99.17 |

*Note:* Estimates (in %) come from model's expectation, namely regression probability = $1 / (1 + \exp(-\alpha - \beta \times \text{BIS}))$ .

**Table 5.** Indirect-threat Accuracy Estimates

| BIS score | Ear | Sentence Type | Mean | SD | HDI 5% | HDI 95% |
| --- | --- | --- | --- | --- | --- | --- |
| 1 | left | Prosody | 90.44 | 2.73 | 86.12 | 94.61 |
| 1 | right | Prosody | 89.67 | 2.96 | 85.07 | 94.31 |
| 1 | left | Semantic | 90.81 | 2.61 | 86.79 | 94.85 |
| 1 | right | Semantic | 90.96 | 2.63 | 86.68 | 94.86 |
| 1 | left | Congruent | 92.95 | 2.17 | 89.72 | 96.40 |
| 1 | right | Congruent | 92.14 | 2.30 | 88.79 | 95.87 |
| 55 | left | Prosody | 93.96 | 2.59 | 90.18 | 97.90 |
| 55 | right | Prosody | 93.41 | 2.81 | 89.24 | 97.57 |
| 55 | left | Semantic | 92.79 | 3.01 | 88.39 | 97.40 |
| 55 | right | Semantic | 91.92 | 3.39 | 86.85 | 97.07 |
| 55 | left | Congruent | 95.95 | 1.83 | 93.26 | 98.69 |
| 55 | right | Congruent | 94.89 | 2.22 | 91.71 | 98.30 |

*Note:* Estimates (in %) come from model's expectation, namely regression probability = $1 / (1 + \exp(-\alpha - \beta \times \text{BIS}))$ .

All reaction time (RT) models sampled well, showing relatively good convergence (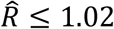, ESS > 300, BFMIs > 0.6); all summaries and plots can be found in present study’s Open Science Framework (OSF) repository (https://osf.io/n5b6h/). Direct-threat task estimates show a small but very uncertain (some HDI overlap) increase as a function of BIS (between 18ms to 52ms across conditions), suggesting no relevant effect. Results do not show relevant effects of ear either. However, Congruent RTs are between 96ms and 143ms lower than other sentence types. These differences do not show a clear or consistent pattern across ear or BIS score, indicating a general speed-up for Congruent sentences (with increased certainty). Indirect-threat estimates indicate a similar pattern, but the increase is slightly higher as a function of BIS for non-Congruent sentences (between 71ms and 103ms), but uncertainty is still relatively high. Again, responses to Congruent were faster than to other sentence types (differences between 97ms and 167ms). No clear differences across ear can be observed. See Table 6 and Table 7 for summaries of direct-threat and indirect-threat tasks respectively.

**Table 6.**
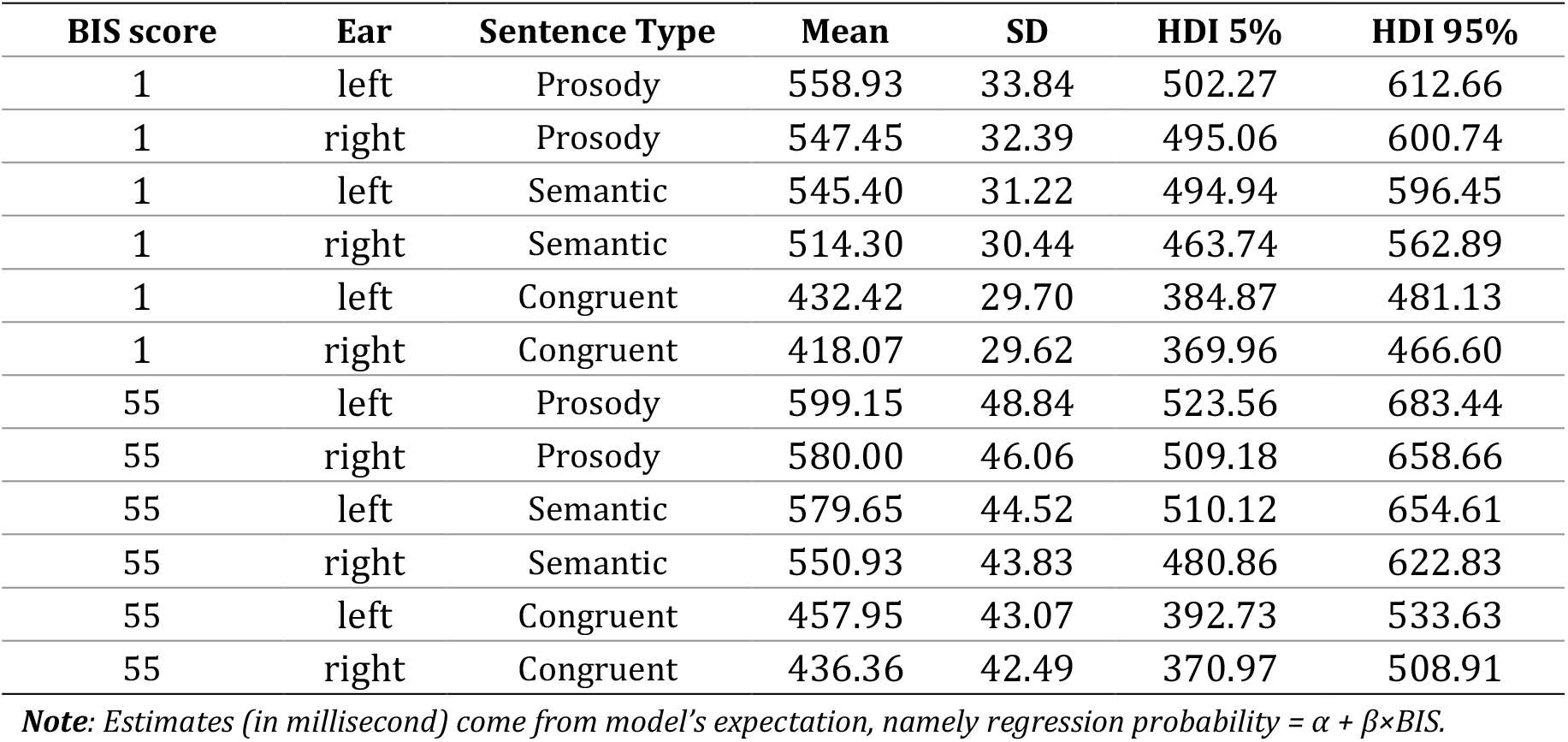
Direct-threat Reaction Times Estimates

| BIS score | Ear | Sentence Type | Mean | SD | HDI 5% | HDI 95% |
| --- | --- | --- | --- | --- | --- | --- |
| 1 | left | Prosody | 558.93 | 33.84 | 502.27 | 612.66 |
| 1 | right | Prosody | 547.45 | 32.39 | 495.06 | 600.74 |
| 1 | left | Semantic | 545.40 | 31.22 | 494.94 | 596.45 |
| 1 | right | Semantic | 514.30 | 30.44 | 463.74 | 562.89 |
| 1 | left | Congruent | 432.42 | 29.70 | 384.87 | 481.13 |
| 1 | right | Congruent | 418.07 | 29.62 | 369.96 | 466.60 |
| 55 | left | Prosody | 599.15 | 48.84 | 523.56 | 683.44 |
| 55 | right | Prosody | 580.00 | 46.06 | 509.18 | 658.66 |
| 55 | left | Semantic | 579.65 | 44.52 | 510.12 | 654.61 |
| 55 | right | Semantic | 550.93 | 43.83 | 480.86 | 622.83 |
| 55 | left | Congruent | 457.95 | 43.07 | 392.73 | 533.63 |
| 55 | right | Congruent | 436.36 | 42.49 | 370.97 | 508.91 |

**Note:** Estimates (in millisecond) come from model's expectation, namely regression probability = $\alpha + \beta \times \text{BIS}$ .

**Table 7.**
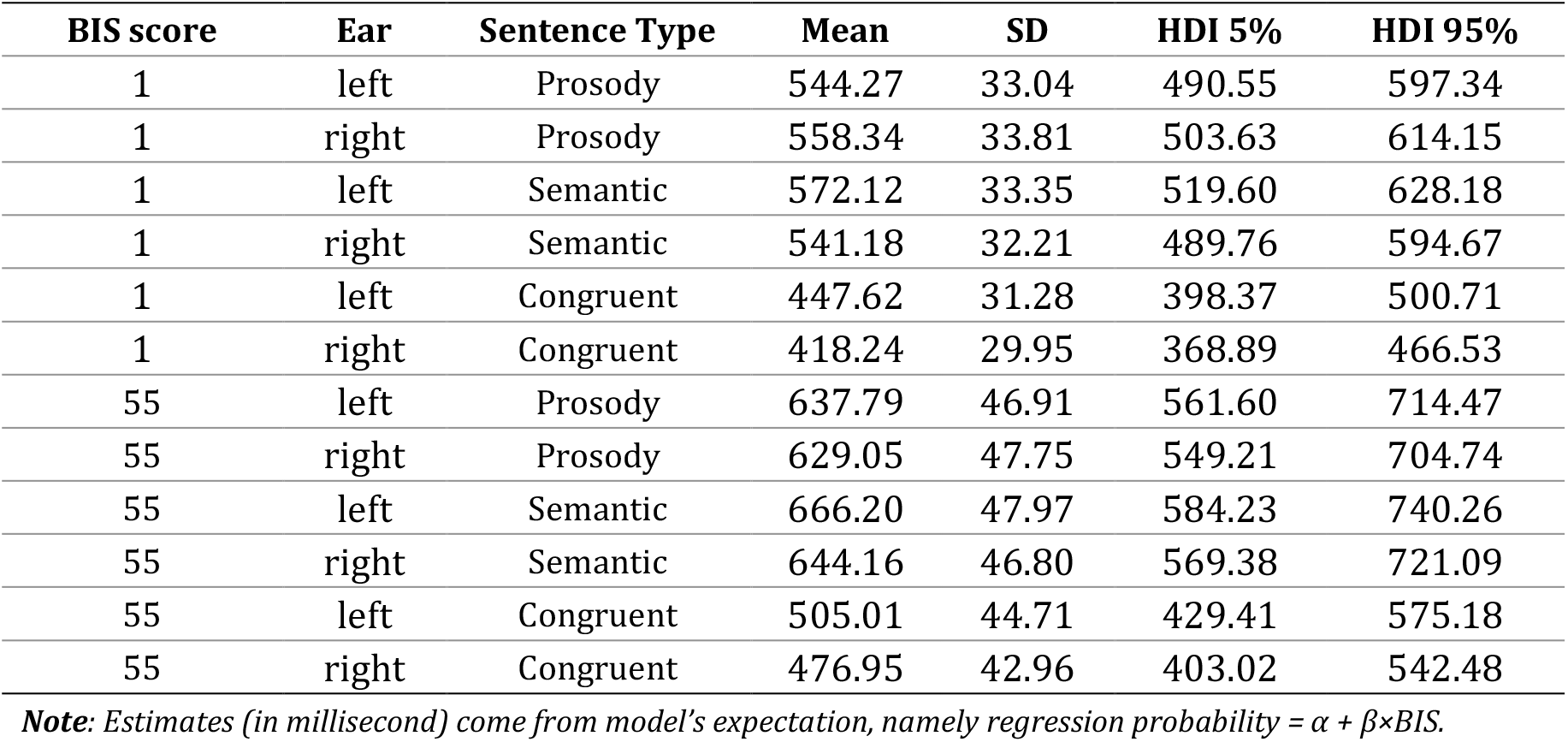
Indirect-threat Reaction Times Estimates

### ERP Results

Although global field power (GFP) at EEG channels does not drift and EEG polarity is aligned (see: Luck, 2014), visual inspection of grand average ERP waves for direct- and indirect-threat conditions also indicate a potentially artifactual drift at EOG activity (Figure 4). This also shows that ERP waves are similar across tasks (direct- and indirect-threat), indicating a systematic amplitude increase after 500ms. Figure 5 and Figure 6 show mean amplitude peaks selected by maximum local GFP, which indicates a spatial standard deviation quantifying electrical activity from all electrodes at specific time-points (Skrandies, 1990), reliably revealing peaks of greater amplitude. We present these only as an initial descriptive inspection of amplitude scalp distribution by BIS terciles (divided into terciles for illustration purposes only).

**Figure 4.**
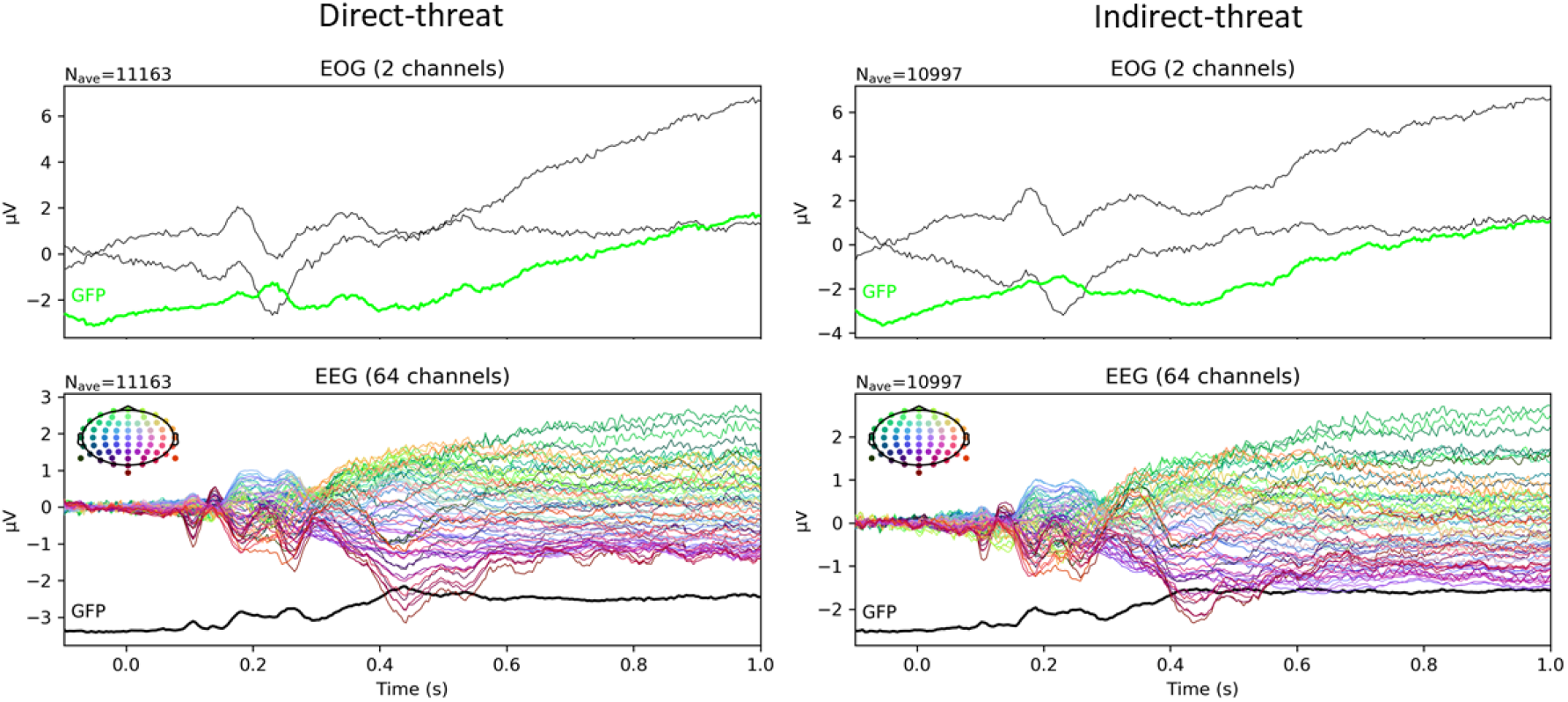
General ERP grand averages summary. Left panels show EOG and EEG waves from the direct-threat task, and right panels from the indirect-threat task. Bottom panels grand averages across all 64 electrodes. EOG: two electrooculogram channels. *N*ave: total number of trials averaged. GFP: global field power (local maximum).

**Figure 5.**
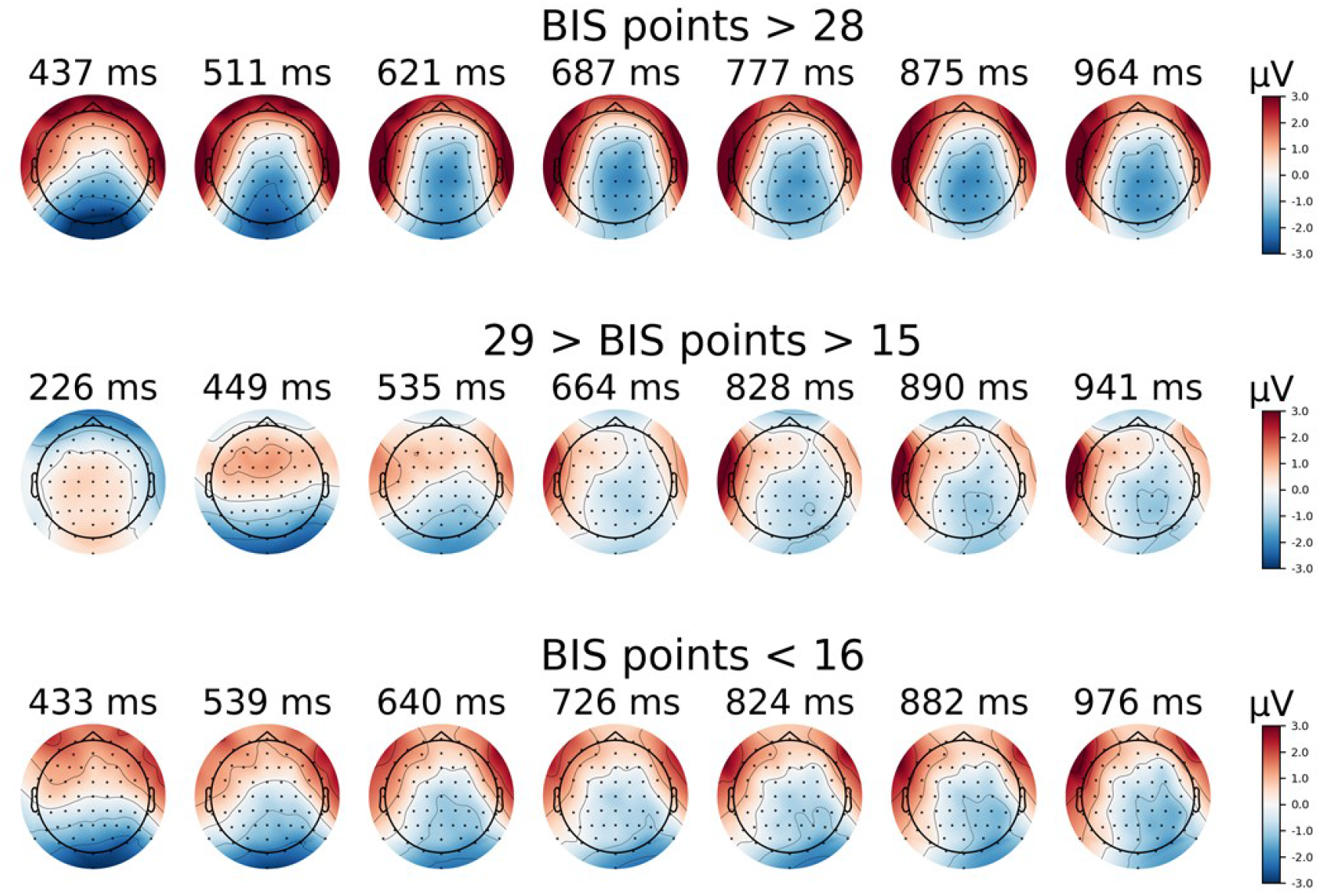
Direct-threat topographies. Images show raw mean amplitude across epoch by BIS tercile (for illustration purposes only). Top: upper BIS score tercile. Middle: mid BIS score tercile. Bottom: lower BIS score tercile. Time-points selected as the local maxim Global Field Power (GFP).

**Figure 6.**
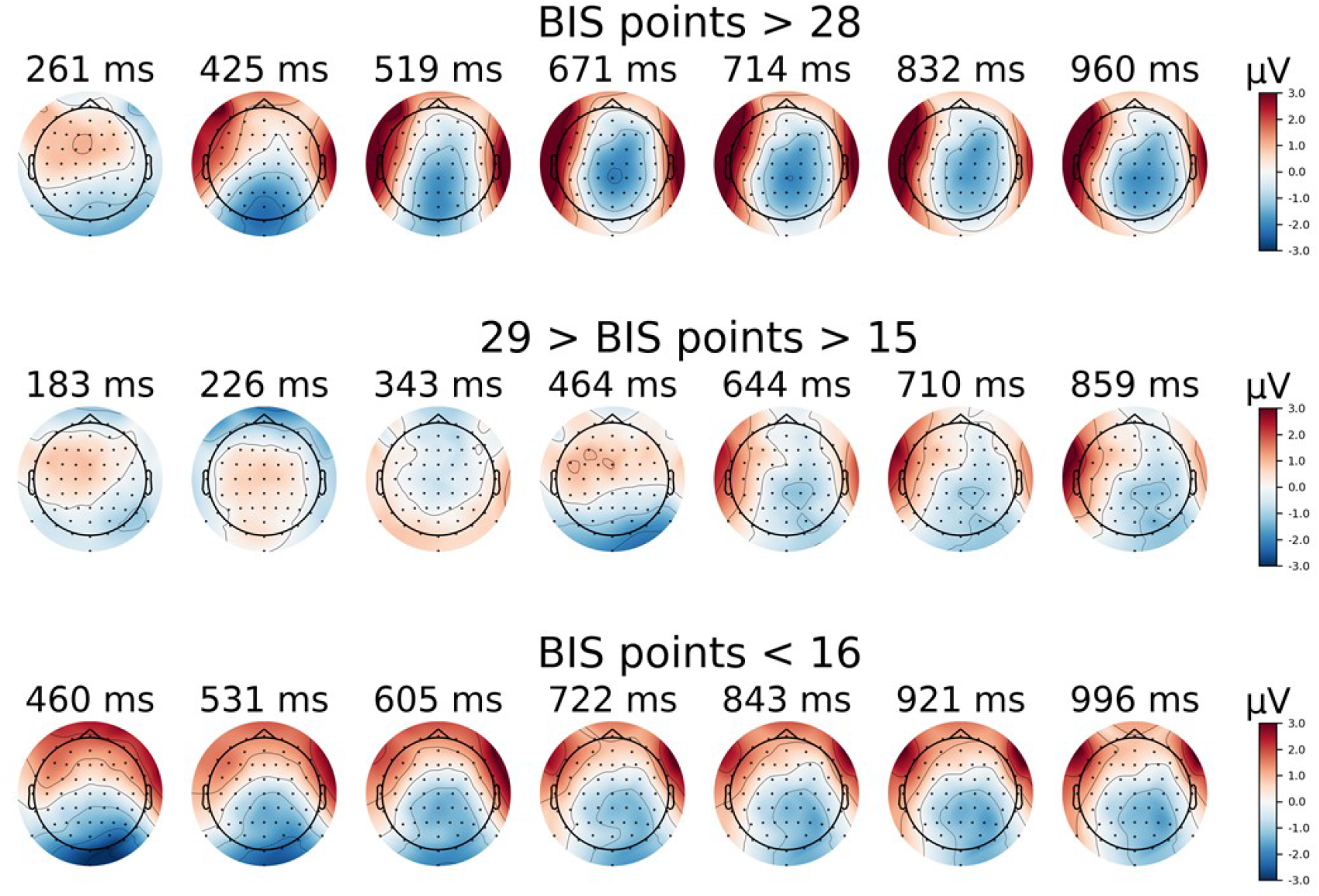
Indirect-threat topographies. Images show raw mean amplitude across epoch by BIS tercile (for illustration purposes only). Top: upper BIS score tercile. Middle: mid BIS score tercile. Bottom: lower BIS score tercile. Time-points selected as the local maxim Global Field Power (GFP).

Hierarchical models indicate that excepting window4 (500-750ms), no other time-window showed relevant effects as a function of sentence-type, ear or BIS (HDIs overlapping zero or widely inside ROPE). All models sampled well, showing excellent convergence (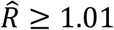, ESS > 300, BFMIs > 0.6). Tables 8 and 9 summarise estimates that represent posteriors from BIS slope of the regression. Table 8 summarises direct-threat results from window4, showing some example electrodes at bilateral positions: TP7 (left) and TP8 (right). For TP7 amplitude increases from lowest BIS (1 point) to highest BIS (55 points) ∼2μV for Congruent at both ears, ∼2.6-2.8μV for Prosody at both ears, and ∼3.6μV for Semantic at both ears. For TP8 amplitude increases from lowest BIS (1 point) to highest BIS (55 points) ∼2.5μV for Congruent at both ears, ∼1.6-1.9μV for Prosody at both ears, and ∼2.2μV for Semantic at both ears. Table 9 summarises the same electrodes from the indirect-threat task (window4), indicating very similar results. Figures 7 and 8 help to visualise these differences at electrode TP7. Similar increases were registered at for electrodes such as T7, T8, P7, P8, P9 and P10. Supplementary materials contain plots of estimates of all previously mentioned relevant electrodes (Supplement 2).

**Table 8.**
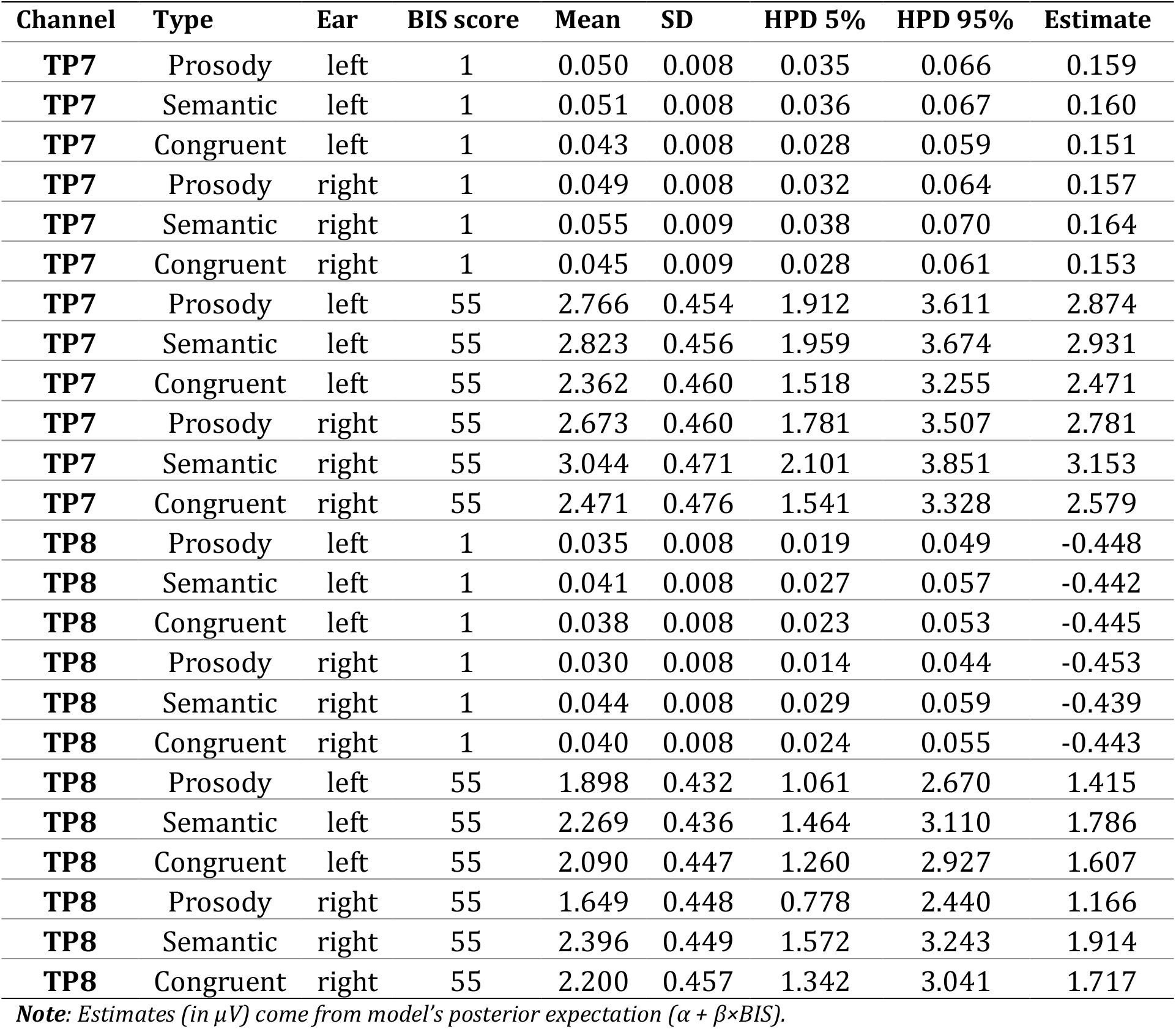
Direct-threat Estimates of Selected Electrodes

**Table 9.** Indirect-threat Estimates of Selected Electrodes

| Channel | Type | Ear | BIS score | Mean | SD | HPD 5% | HPD 95% | Estimate |
| --- | --- | --- | --- | --- | --- | --- | --- | --- |
| TP7 | Prosody | left | 1 | 0.047 | 0.009 | 0.030 | 0.063 | 0.124 |
| TP7 | Semantic | left | 1 | 0.068 | 0.008 | 0.053 | 0.084 | 0.145 |
| TP7 | Congruent | left | 1 | 0.035 | 0.008 | 0.020 | 0.051 | 0.113 |
| TP7 | Prosody | right | 1 | 0.052 | 0.008 | 0.037 | 0.069 | 0.129 |
| TP7 | Semantic | right | 1 | 0.066 | 0.009 | 0.050 | 0.082 | 0.143 |
| TP7 | Congruent | right | 1 | 0.039 | 0.008 | 0.024 | 0.054 | 0.116 |
| TP7 | Prosody | left | 55 | 2.600 | 0.485 | 1.659 | 3.483 | 2.677 |
| TP7 | Semantic | left | 55 | 3.732 | 0.462 | 2.888 | 4.622 | 3.809 |
| TP7 | Congruent | left | 55 | 1.944 | 0.467 | 1.079 | 2.816 | 2.021 |
| TP7 | Prosody | right | 55 | 2.867 | 0.457 | 2.058 | 3.774 | 2.944 |
| TP7 | Semantic | right | 55 | 3.621 | 0.469 | 2.739 | 4.516 | 3.699 |
| TP7 | Congruent | right | 55 | 2.125 | 0.448 | 1.312 | 2.971 | 2.202 |
| TP8 | Prosody | left | 1 | 0.030 | 0.009 | 0.015 | 0.047 | -0.363 |
| TP8 | Semantic | left | 1 | 0.041 | 0.009 | 0.025 | 0.057 | -0.352 |
| TP8 | Congruent | left | 1 | 0.044 | 0.009 | 0.027 | 0.060 | -0.350 |
| TP8 | Prosody | right | 1 | 0.036 | 0.009 | 0.020 | 0.051 | -0.358 |
| TP8 | Semantic | right | 1 | 0.040 | 0.008 | 0.025 | 0.056 | -0.353 |
| TP8 | Congruent | right | 1 | 0.046 | 0.008 | 0.031 | 0.062 | -0.347 |
| TP8 | Prosody | left | 55 | 1.675 | 0.473 | 0.801 | 2.559 | 1.281 |
| TP8 | Semantic | left | 55 | 2.268 | 0.469 | 1.386 | 3.136 | 1.875 |
| TP8 | Congruent | left | 55 | 2.396 | 0.483 | 1.466 | 3.277 | 2.003 |
| TP8 | Prosody | right | 55 | 1.953 | 0.468 | 1.080 | 2.811 | 1.559 |
| TP8 | Semantic | right | 55 | 2.199 | 0.465 | 1.371 | 3.090 | 1.806 |
| TP8 | Congruent | right | 55 | 2.549 | 0.458 | 1.692 | 3.430 | 2.156 |

*Note: Estimates (in $\mu V$ ) come from model's posterior expectation ( $\alpha + \beta \times BIS$ ).*

**Figure 7.**
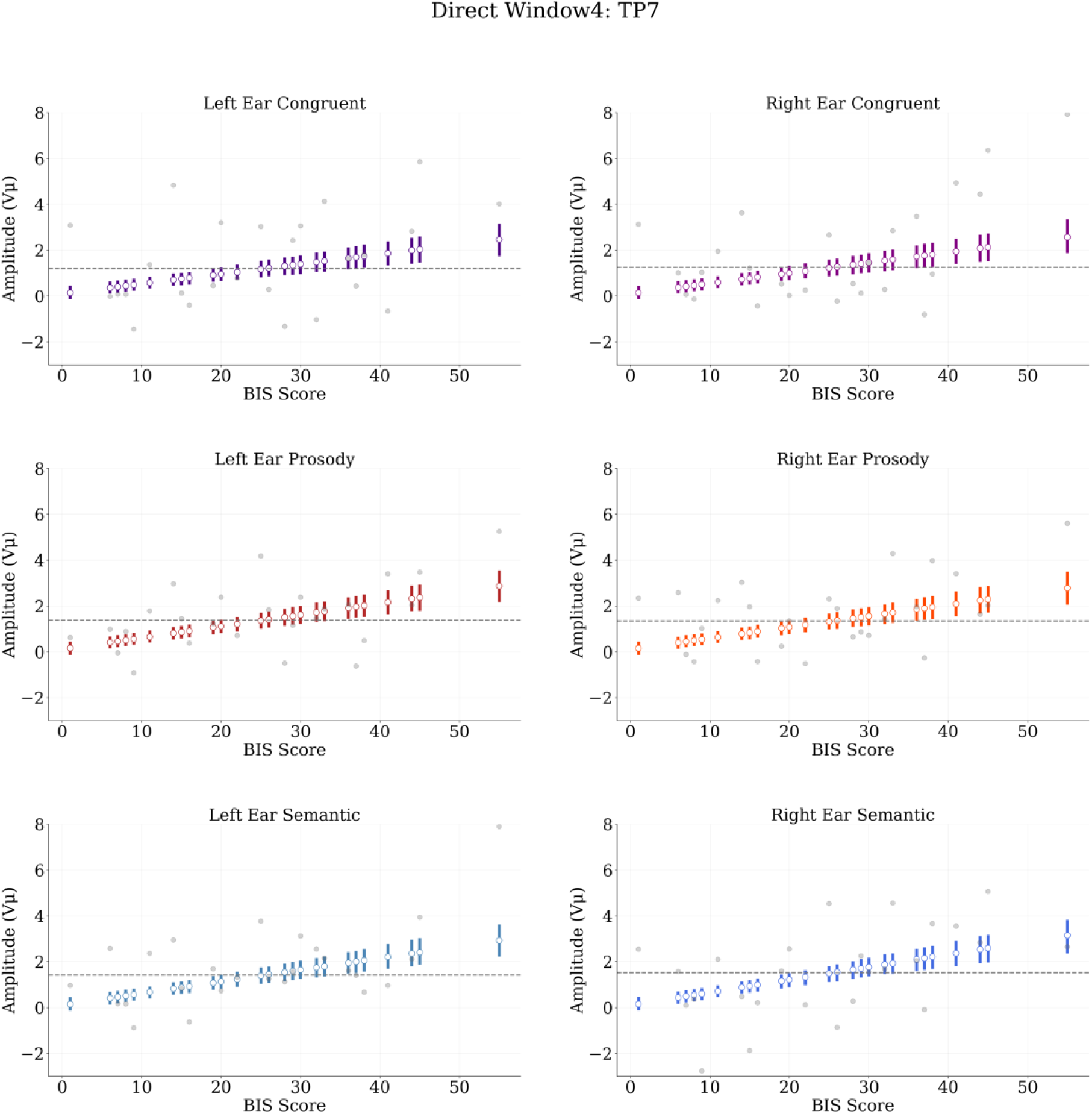
Direct-threat regressions at electrode TP7, Window4 (500-750ms). White circles represent posterior means by BIS score. Bars represent highest density intervals (HDIs). Grey dots are mean amplitudes by BIS score. Dashed grey line indicates posterior median BIS.

**Figure 8.**
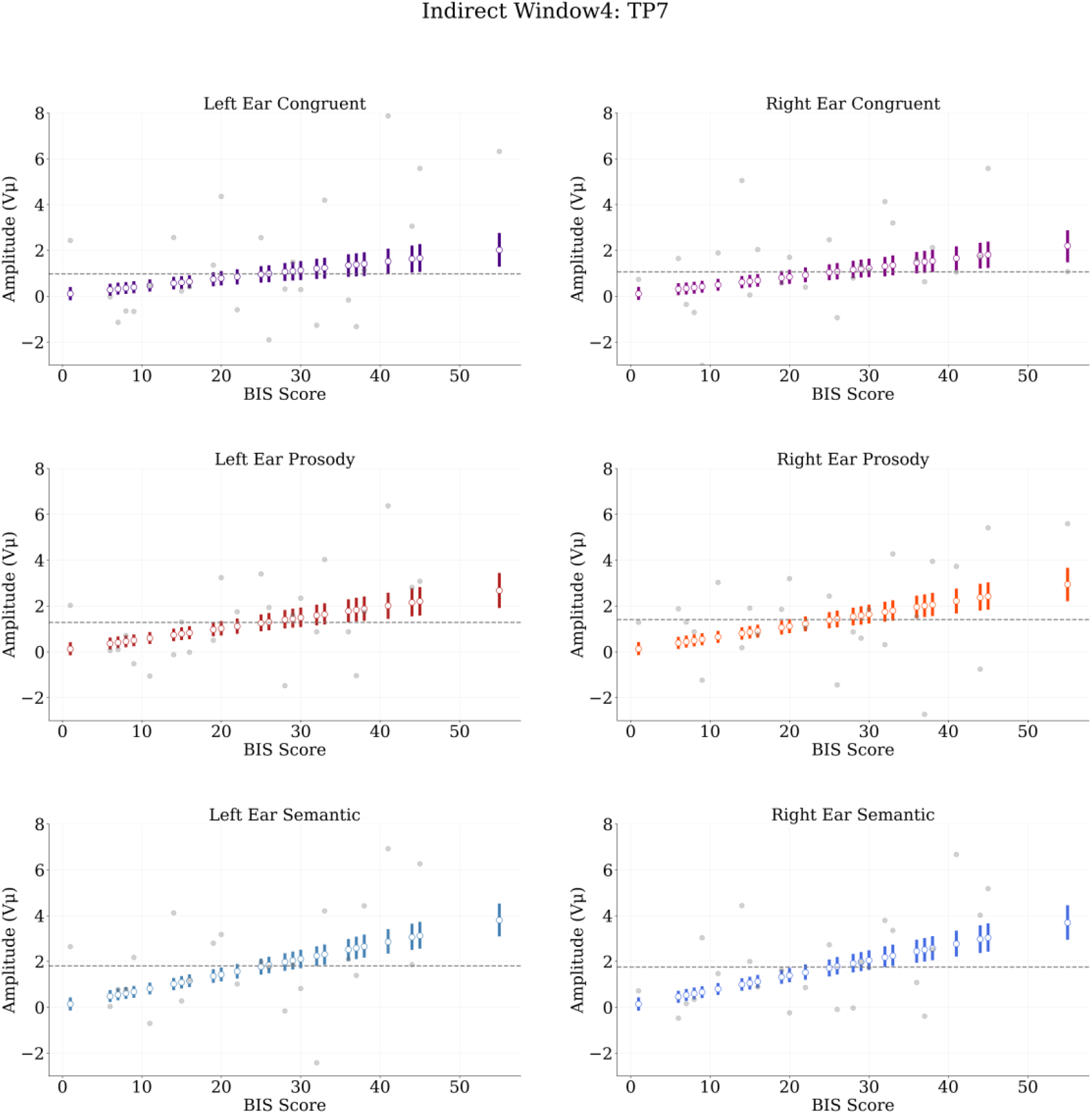
Indirect-threat regressions at electrode TP7, Window4 (500-750ms). White circles represent posterior means by BIS score. Bars represent highest density intervals (HDIs). Grey dots are mean amplitudes by BIS score. Dashed grey line indicates posterior median BIS.

Note that in both tables highest BIS scores HDIs are far from overlapping with the lowest BIS scores (clearly seen in Figures 7 and 8). This implies that the magnitude of effects shows good certainty, indicating a sustained increase in amplitude from the lowest BIS level (1 point) to the highest BIS level (55 points), with good precision (HDI widths do not surpass ROPE width). This indicates that the greatest amplitude increases in window4 (500-750ms) occur at bilateral temporo-parietal electrodes, but with a clearer LPC pattern at temporal sites, in particular in the LH (e.g. TP7) as a function of BIS. Figures 9 and 10 show average amplitude waves from example electrodes from the direct- and indirect-threat tasks respectively. Although earlier time-windows suggest some differences, these are subject to high variation, thus estimates barely change as a function of BIS (see supplement 3). Window3 shows a small BIS-related increase for non-congruent conditions (see supplement 3). These are not directly reported here, as they could be related to early LPC onset rather than to particular Window3 activity (i.e. low BIS inducing N400 effects).

**Figure 9.**
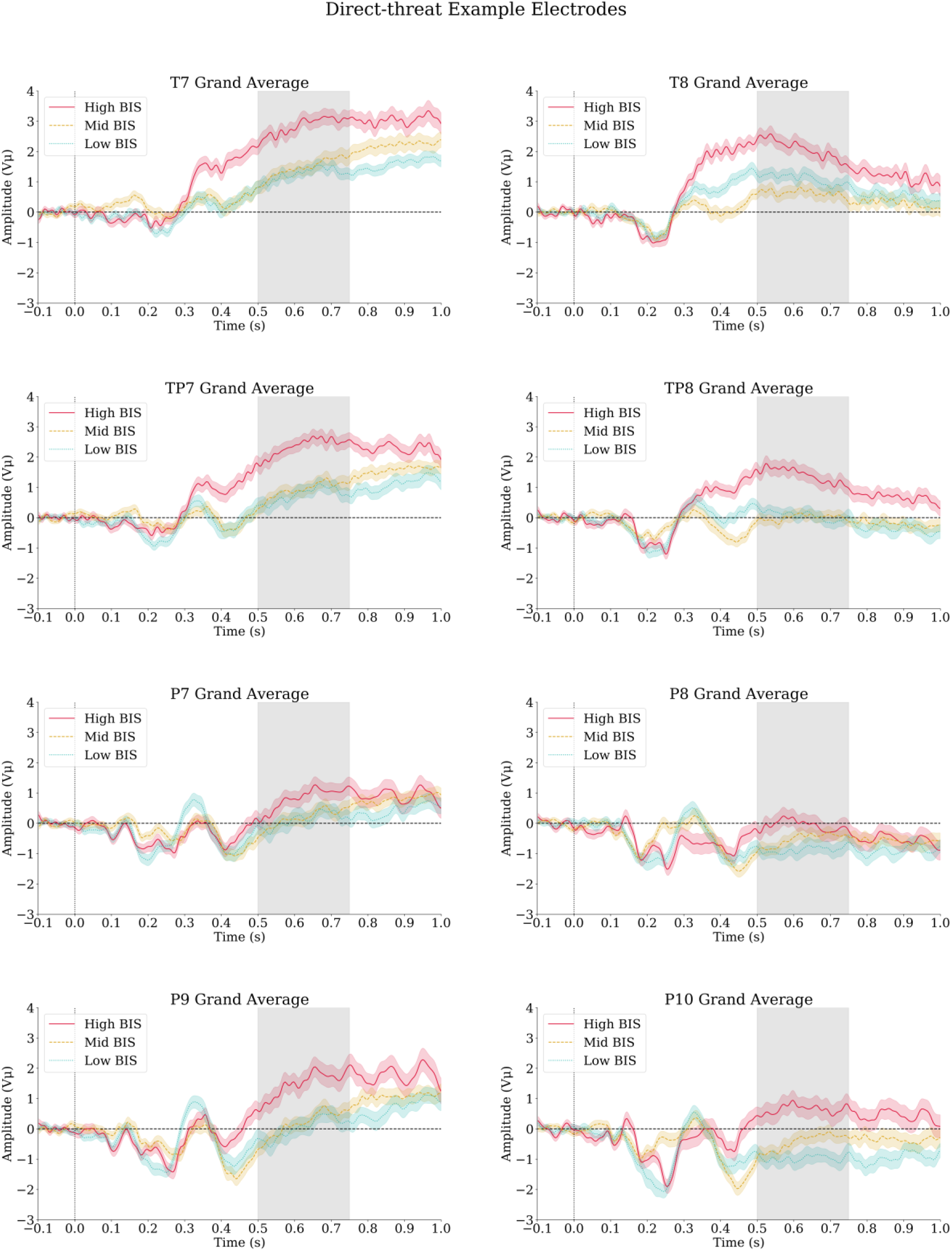
Direct-threat example electrodes. Plots show grand averages (−100-1000ms epoch), low-pass filtered at 40Hz, collapsed across ear and sentence type, and divided into BIS terciles for display purposes only. High, red solid line: BIS score > 28. Mid, yellow dashed line: 29 > BIS score > 15. Low, blue dotted line: BIS score < 16. Shadows indicate standard error of the mean (SEM). Black dotted vertical line indicates sentence onset, period before the dotted line is the 100ms baseline. Grey shaded region indicates time window4.

**Figure 10.**
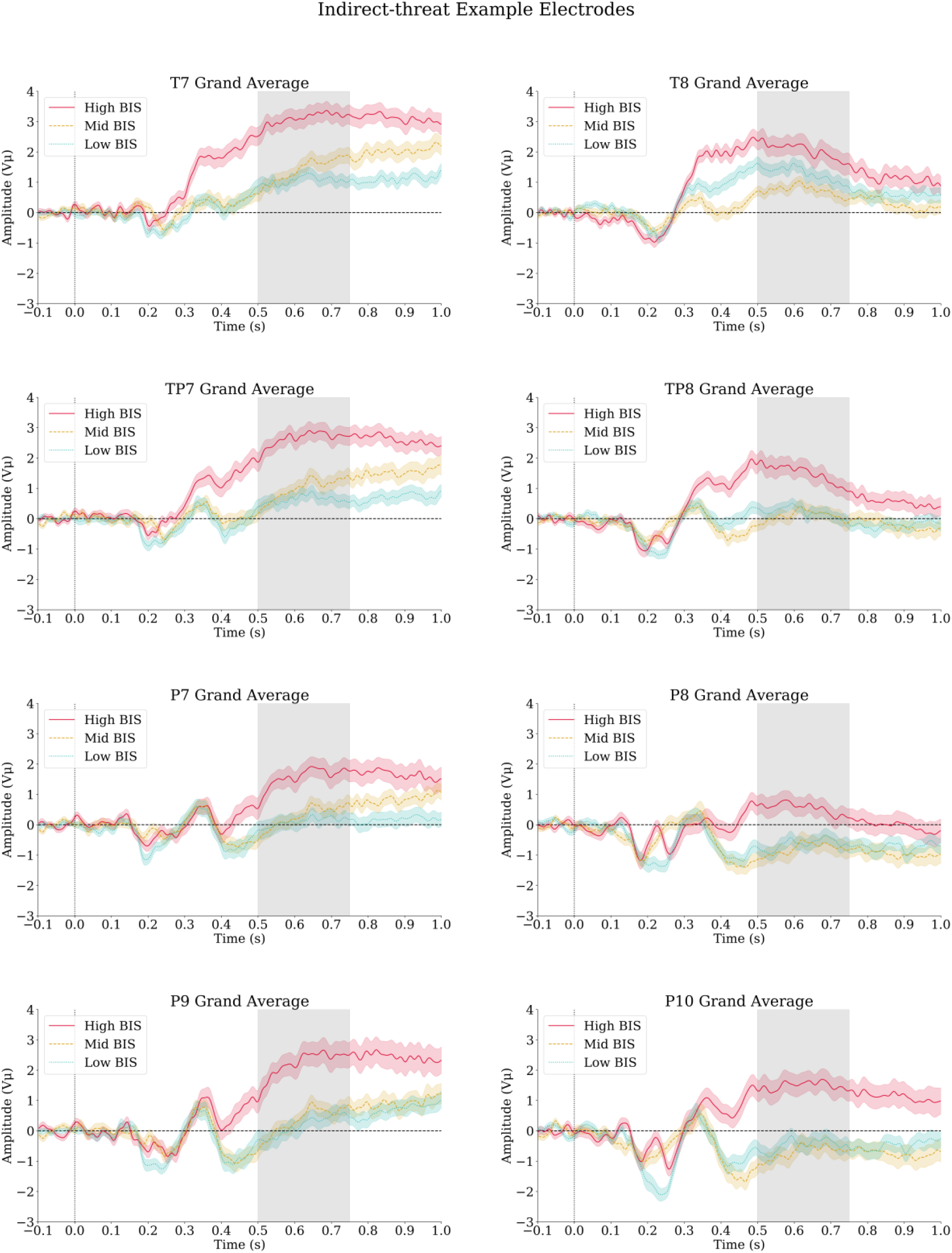
Indirect-threat example electrodes. Plots show grand averages (−100-1000ms epoch), low-pass filtered at 40Hz, collapsed across ear and sentence type, and divided into BIS terciles for display purposes only. High, red solid line: BIS score > 28. Mid, yellow dashed line: 29 > BIS score > 15. Low, blue dotted line: BIS score < 16. Shadows indicate standard error of the mean (SEM). Black dotted vertical line indicates sentence onset, period before the dotted line is the 100ms baseline. Grey shaded region indicates time window4.

In short, both direct- and indirect-threat tasks show similar patterns of activity, consistent with models’ estimates. Window4 results are reported in more detail as main results, as they express the clearer effects. These suggest that irrespective of task (answer to threat or neutral), ear presentation (left or right) or sentence-type (Congruent, Prosody or Semantic), between 500ms and 750ms amplitude increases, on average, as a function of BIS score at bilateral temporo-parietal electrodes, but showing slightly higher amplitude at LH electrodes.

## Discussion

Present behavioural results indicate that threatening stimuli in both direct- and indirect-threat tasks induce little to no effect on reaction times (RTs) or accuracy as a function of BIS (trait anxiety), no effects for ear, and a general speed up for the Congruent condition. This implies that for delayed response tasks, effects of trait anxiety (worry or BIS) are of a small magnitude or negligible. Models’ estimates indicate that in the 500-750ms time-window (window4) amplitude increases, on average, between 1.6μV and 3.6μV in temporo-parietal electrodes as a function of BIS (slightly stronger increases at the left hemisphere). This can be interpreted as a BIS-related late positive complex (LPC) increase. This is consistent with part of our hypotheses, indicating late phase effects of trait anxiety on threatening language processing during orientative (deliberation) phases. However, no clear earlier effects can be observed as a function of BIS, which goes contrary to our hypotheses predicting the influence of trait anxiety at all time-windows from sentences’ onsets. In the Mid-phase window3 (250-500ms) also shows increases for non-congruent conditions (i.e. Prosody and Semantic). We have reported these in supplement 3, as these increases could be attributed to earlier LPC onset for some electrodes/conditions and do not seem to be associated with a particularly strong negative deflection (e.g. N400).

Nevertheless, effects on Window3 could still provide evidence that at lower BIS scores evaluation processes are different, and that extended evaluation has an earlier onset for higher BIS scores (i.e. before 500ms) and simply extends towards later stages. This would be contrary to our interpretation of LPC and LPC-like late phase activity representing a deliberation phase. Instead, it may fit into the conventional scheme of a triphasic process, such as the one proposed by the multistep model of emotional language (Kotz and Paulmann, 2011). This could explain why these effects are not so evident for the Congruent condition, simply implying that stimuli in this category are more easily recognisable and do not elicit N400-like effects. Namely, a more classic prosodic-semantic congruency effect (e.g. Schirmer and Kotz, 2003) cannot be discarded as an explanation. Having said this, the strong and clear LPC effects may require a complementary explanation.

Previous EEG studies have observed that worry is associated with a parietal late positive potential (LPP) increases, related to negative reappraisal of visual stimuli (Moser et al., 2014). This is understood to be associated with sustained attention inducing increased late-phase processing (Hajcak et al., 2010). This may correspond to delayed disengagement from threat (Cisler and Koster, 2010), which in the present paper we understand as over-engagement with threat during deliberation phases, occurring after 500ms (i.e. LPC time-window), in light of the association between higher BIS and anxiety, worry and rumination (McNaughton, 2011). However, studies using a cross-modal paradigm, priming words with emotional faces to observe implicit reappraisal have instead observed early-phase (e.g. N170) but not late-phase (i.e. LPP) effects (Liu et al., 2018), suggesting implicit rather than explicit engagement with threat when stimuli are short and/or fear-inducing. EEG studies focusing on social anxiety have shown that abusive words (short duration stimuli) induce effects on early- or mid-phase ERPs (i.e. P1, N400) which could be better related to over-attention to threat or threat evaluation issues (Wabnitz et al., 2015). Other studies have shown that angry vocalisations (prosody) are associated with early (i.e. P1) but not late (i.e. LPC) amplitude increases associated with anxiety (Pell et al., 2015). For anxious participants to over-engage with threat, longer stimuli that are also less directly threatening (i.e. not fear inducing or directly harming) stimuli may be required (Cisler and Koster, 2010).

This could imply that the present LPC, peaking ∼600ms after sentence onset, requires the deliberation time provided by long duration sentences. For the present experiment, sentences had a duration of ∼1646ms on average, which gives plenty of time (during sentence) for assessing and re-assessing stimuli before responses are executed after sentence’s offset. In other words, with sufficient time to over-engage with threatening stimuli in a decision task, effects were mainly driven by trait anxiety; namely higher BIS levels induced over-engagement with threat, resulting in increased processing during late-phase deliberation (after 500ms). This is consistent with LPC evidencing decision modulation in relation to evaluation through memory processes (Finningan, 2002; Yang et al., 2019), or LPP/LPC evidencing late-phase processing associated with anxious negative re-appraisal (Hajcak et al., 2010; Moser et al., 2014). In this line, trait anxiety has been strongly associated with patterns of repetitive thinking (McEvoy et al., 2010; McLaughlin et al., 2007), which is a mechanism for reappraisal and over-engagement. We propose that in the present experiment, threatening speech may have induced anxious participants to engage in verbal repetitive thinking, which can be associated with both memory-related evaluation and re-appraisal. This is theoretically sound with respect to BIS and anxiety models (Bar-Haim et al., 2007; Cisler and Koster, 2010; McNaughton, 2011; Robinson et al., 2019); although verbal repetitive thinking is not the only possible mechanism for explaining over-engagement with threat. It is still possible that a more general over-engagement, or delayed disengagement mechanism (Cisler and Koster, 2010) could have induced extended or delayed evaluation of emotional stimuli. This would fit a more classical interpretation from the multistep model perspective (Kotz and Paulmann, 2011), also suggesting that at those late evaluation phases prosody or semantics have no distinct or specific relevant relationships with anxiety.

Another important aspect of our results, contradicting our predictions, is the lack of clear laterality effects. Although some EEG studies show lateralization effects associated with emotional semantic and prosody variation (Kotz and Paulmann, 2007), recent research does not show much evidence for these effects (Chen et al., 2011; Paulmann et al., 2012). One proposed explanation is that concurrent multi-information-channel (prosody and semantic) information obscures laterality effects (Paulmann et al., 2012). In addition, the present LPC is not modulated by stimulus-type. This differs from previous research, which did not include anxiety measures, finding late positive amplitudes in association with prosody/semantic emotional variations and/or congruency effects (Astésano et al., 2004; Chen et al., 2011; Zhao et al., 2015). Given this, the present LPC, which shows a different distribution and different preceding ERPs from the aforementioned studies, could be understood as signalling something particular to anxiety rather than being directly associated with prosody or semantics. Even so, despite that amplitude differences were barely affected by ear presentation, we did observe a small lateralisation effect at LH-located electrodes. On average, these were around one microvolt higher in amplitude compared to their right hemisphere (RH) counterparts. Although some slight HDI overlap between LH and RH electrodes (e.g. TP7 and TP8) was observed at lower BIS scores, at higher BIS scores it was not so for most of the relevant electrodes. If this effect is not artefactual, it may suggest a more general late-stage lateralisation pattern, insensitive to dichotic effects, which may have been more evident at earlier time-windows given more controlled or shorter duration stimuli.

Although present research provides evidence of both behavioural and EEG effects (or lack thereof) of trait anxiety (BIS) on responses to threatening speech, we need to address some relevant limitations. Firstly, we do not have a direct neutral-speech control, as our only alternative task requiring participants to answer to neutral sentences (indirect-threat), usually intended to control for attentional differences (e.g. Peschard et al., 2016; Sander et al., 2005), still exposes participants to threatening speech. Secondly, even though we have applied an automated pre-processing and cleaning procedure to EEG data, rigorously based on previous literature, we acknowledge that there may still be a positive artefactual drift present in our epoched data. However, if the observed LPC component is not really an LPC (i.e. derived from brain activity), its association with BIS (trait anxiety) still needs to be explained. For instance, it may be the case that increased base muscle tension (e.g. jaw clenching) or rumination-related oro-facial muscle activity, characteristic of anxiety (Nalborczyk, 2017), induced artifacts on the ERP waves. However, muscular activity should induce higher frequency activity rather than lower frequency drifts, especially if related with temporalis muscles affecting temporal electrodes (Luck, 2014). Such artifacts, nonetheless, may have been detrimental for earlier activity (before 400-500ms), which may also explain the difficulty in observing earlier ERP effects. Despite this, excessive muscle tension could have indirectly induced eye artifacts at lower frequencies and amplitudes, which may have not been caught by ICA and other cleaning procedures. However, here we cannot provide direct evidence for this possible anxiety-related EOG activity, and we have not succeeded in finding an account of this possibility in previous literature.

Furthermore, there are some design limitations that should be acknowledged. Presently, the high variability of type and sentence position of threatening lexical items in Semantic and Congruent conditions may have led participants to primarily carry out threat recognition by focusing on threatening prosody, which always started at sentence onset. Thus, possible ear differences may have become very difficult to observe at early processing stages and/or quickly receded before later stages. In addition, this possibly strategic response of participants (focus on prosody) may have obscured sentence-type effects. This is associated to another limitation, namely that some sentences have threatening words at the end (last word) or very late within the sentence (i.e. after 600ms). These sentences conflict with our epoch decision: from sentence onset to 1000ms, as some of these threatening words fell outside the epochs’ offset. However, this decision was necessary to avoid movement contamination from reaction times, as many sentences were barely longer than one second. Furthermore, even when threatening words were placed at the end, evaluation or deliberation effects could still be present before sentence end as part of an orientation to response. This implies, nonetheless, another limitation. The presently observed effect of BIS on amplitude may be task-related instead of being associated with threatening speech.

Better controlled semantic stimuli could help to address these limitations, albeit at the cost of the semi-naturalness of stimuli. Also, it may be useful to employ methods that can directly induce response inhibition to differentiated stimuli (i.e. responses to threatening content or sound), rather than decision tasks as employed here. Previous research has indeed proposed that tasks such as go/no-go are better for understanding BIS processes, because they directly involve inhibiting responses as part of their design (e.g. McNaughton et al., 2013; Neo et al., 2011). Future research could implement such tasks by, for instance, including responses to stimulus type instead to threat in general. In addition, it would be relevant to implement tasks which can compare language stimuli with other type of stimuli and have a proper neutral-stimuli control; this could help to determine whether anxiety effects are specifically related to threatening speech or not. In other words, it would help determine whether the observed increasing LPC is threat-induced or task-induced.

In conclusion, present ERP analyses show a clear positive amplitude deflection in temporo-parietal electrodes peaking at around 600ms as a function of BIS scores. We interpreted this ERP as an LPC and suggest two possible interpretations. 1) This LPC is part of an extended evaluation process, which would be evidenced by lower BIS-related amplitude decreases in response to prosodic-semantic congruency, but higher BIS-related positive deflections (i.e. early onset LPC), which extend to later phases. 2) Present LPC is associated with trait anxiety affecting deliberation processes through verbal repetitive thinking. As BIS scores increased, the LPC became more positive, suggesting a disruptive deliberation process, such as induced by over-engagement with threat, during an orientative stage or perhaps an extended evaluation stage. Both interpretations may be complementary, but further investigation is required to establish that link. Overall, this experiment paves the way for future research on the relationship between speech, individual differences and emotional language in terms of information channels, anxiety and threatening language.

## Authors Contributions

Contributions to the study were as follows. **Busch-Moreno**: Conceptualization, Methodology, Software, Analysis, Investigation, Resources, Curation, Writing, Visualization, Administration. **Vinson**: Conceptualization, Methodology, Revision, Supervision, Administration. **Tuomainen**: Conceptualization, Resources.

## Data statement

All data, analyses’ scripts, and additional info can be found at our OSF repository: https://osf.io/n5b6h/. DOI: 10.17605/OSF.IO/N5B6H. Partial results from this experiment were presented at the Federation of European Neuroscience Societies (FENS) virtual forum, July 2020. A previous, not updated, preprint version of this paper is published in BioArxiv, DOI: https://doi.org/10.1101/2020.10.02.323642.

## Declaration of interests

None.

## Funding Sources

This research is supported by CONICYT Becas Chile 72170145 (https://www.conicyt.cl/).

## Notes

https://osf.io/n5b6h/?view_only=358e8b2d52684c92ac63aded03b22325

### Competing Interest Statement

The authors have declared no competing interest.

### Summary of Updates

Correction of behavioural results.

## References

Ablin, P.; Cardoso, J.; Gramfort, A. (2017). Faster independent component analysis by preconditioning with Hessian approximations. ArXiv Preprint, https://arxiv.org/abs/1706.08171

Astésano, C., Besson, M., & Alter, K. (2004). Brain potentials during semantic and prosodic processing in French. Cognitive Brain Research, 18(2), 172–184. https://doi.org/10.1016/j.cogbrainres.2003.10.002

Banse, R., & Scherer, K. R. (1996). Acoustic profiles in vocal emotion expression. Journal of Personality and Social Psychology, 70(3), 614–636. https://doi.org/10.1037//0022-3514.70.3.614

Bar-Haim, Y., Lamy, D., Pergamin, L., Bakermans-Kranenburg, M. J., & van IJzendoorn, M. H. (2007). Threat-related attentional bias in anxious and nonanxious individuals: A meta-analytic study. Psychological Bulletin, 133(1), 1–24. https://doi.org/10.1037/0033-2909.133.1.1

Belin, P., Fecteau, S., & Bédard, C. (2004). Thinking the voice: neural correlates of voice perception. Trends in Cognitive Sciences, 8(3), 129–135. https://doi.org/10.1016/j.tics.2004.01.008

Bruder, G. E., Schneier, F. R., Stewart, J. W., McGrath, P. J., & Quitkin, F. (2004). Left Hemisphere Dysfunction During Verbal Dichotic Listening Tests in Patients Who Have Social Phobia With or Without Comorbid Depressive Disorder. American Journal of Psychiatry, 161(1), 72–78. https://doi.org/10.1176/appi.ajp.161.1.72

Chen, X., Zhao, L., Jiang, A., & Yang, Y. (2011). Event-related potential correlates of the expectancy violation effect during emotional prosody processing. Biological Psychology, 86(3), 158–167. https://doi.org/10.1016/j.biopsycho.2010.11.004

Cisler, J. M., & Koster, E. H. W. (2010). Mechanisms of attentional biases towards threat in anxiety disorders: An integrative review. Clinical Psychology Review, 30(2), 203–216. https://doi.org/10.1016/j.cpr.2009.11.003

Corr, P. J., & Cooper, A. J. (2016). The Reinforcement Sensitivity Theory of Personality Questionnaire (RST-PQ): Development and validation. Psychological Assessment, 28(11), 1427–1440. https://doi.org/10.1037/pas0000273

Corr, P. J., & McNaughton, N. (2012). Neuroscience and approach/avoidance personality traits: A two stage (valuation–motivation) approach. Neuroscience & Biobehavioral Reviews, 36(10), 2339–2354. https://doi.org/10.1016/j.neubiorev.2012.09.013

De Pascalis, V., Cozzuto, G., Caprara, G. V., & Alessandri, G. (2013). Relations among EEG-alpha asymmetry, BIS/BAS, and dispositional optimism. Biological Psychology, 94(1), 198–209. https://doi.org/10.1016/j.biopsycho.2013.05.016

Dien, J. (1998). Issues in the application of the average reference: Review, critiques, and recommendations. Behavior Research Methods, Instruments, & Computers, 30(1), 34–43. doi: 10.3758/bf03209414

Finnigan, S. (2002). ERP “old/new” effects: memory strength and decisional factor(s). Neuropsychologia, 40(13), 2288–2304. https://doi.org/10.1016/s0028-3932(02)00113-6

Gable, P. A., Neal, L. B., & Threadgill, A. H. (2017). Regulatory behavior and frontal activity: Considering the role of revised-BIS in relative right frontal asymmetry. Psychophysiology, 55(1), e12910. https://doi.org/10.1111/psyp.12910

Gadea, M., Espert, R., Salvador, A., & Martí-Bonmatí, L. (2011). The sad, the angry, and the asymmetrical brain: Dichotic Listening studies of negative affect and depression. Brain and Cognition, 76(2), 294–299. https://doi.org/10.1016/j.bandc.2011.03.003

Godfrey, H. K., & Grimshaw, G. M. (2015). Emotional language is all right: Emotional prosody reduces hemispheric asymmetry for linguistic processing. Laterality: Asymmetries of Body, Brain and Cognition, 21(4–6), 568–584. https://doi.org/10.1080/1357650x.2015.1096940

Gramfort, A., Luessi, M., Larson, E., Engemann, D. A., Strohmeier, D., Brodbeck, C., … Hämäläinen, M. S. (2014). MNE software for processing MEG and EEG data. NeuroImage, 86(2014), 446–460. https://doi.org/10.1016/j.neuroimage.2013.10.027

Grimshaw, G. M., Kwasny, K. M., Covell, E., & Johnson, R. A. (2003). The dynamic nature of language lateralization: effects of lexical and prosodic factors. Neuropsychologia, 41(8), 1008–1019. https://doi.org/10.1016/s0028-3932(02)00315-9

Hajcak, G., MacNamara, A., & Olvet, D. M. (2010). Event-Related Potentials, Emotion, and Emotion Regulation: An Integrative Review. Developmental Neuropsychology, 35(2), 129–155. https://doi.org/10.1080/87565640903526504

Hammerschmidt, K., & Jürgens, U. (2007). Acoustical Correlates of Affective Prosody. Journal of Voice, 21(5), 531–540. https://doi.org/10.1016/j.jvoice.2006.03.002

Heller, W., Nitschke, J. B., & Lindsay, D. L. (1997). Neuropsychological Correlates of Arousal in Self-reported Emotion. Cognition & Emotion, 11(4), 383–402. https://doi.org/10.1080/026999397379854

Hugdahl, K. (2011). Fifty years of dichotic listening research – Still going and going and…. Brain and Cognition, 76(2), 211–213. https://doi.org/10.1016/j.bandc.2011.03.006

Hunter, J. D. (2007). Matplotlib: A 2D Graphics Environment. Computing in Science & Engineering, 9(3), 90–95. https://doi.org/10.1109/mcse.2007.55

Jadoul, Y., Thompson, B., & de Boer, B. (2018). Introducing Parselmouth: A Python interface to Praat. Journal of Phonetics, 71(71), 1–15. https://doi.org/10.1016/j.wocn.2018.07.001

Jas, M., Engemann, D., Bekhti, Y., Raimondo, F., & Gramfort, A. (2017). Autoreject: Automated artifact rejection for MEG and EEG data. Neuroimage, 159, 417–429. doi: 10.1016/j.neuroimage.2017.06.030

Jas, M., Larson, E., Engemann, D., Leppäkangas, J., Taulu, S., Hämäläinen, M., & Gramfort, A. (2018). A Reproducible MEG/EEG Group Study With the MNE Software: Recommendations, Quality Assessments, and Good Practices. Frontiers In Neuroscience, 12. doi: 10.3389/fnins.2018.00530

Keune, P. M., Bostanov, V., Kotchoubey, B., & Hautzinger, M. (2012). Mindfulness versus rumination and behavioral inhibition: A perspective from research on frontal brain asymmetry. Personality and Individual Differences, 53(3), 323– 328. https://doi.org/10.1016/j.paid.2012.03.034

Kotz, S. A., & Paulmann, S. (2007). When emotional prosody and semantics dance cheek to cheek: ERP evidence. Brain Research, 1151(2007), 107–118. https://doi.org/10.1016/j.brainres.2007.03.015

Kotz, S. A., & Paulmann, S. (2011). Emotion, Language, and the Brain. Language and Linguistics Compass, 5(3), 108–125. https://doi.org/10.1111/j.1749-818x.2010.00267.x

Kruschke, J. K. (2013). Bayesian estimation supersedes the t test. Journal of Experimental Psychology: General, 142(2), 573–603. https://doi.org/10.1037/a0029146

Kruschke, J. K. (2015). Doing Bayesian data analysis: a tutorial with R, JAGS, and stan. Amsterdam Etc: Elsevier, Academic Press, Cop.

Kumar, R., Carroll, C., Hartikainen, A., & Martin, O. (2019). ArviZ a unified library for exploratory analysis of Bayesian models in Python. Journal of Open Source Software, 4(33), 1143. https://doi.org/10.21105/joss.01143

Lei, X., & Liao, K. (2017). Understanding the Influences of EEG Reference: A Large-Scale Brain Network Perspective. Frontiers In Neuroscience, 11. doi: 10.3389/fnins.2017.00205

Leshem, R. (2018). Trait Anxiety and Attention: Cognitive Functioning as a Function of Attentional Demands. Current Psychology. https://doi.org/10.1007/s12144-018-9884-9

Liebenthal, E., Desai, R., Ellingson, M. M., Ramachandran, B., Desai, A., & Binder, J. R. (2010). Specialization along the Left Superior Temporal Sulcus for Auditory Categorization. Cerebral Cortex, 20(12), 2958–2970. https://doi.org/10.1093/cercor/bhq045

Luck, S. J. (2014). An introduction to the event-related potential technique (Second). MIT Press.

Martin, O. (2018). Bayesian analysis with Python:introduction to statistical modeling and probabilistic programming using PyMC3 and ArviZ. Birmingham, Uk: Packt Publishing.

McElreath, R. (2020). Statistical Rethinking: A Bayesian Course with Examples in R and STAN (Second). CRC Press.

McEvoy, P. M., Mahoney, A. E. J., & Moulds, M. L. (2010). Are worry, rumination, and post-event processing one and the sameã Journal of Anxiety Disorders, 24(5), 509–519. https://doi.org/10.1016/j.janxdis.2010.03.008

McLaughlin, K. A., Borkovec, T. D., & Sibrava, N. J. (2007). The Effects of Worry and Rumination on Affect States and Cognitive Activity. Behavior Therapy, 38(1), 23–38. https://doi.org/10.1016/j.beth.2006.03.003

McNaughton, N., & Gray, J. A. (2000). Anxiolytic action on the behavioural inhibition system implies multiple types of arousal contribute to anxiety. Journal of Affective Disorders, 61(3), 161–176. https://doi.org/10.1016/s0165-0327(00)00344-x

McNaughton, N., Swart, C., Neo, P., Bates, V., & Glue, P. (2013). Anti-anxiety drugs reduce conflict-specific “theta”—A possible human anxiety-specific biomarker. Journal of Affective Disorders, 148(1), 104–111. https://doi.org/10.1016/j.jad.2012.11.057

Moser, J. S., Hartwig, R., Moran, T. P., Jendrusina, A. A., & Kross, E. (2014). Neural markers of positive reappraisal and their associations with trait reappraisal and worry. Journal of Abnormal Psychology, 123(1), 91–105. https://doi.org/10.1037/a0035817

Nalborczyk, L., Perrone-Bertolotti, M., Baeyens, C., Grandchamp, R., Polosan, M., Spinelli, E., Koster, E. H. W., & Lœvenbruck, H. (2017). Orofacial electromyographic correlates of induced verbal rumination. Biological Psychology, 127(127), 53–63. https://doi.org/10.1016/j.biopsycho.2017.04.013

Neal, L. B., & Gable, P. A. (2017). Regulatory control and impulsivity relate to resting frontal activity. Social Cognitive and Affective Neuroscience, 12(9), 1377–1383. https://doi.org/10.1093/scan/nsx080

Neo, P. S.-H., Thurlow, J. K., & McNaughton, N. (2011). Stopping, goal-conflict, trait anxiety and frontal rhythmic power in the stop-signal task. Cognitive, Affective, & Behavioral Neuroscience, 11(4), 485–493. https://doi.org/10.3758/s13415-011-0046-x

Nitschke, J. B., Heller, W., Palmieri, P. A., & Miller, G. A. (1999). Contrasting patterns of brain activity in anxious apprehension and anxious arousal. Psychophysiology, 36(5), 628–637. https://doi.org/10.1111/1469-8986.3650628

Nygaard, L. C., Herold, D. S., & Namy, L. L. (2009). The Semantics of Prosody: Acoustic and Perceptual Evidence of Prosodic Correlates to Word Meaning. Cognitive Science, 33(1), 127–146. https://doi.org/10.1111/j.1551-6709.2008.01007.x

Paulmann, S., Jessen, S., & Kotz, S. A. (2012). It’s special the way you say it: An ERP investigation on the temporal dynamics of two types of prosody. Neuropsychologia, 50(7), 1609–1620. https://doi.org/10.1016/j.neuropsychologia.2012.03.014

Peirce, J., Gray, J. R., Simpson, S., MacAskill, M., Höchenberger, R., Sogo, H., … Lindeløv, J. K. (2019). PsychoPy2: Experiments in behavior made easy. Behavior Research Methods, 51(1), 195–203. https://doi.org/10.3758/s13428-018-01193-y

Pell, M. D., Rothermich, K., Liu, P., Paulmann, S., Sethi, S., & Rigoulot, S. (2015). Preferential decoding of emotion from human non-linguistic vocalizations versus speech prosody. Biological Psychology, 111(2015), 14–25. https://doi.org/10.1016/j.biopsycho.2015.08.008

Peschard, V., Gilboa-Schechtman, E., & Philippot, P. (2016). Selective attention to emotional prosody in social anxiety: a dichotic listening study. Cognition and Emotion, 31(8), 1749–1756. https://doi.org/10.1080/02699931.2016.1261012

Poeppel, D., Idsardi, W. J., & van Wassenhove, V. (2008). Speech perception at the interface of neurobiology and linguistics. Philosophical Transactions of the Royal Society B: Biological Sciences, 363(1493), 1071–1086. https://doi.org/10.1098/rstb.2007.2160

Robinson, O. J., Pike, A. C., Cornwell, B., & Grillon, C. (2019). The translational neural circuitry of anxiety. Journal of Neurology, Neurosurgery & Psychiatry, jnnp-2019-321400. https://doi.org/10.1136/jnnp-2019-321400

Salvatier, J., Wiecki, T. V., & Fonnesbeck, C. (2016). Probabilistic programming in Python using PyMC3. PeerJ Computer Science, 2, e55. https://doi.org/10.7717/peerj-cs.55

Sander, D., Grandjean, D., Pourtois, G., Schwartz, S., Seghier, M. L., Scherer, K. R., & Vuilleumier, P. (2005). Emotion and attention interactions in social cognition: Brain regions involved in processing anger prosody. NeuroImage, 28(4), 848– 858. https://doi.org/10.1016/j.neuroimage.2005.06.023

Sassenhagen, J., Schlesewsky, M., & Bornkessel-Schlesewsky, I. (2014). The P600-as-P3 hypothesis revisited: Single-trial analyses reveal that the late EEG positivity following linguistically deviant material is reaction time aligned. Brain and Language, 137, 29–39. https://doi.org/10.1016/j.bandl.2014.07.010

Schirmer, A., & Kotz, S. A. (2003). ERP Evidence for a Sex-Specific Stroop Effect in Emotional Speech. Journal of Cognitive Neuroscience, 15(8), 1135–1148. https://doi.org/10.1162/089892903322598102

Schirmer, A., & Kotz, S. A. (2006). Beyond the right hemisphere: brain mechanisms mediating vocal emotional processing. Trends in Cognitive Sciences, 10(1), 24– 30. https://doi.org/10.1016/j.tics.2005.11.009

Spielberg, J. M., De Leon, A. A., Bredemeier, K., Heller, W., Engels, A. S., Warren, S. L., … Miller, G. A. (2013). Anxiety type modulates immediate versus delayed engagement of attention-related brain regions. Brain and Behavior, 3(5), 532– 551. https://doi.org/10.1002/brb3.157

Tanner, D., Morgan-Short, K., & Luck, S. J. (2015). How inappropriate high-pass filters can produce artifactual effects and incorrect conclusions in ERP studies of language and cognition. Psychophysiology, 52(8), 997–1009. https://doi.org/10.1111/psyp.12437

van Heuven, W. J. B., Mandera, P., Keuleers, E., & Brysbaert, M. (2014). Subtlex-UK: A New and Improved Word Frequency Database for British English. Quarterly Journal of Experimental Psychology, 67(6), 1176–1190. https://doi.org/10.1080/17470218.2013.850521

VanRullen, R. (2011). Four Common Conceptual Fallacies in Mapping the Time Course of Recognition. Frontiers in Psychology, 2. https://doi.org/10.3389/fpsyg.2011.00365

Wabnitz, P., Martens, U., & Neuner, F. (2015). Written threat: Electrophysiological evidence for an attention bias to affective words in social anxiety disorder. Cognition and Emotion, 30(3), 516–538. https://doi.org/10.1080/02699931.2015.1019837

Warriner, A. B., Kuperman, V., & Brysbaert, M. (2013). Norms of valence, arousal, and dominance for 13,915 English lemmas. Behavior Research Methods, 45(4), 1191–1207. https://doi.org/10.3758/s13428-012-0314-x

Widmann, A., Schröger, E., & Maess, B. (2015). Digital filter design for electrophysiological data – a practical approach. Journal Of Neuroscience Methods, 250, 34–46. doi: 10.1016/j.jneumeth.2014.08.002

Winkler, I., Debener, S., Muller, K., & Tangermann, M. (2015). On the influence of high-pass filtering on ICA-based artifact reduction in EEG-ERP. 2015 37Th Annual International Conference Of The IEEE Engineering In Medicine And Biology Society (EMBC). doi: 10.1109/embc.2015.7319296

Yang, H., Laforge, G., Stojanoski, B., Nichols, E. S., McRae, K., & Köhler, S. (2019). Late positive complex in event-related potentials tracks memory signals when they are decision relevant. Scientific Reports, 9(1). https://doi.org/10.1038/s41598-019-45880-y

Zatorre, R. J., Belin, P., & Penhune, V. B. (2002). Structure and function of auditory cortex: music and speech. Trends in Cognitive Sciences, 6(1), 37–46. https://doi.org/10.1016/s1364-6613(00)01816-7

Zhao, J., Liang, W.-K., Juan, C.-H., Wang, L., Wang, S., & Zhu, Z. (2015). Dissociated stimulus and response conflict effect in the Stroop task: Evidence from evoked brain potentials and brain oscillations. Biological Psychology, 104(2015), 130– 138. https://doi.org/10.1016/j.biopsycho.2014.12.001

