## Supplement 1 for "Trait Anxiety Effects on Late Phase Threatening Speech Processing: Evidence from Electroencephalography"

### Norming

Below we present two supplementary tables related to the rating tasks presented in the main text. Supplementary Table 1.1 summarises the raw averages from threat ratings given in both Semantic and Prosody rating tasks. Supplementary Table 1.2 summarises the posteriors from the ordered-logistic model, showing posterior means, SDs and HDIs (all in ordered log-odds). Note that HDIs for Arousal and Valence overlap zero for the Prosody task, indicating that these semantic measures had little influence on Prosody ratings. In the Semantic task Median Pitch overlaps zero and Hammarberg Index, though it does not overlap zero, is well within the ROPE (2SDs); which indicates little influence of acoustic measures on Semantic ratings. A list of all sentences used for these ratings appears at the end of this supplement.

Supplementary Table 1.1. Average Threat Ratings from Semantic and Prosody Rating Tasks

| Task | Sentence-type | Mean | SD |
|---|---|---|---|
| **Prosody** | Neutral | 1.13 | 1.51 |
| **Prosody** | Prosody | 5.01 | 2.01 |
| **Semantic** | Neutral | 1.29 | 1.83 |
| **Semantic** | Prosody | 5.60 | 2.19 |

Supplementary Table 1.2. Ordered-logistic Model Estimates

| | | **Prosody** | | |
|---|---|---|---|---|
| **Measure** | **Mean** | **SD** | **HDI 5%** | **HDI 95%** |
| **Median Pitch** | 0.033 | 0.003 | 0.029 | 0.037 |
| **Hammarberg Index** | -0.113 | 0.013 | -0.134 | -0.091 |
| **Arousal** | 0.007 | 0.105 | -0.167 | 0.168 |
| **Valence** | -0.074 | 0.158 | -0.319 | 0.199 |
| | | **Semantic** | | |
| **Measure** | **Mean** | **SD** | **HDI 5%** | **HDI 95%** |
| **Median Pitch** | -0.012 | 0.010 | -0.029 | 0.004 |
| **Hammarberg Index** | 0.028 | 0.014 | 0.002 | 0.050 |
| **Arousal** | 0.518 | 0.083 | 0.385 | 0.653 |
| **Valence** | -0.624 | 0.079 | -0.755 | -0.497 |

***Note****: Estimates are in ordered log-odds.*

**Stimuli**

Supplementary Table 1.3 presents a list of all the sentences used for rating tasks. Below this list, Supplementary Tables 1.4 and 1.5 contain all of the sentences used for creating the dichotic pairs for the main experiment, with the first column indicating which threatening and neutral sentences were paired for dichotic listening. Note that these match sentences from the first list (Supplementary Table 1.3), excepting neutral sentences, for which 54 sentences were selected at random for the rating task.

Supplementary Table 1.3. List of sentences used for rating tasks.

| Type | Sentence | Word | Val | Aro | MP | HI |
|---|---|---|---|---|---|---|
| **neutral** | Let's try that. | try | 5.64 | 4 | 96.39 | 14.47 |
| **neutral** | I'll get my things together. | thing | 5.55 | 3.43 | 98.34 | 20.98 |
| **neutral** | We went to school together. | school | 5.41 | 4.57 | 98.96 | 17.78 |
| **neutral** | I'm rebranding him. | brand | 5.05 | 2.82 | 91.71 | 21.05 |
| **neutral** | You just tell him. | tell | 5.27 | 3.86 | 93.46 | 15.54 |
| **neutral** | You're packed already. | pack | 5.55 | 4.1 | 98.61 | 19.76 |
| **neutral** | They got the metric system. | system | 5.5 | 3.38 | 97.54 | 22.47 |
| **neutral** | That's how things really are. | thing | 5.55 | 3.43 | 105.35 | 25.76 |
| **neutral** | Technically I'm not a doctor yet. | doctor | 5.93 | 4.05 | 103.39 | 25.78 |
| **neutral** | It was on the dresser. | dresser | 5.28 | 2.58 | 99.33 | 14.38 |
| **neutral** | It's legal to own it. | legal | 5.24 | 3.35 | 105.38 | 20.86 |
| **neutral** | I want to go with him | want | 6 | 5.29 | 107.07 | 24.53 |
| **neutral** | He can read the funny papers. | paper | 5.42 | 3.52 | 102.34 | 26.17 |
| **neutral** | We stay here for two weeks. | week | 5.27 | 3.33 | 109.5 | 18.43 |
| **neutral** | It's moving around in there. | move | 4.95 | 4.24 | 100.5 | 26.4 |
| **neutral** | He was a doctor. | doctor | 5.93 | 4.05 | 98.53 | 19.75 |
| **neutral** | I didn't get your name. | name | 5.62 | 3.04 | 107.37 | 21.69 |
| **neutral** | No, we have one room booked. | room | 5.55 | 3.1 | 106.65 | 31.82 |
| **neutral** | Get a hold of those minutes. | minute | 5.5 | 3.76 | 103.3 | 20.69 |
| **neutral** | He fixes the cable. | cable | 5.9 | 3.52 | 99.89 | 21.55 |
| **neutral** | I'll be back for you. | back | 4.76 | 2.59 | 108.35 | 23.14 |
| **neutral** | Those aren't leather seats. | seat | 5.22 | 3 | 103.24 | 14.88 |
| **neutral** | You sound different. | different | 5.91 | 3.95 | 104.55 | 22.36 |
| **neutral** | You could meet her next time. | time | 5.6 | 3.41 | 103.4 | 22.22 |
| **neutral** | We didn't do nothing. | thing | 5.55 | 3.43 | 102.65 | 21.44 |
| **neutral** | I'm used to the bus. | bus | 5.12 | 3.83 | 98.33 | 18.03 |
| **neutral** | You always say that. | say | 5.91 | 4.43 | 107.35 | 19.9 |
| **neutral** | I have clients to see tomorrow morning. | client | 5.22 | 2.95 | 96.45 | 20.01 |
| **neutral** | I'll give you a clue. | clue | 5.38 | 3.52 | 98.86 | 25.21 |
| **neutral** | He didn't actually say anything. | thing | 5.55 | 3.43 | 96.96 | 18.37 |
| **neutral** | I would remember her name. | name | 5.62 | 3.04 | 98.24 | 28.51 |
| **neutral** | I know we have a standard routine. | routine | 4.95 | 3.15 | 106.48 | 21.09 |
| **neutral** | You should get moving. | move | 4.95 | 4.24 | 100.22 | 19.38 |

| | | | | | | |
|---|---|---|---|---|---|---|
| **neutral** | She asked around. | ask | 5.95 | 3.48 | 97.62 | 15.75 |
| **neutral** | I'll leave that with you. | leave | 4.68 | 4.48 | 95.51 | 22.31 |
| **neutral** | It's in your drawer. | drawer | 4.67 | 3 | 102.48 | 24.31 |
| **neutral** | I just need your signature. | signature | 5.57 | 3.3 | 103.03 | 17.18 |
| **neutral** | Tell us a little bit about it. | tell | 5.27 | 3.86 | 115.59 | 18.49 |
| **neutral** | It's got your name on it. | name | 5.62 | 3.04 | 102.3 | 26.65 |
| **neutral** | I suppose it's cheaper. | cheap | 5.24 | 4.47 | 103.32 | 17 |
| **neutral** | No, she just liked the name. | name | 5.62 | 3.04 | 101.28 | 16.99 |
| **neutral** | You dropped something. | thing | 5.55 | 3.43 | 89.51 | 17.08 |
| **neutral** | I see other guys my age. | age | 5.78 | 3.71 | 99.29 | 20.46 |
| **neutral** | You can use my straw. | straw | 5.89 | 2.35 | 100.56 | 20.67 |
| **neutral** | I've been transferred. | transfer | 5.29 | 3.22 | 98.53 | 18.8 |
| **neutral** | It's under my mother's name. | name | 5.62 | 3.04 | 102.71 | 28.33 |
| **neutral** | That's all I wanted to do. | want | 6 | 5.29 | 108.63 | 21.44 |
| **neutral** | I'll see you in a sec. | second | 5.23 | 3.48 | 123.06 | 18.06 |
| **neutral** | You might have told somebody. | tell | 5.27 | 3.86 | 96.36 | 16.91 |
| **neutral** | Make it work with one line. | line | 4.82 | 3.24 | 98.29 | 25.11 |
| **neutral** | It's under her name. | name | 5.62 | 3.04 | 106.73 | 28.21 |
| **neutral** | Don't get attached to it. | attach | 5.26 | 3.81 | 104.73 | 20.29 |
| **neutral** | He thinks it's good for business. | business | 5.48 | 3.71 | 112.13 | 21.36 |
| **neutral** | The whole office is going. | office | 4.54 | 3.05 | 91.85 | 24.89 |
| **prosody** | I'm at work. | work | 5.05 | 4.33 | 143.18 | 2.54 |
| **prosody** | I'll explain it to you. | explain | 5.41 | 4 | 162.61 | -2.79 |
| **prosody** | Just recap for me. | recap | 5.33 | 3.26 | 181.6 | 4.61 |
| **prosody** | I got tickets. | ticket | 5.28 | 3.95 | 167.93 | 7.55 |
| **prosody** | Look at me. | look | 5.95 | 3.76 | 181.3 | 2.75 |
| **prosody** | I don't want that. | want | 6 | 5.29 | 167.16 | 10.33 |
| **prosody** | You got work to do. | work | 5.05 | 4.33 | 177.32 | 4.19 |
| **prosody** | You see what happens. | happen | 5.5 | 3.43 | 160.81 | 9.77 |
| **prosody** | Let me tell you something. | thing | 5.55 | 3.43 | 184.13 | 10.15 |
| **prosody** | You wait here. | wait | 4.55 | 3.62 | 185.05 | 8.77 |
| **prosody** | She's my assistant. | assistant | 5.91 | 4.8 | 152.6 | 8.75 |
| **prosody** | You give me a week. | week | 5.27 | 3.33 | 161.06 | 3.2 |
| **prosody** | There is nothing to tell. | tell | 5.27 | 3.86 | 160.61 | 8.55 |
| **prosody** | You shouldn't say anything. | say | 5.91 | 4.43 | 132.74 | 4.12 |
| **prosody** | This one was a captain. | captain | 5.71 | 3.86 | 192.24 | 8.48 |
| **prosody** | Look at it. | look | 5.95 | 3.76 | 164.21 | 5.38 |
| **prosody** | We'll work it out. | work | 5.05 | 4.33 | 185.64 | 5.67 |
| **prosody** | I need to sit for a minute. | sit | 5.82 | 3.19 | 201.37 | 10.63 |
| **prosody** | I mean, all the time. | time | 5.6 | 3.41 | 179.95 | 2.98 |
| **prosody** | Take off your coat. | coat | 5.29 | 3.1 | 168.96 | 9.37 |
| **prosody** | Wait a minute. | minute | 5.5 | 3.76 | 160.27 | 8.27 |
| **prosody** | You know, acting. | act | 5.64 | 4.19 | 186.54 | 7.41 |
| **prosody** | Get to the point. | point | 5.45 | 3.86 | 170.26 | 7.5 |
| **prosody** | Say what you want. | want | 6 | 5.29 | 174.31 | 10.38 |
| **prosody** | I will tell you. | tell | 5.27 | 3.86 | 180.75 | 13.14 |

| | | | | | | |
|---|---|---|---|---|---|---|
| **prosody** | Look who's here. | look | 5.95 | 3.76 | 172.01 | 6.79 |
| **prosody** | We play by the rules. | rule | 4.5 | 3.67 | 184.33 | 11 |
| **prosody** | You said there was something else. | thing | 5.55 | 3.43 | 158.38 | 9.49 |
| **prosody** | I did the work. | work | 5.05 | 4.33 | 203.14 | 4.49 |
| **prosody** | I haven't seen you in a month. | month | 5.78 | 3.64 | 158.17 | 4.64 |
| **prosody** | She works with me. | work | 5.05 | 4.33 | 196.64 | 9.58 |
| **prosody** | I don't know how many times. | time | 5.6 | 3.41 | 186.51 | 7.88 |
| **prosody** | I'm telling you. | tell | 5.27 | 3.86 | 162.39 | 6.5 |
| **prosody** | Let's get to work. | work | 5.05 | 4.33 | 169.5 | 7 |
| **prosody** | I made it happen. | happen | 5.5 | 3.43 | 183.45 | 4.09 |
| **prosody** | Let's go out. | let | 5.95 | 2.71 | 190.31 | 10.3 |
| **prosody** | Tell him what I said. | say | 5.91 | 4.43 | 162.8 | 10.04 |
| **prosody** | She'll be back in a minute. | minute | 5.5 | 3.76 | 170.06 | 5.52 |
| **prosody** | He's done nothing. | thing | 5.55 | 3.43 | 163.99 | 5.55 |
| **prosody** | He's looking into it. | look | 5.95 | 3.76 | 182.58 | 9.37 |
| **prosody** | This is exactly my point | point | 5.45 | 3.86 | 183.97 | 10.23 |
| **prosody** | Look who's here. | look | 5.95 | 3.76 | 176.46 | 6.45 |
| **prosody** | Tell them I'm through the door. | tell | 5.27 | 3.86 | 159.82 | 11.97 |
| **prosody** | Come and sit down. | sit | 5.82 | 3.19 | 189.79 | 11.71 |
| **prosody** | Just a second here. | second | 5.23 | 3.48 | 160.42 | 8.91 |
| **prosody** | You know the situation. | situation | 5.33 | 4.05 | 195.74 | 11.04 |
| **prosody** | I found it in her bag. | bag | 5.05 | 3.43 | 191.87 | 12.43 |
| **prosody** | I'll say it again. | say | 5.91 | 4.43 | 194.05 | 6.71 |
| **prosody** | Tell me your last name. | name | 5.62 | 3.04 | 184.67 | 4.54 |
| **prosody** | Man, don't say that. | say | 5.91 | 4.43 | 166.5 | 8.64 |
| **prosody** | That's your name. | name | 5.62 | 3.04 | 212.18 | 13.27 |
| **prosody** | Keep your seat. | seat | 5.22 | 3 | 144.42 | 9.04 |
| **prosody** | You and your buildings. | building | 5.47 | 3.35 | 201.04 | 10.07 |
| **prosody** | I'll be on that stage. | stage | 5.67 | 3.76 | 184.16 | 8.83 |
| **semantic** | Blow the roof off this motherfucker. | motherfucker | 1.61 | 7.33 | 98.66 | 17.8 |
| **semantic** | Found it, motherfucker. | motherfucker | 1.61 | 7.33 | 101.13 | 26.36 |
| **semantic** | You mad bastard. | mad | 2.47 | 5.59 | 99.69 | 23.85 |
| **semantic** | Don't be so bloody daft. | bloody | 2.86 | 5.76 | 108.23 | 19.98 |
| **semantic** | Don't give me that bullshit. | bullshit | 2.33 | 7 | 92.95 | 15.7 |
| **semantic** | Don't bullshit me. | bullshit | 2.33 | 7 | 110.68 | 19.44 |
| **semantic** | Where is my goddamn money. | damn | 4.32 | 5.1 | 91.79 | 21.22 |
| **semantic** | You lying son of a bitch. | bitch | 2.55 | 5.95 | 105.53 | 24.54 |
| **semantic** | You insolent bastard. | bastard | 3.31 | 5.1 | 105.35 | 16.59 |
| **semantic** | This is bullshit, man. | bullshit | 2.33 | 7 | 106.92 | 20.12 |
| **semantic** | No more bullshit. | shit | 3.91 | 5.67 | 91.12 | 31.96 |
| **semantic** | You little fucking asshole. | asshole | 2.11 | 6.38 | 110.15 | 27.83 |
| **semantic** | You nobody piece of shit. | shit | 3.91 | 5.67 | 95.94 | 23.66 |
| **semantic** | You're a ruthless little cunt. | cunt | 3.7 | 6.1 | 108.38 | 20.68 |
| **semantic** | What a fucking bitch. | bitch | 2.55 | 5.95 | 103.06 | 17.54 |
| **semantic** | Fuck that motherfucker. | motherfucker | 1.61 | 7.33 | 101.55 | 12.34 |
| **semantic** | Your father was a bitch. | bitch | 2.55 | 5.95 | 101.67 | 25.88 |

| | | | | | | |
|---|---|---|---|---|---|---|
| **semantic** | You are a boring fucking cunt. | cunt | 3.7 | 6.1 | 100.27 | 29.16 |
| **semantic** | You fucking faggot. | faggot | 2.32 | 5.68 | 95.48 | 19.61 |
| **semantic** | Shoot him in the face. | shoot | 3.5 | 6 | 102.66 | 20.03 |
| **semantic** | Don't move, motherfucker. | motherfucker | 1.61 | 7.33 | 109.4 | 22.28 |
| **semantic** | Just give me the gun. | gun | 3.66 | 7.74 | 100.13 | 19.68 |
| **semantic** | Call that little shit. | shit | 3.91 | 5.67 | 103.27 | 25.29 |
| **semantic** | You bastards murdered her. | murder | 1.48 | 6.24 | 101.59 | 23.01 |
| **semantic** | I want to shoot the pigeons. | shoot | 3.5 | 6 | 102.98 | 18.91 |
| **semantic** | You're such an asshole. | asshole | 2.11 | 6.38 | 103.47 | 25.78 |
| **semantic** | The bitch shot my finger off. | shoot | 3.5 | 6 | 101.32 | 22.74 |
| **semantic** | Get out of the way, asshole. | asshole | 2.11 | 6.38 | 96 | 16.72 |
| **semantic** | Yes, I have killed people. | kill | 1.81 | 6.81 | 94.35 | 16.23 |
| **semantic** | Because I kill the motherfucker. | kill | 1.81 | 6.81 | 116.68 | 18.23 |
| **semantic** | You vicious fucking cunt. | cunt | 3.7 | 6.1 | 113.17 | 13.36 |
| **semantic** | I start killing people. | kill | 1.81 | 6.81 | 100.22 | 21.96 |
| **semantic** | Tell me, you bastard. | bastard | 3.31 | 5.1 | 103.5 | 26.75 |
| **semantic** | Fucking asshole dog-fucker. | asshole | 2.11 | 6.38 | 99.06 | 21.5 |
| **semantic** | Shoot him in the kneecap. | shoot | 3.5 | 6 | 109.86 | 21.79 |
| **semantic** | Don't give me that shit. | shit | 3.91 | 5.67 | 98.21 | 19.14 |
| **semantic** | I shoot you, you go down. | shoot | 3.5 | 6 | 108.11 | 22.17 |
| **semantic** | I'm gonna fucking kill you. | kill | 1.81 | 6.81 | 102.05 | 24.43 |
| **semantic** | Screw you, motherfucker. | motherfucker | 1.61 | 7.33 | 105.43 | 21.8 |
| **semantic** | Fucking shoot me. | shoot | 3.5 | 6 | 95.19 | 22.42 |
| **semantic** | They should be fucking killed. | kill | 1.81 | 6.81 | 95.17 | 26.75 |
| **semantic** | You've always been a cunt. | cunt | 3.7 | 6.1 | 112.94 | 15.95 |
| **semantic** | Keep that motherfucker here. | motherfucker | 1.61 | 7.33 | 99.1 | 16.72 |
| **semantic** | We batted them sons of bitches. | bitch | 2.55 | 5.95 | 96.64 | 18.82 |
| **semantic** | I'm going to fucking stab you. | stab | 3.05 | 5.17 | 105.51 | 17.29 |
| **semantic** | I shoot people for money. | shoot | 3.5 | 6 | 103.06 | 29.69 |
| **semantic** | You poisonous little toad. | poisonous | 2.52 | 5.67 | 97.96 | 20.36 |
| **semantic** | You hurt him, you die. | die | 1.67 | 6.9 | 106.42 | 22.79 |
| **semantic** | It will end in violence. | violence | 2.71 | 5.95 | 98.93 | 20.33 |
| **semantic** | Shut up, bitch. | bitch | 2.55 | 5.95 | 99.81 | 16.01 |
| **semantic** | You muggy cunt. | cunt | 3.7 | 6.1 | 104.04 | 23.89 |
| **semantic** | Fuck you, you cunts. | cunt | 3.7 | 6.1 | 101.58 | 14.59 |
| **semantic** | I didn't kill anybody. | kill | 1.81 | 6.81 | 106.65 | 22.4 |
| **semantic** | Because you're a cunt. | cunt | 3.7 | 6.1 | 100.88 | 24.32 |
| **congruent** | You'd be a dead man. | cunt | 3.7 | 6.1 | 216.35 | 7.08 |
| **congruent** | You know what this shit is. | shit | 3.91 | 5.67 | 207.16 | -0.68 |
| **congruent** | Get that shit up. | shit | 3.91 | 5.67 | 172.62 | 11.71 |
| **congruent** | You dirty, murdering bastard. | murder | 1.48 | 6.24 | 159.51 | -0.36 |
| **congruent** | I'll kill your fucking car. | kill | 1.81 | 6.81 | 175.97 | 11.46 |
| **congruent** | I could've just killed you now. | kill | 1.81 | 6.81 | 196.67 | 3.53 |
| **congruent** | They're gonna kill her. | kill | 1.81 | 6.81 | 196.73 | 0.27 |
| **congruent** | The guy's a fucking psychopath. | psychopath | 2.62 | 5.41 | 205.06 | 6.32 |
| **congruent** | You nasty piece of shit. | shit | 3.91 | 5.67 | 134.27 | 9.74 |

| | | | | | | |
|---|---|---|---|---|---|---|
| **congruent** | I'm fucking tired of this shit. | shit | 3.91 | 5.67 | 139.84 | 6.88 |
| **congruent** | The bastard's gonna hit me. | bastard | 3.31 | 5.1 | 182.79 | 4.52 |
| **congruent** | Get off of me, bitch. | bitch | 2.55 | 5.95 | 159.87 | 3.89 |
| **congruent** | I killed a little boy. | kill | 1.81 | 6.81 | 125.39 | 0.05 |
| **congruent** | Asshole, get out of here. | asshole | 2.11 | 6.38 | 153.28 | 3.74 |
| **congruent** | Shoot the bastards. | shoot | 3.5 | 6 | 153.33 | 9.74 |
| **congruent** | Talk to me, you motherfucker. | motherfucker | 1.61 | 7.33 | 178.92 | 5.44 |
| **congruent** | You little shit. | shit | 3.91 | 5.67 | 173.48 | 7.78 |
| **congruent** | Those Communist motherfuckers. | motherfucker | 1.61 | 7.33 | 113.37 | 6.43 |
| **congruent** | You're a fucking fraud. | fraud | 2.05 | 5.18 | 135.6 | 1.08 |
| **congruent** | There's a line, asshole | asshole | 2.11 | 6.38 | 128.34 | 8.3 |
| **congruent** | You drive around like an asshole. | asshole | 2.11 | 6.38 | 162.97 | 3.38 |
| **congruent** | I said hold that shit. | shit | 3.91 | 5.67 | 129.02 | 7.04 |
| **congruent** | You want to bullshit me. | bullshit | 2.33 | 7 | 168.28 | 9.05 |
| **congruent** | Do it, goddammit. | damn | 4.32 | 5.1 | 166.36 | 4.9 |
| **congruent** | Dry me off, you cunt. | cunt | 3.7 | 6.1 | 156.18 | 5.6 |
| **congruent** | You stop talking shit. | shit | 3.91 | 5.67 | 152.66 | 14.65 |
| **congruent** | You've ruined my life. | ruin | 2.32 | 5.4 | 154.61 | 2.56 |
| **congruent** | Shut that motherfucker. | motherfucker | 1.61 | 7.33 | 189.48 | 8.13 |
| **congruent** | There's bombs everywhere. | bomb | 2.47 | 5.71 | 174.3 | 7.08 |
| **congruent** | You drunk motherfucker. | motherfucker | 1.61 | 7.33 | 187.35 | 15.88 |
| **congruent** | Shoot the fucker. | shoot | 3.5 | 6 | 162.26 | 7.51 |
| **congruent** | You little fucking cunt. | cunt | 3.7 | 6.1 | 173.67 | 16.04 |
| **congruent** | He killed your friend. | kill | 1.81 | 6.81 | 173.95 | 13.42 |
| **congruent** | Let him go, you bastards. | bastard | 3.31 | 5.1 | 148.42 | 9.2 |
| **congruent** | You're a real tight fucker. | fucker | 3.52 | 6.32 | 158.72 | 11 |
| **congruent** | I am shitting you. | shit | 3.91 | 5.67 | 202.63 | 10.34 |
| **congruent** | He puts a gun to my head. | gun | 3.66 | 7.74 | 164.14 | 10.33 |
| **congruent** | I fucking shot her. | shoot | 3.5 | 6 | 148.18 | 11.74 |
| **congruent** | I'm gonna rip your eyes out. | rip | 3.28 | 5.26 | 172.66 | 9.59 |
| **congruent** | It really makes me hate you. | hate | 1.96 | 6.26 | 197.15 | 8.25 |
| **congruent** | I wanted to kill this little fuck. | kill | 1.81 | 6.81 | 193.94 | 11.76 |
| **congruent** | Stop that shit. | shit | 3.91 | 5.67 | 192.82 | 6.73 |
| **congruent** | You son of a bitch. | bitch | 2.55 | 5.95 | 187.64 | 9.71 |
| **congruent** | I'll shoot you in the cock. | shoot | 3.5 | 6 | 191.14 | 7.86 |
| **congruent** | You're a fucking lunatic. | lunatic | 3.47 | 5.64 | 179.25 | 7.28 |
| **congruent** | You've had your friend killed. | kill | 1.81 | 6.81 | 169.15 | 10.23 |
| **congruent** | I'm gonna bash his fucking head. | bash | 3.68 | 5.14 | 177.49 | 7.08 |
| **congruent** | Wake up, you piece of shit. | shit | 3.91 | 5.67 | 155.28 | 10.76 |
| **congruent** | You people are full of shit. | shit | 3.91 | 5.67 | 147.47 | 8.9 |
| **congruent** | I will kill you very slowly. | kill | 1.81 | 6.81 | 160.92 | 17.86 |
| **congruent** | Die, you motherfucker. | die | 1.67 | 6.9 | 159.31 | 11.21 |
| **congruent** | You fucking bullshitter. | bullshit | 2.33 | 7 | 174.91 | 10.36 |
| **congruent** | I'm shooting the dog | shoot | 3.5 | 6 | 137.55 | 2.39 |
| **congruent** | You suicidal self-loathing and shit. | shit | 3.91 | 5.67 | 159.44 | 12.08 |

Supplementary Table 1.4. List of threatening sentences used for main experiment's dichotic pair

| Pair# | Threat Sentence | Word | Val | Aro | Freq | Movie |
|---|---|---|---|---|---|---|
| **nstp001** | I'm at work. | work | 5.05 | 4.33 | 5.99 | HotFuzz |
| **nstp002** | I'll explain it to you. | explain | 5.41 | 4 | 4.83 | WhateverWorks |
| **nstp003** | Just recap for me. | recap | 5.33 | 3.26 | 3.68 | InLoop |
| **nstp004** | I got tickets. | ticket | 5.28 | 3.95 | 4.51 | EasternPromises |
| **nstp005** | Look at me. | look | 5.95 | 3.76 | 6.23 | Goodfellas |
| **nstp006** | I don't want that. | want | 6 | 5.29 | 6.22 | InNameFather |
| **nstp007** | You got work to do. | work | 5.05 | 4.33 | 5.99 | EasternPromises |
| **nstp008** | You see what happens. | happen | 5.5 | 3.43 | 5.32 | BigLebowski |
| **nstp009** | Let me tell you something. | thing | 5.55 | 3.43 | 5.94 | BigLebowski |
| **nstp010** | You wait here. | wait | 4.55 | 3.62 | 5.37 | DeathFuneral |
| **nstp011** | She's my assistant. | assistant | 5.91 | 4.8 | 4.29 | Birdman |
| **nstp012** | You give me a week. | week | 5.27 | 3.33 | 5.66 | InLoop |
| **nstp013** | There is nothing to tell. | tell | 5.27 | 3.86 | 5.82 | EasternPromises |
| **nstp014** | You shouldn't say anything. | say | 5.91 | 4.43 | 6.13 | InLoop |
| **nstp015** | This one was a captain. | captain | 5.71 | 3.86 | 4.85 | EasternPromises |
| **nstp016** | Look at it. | look | 5.95 | 3.76 | 6.23 | BigLebowski |
| **nstp017** | We'll work it out. | work | 5.05 | 4.33 | 5.99 | DeathFuneral |
| **nstp018** | I need to sit for a minute. | sit | 5.82 | 3.19 | 5.14 | WhateverWorks |
| **nstp019** | I mean, all the time. | time | 5.6 | 3.41 | 6.35 | DeconstructingHarry |
| **nstp020** | Take off your coat. | coat | 5.29 | 3.1 | 4.41 | DeconstructingHarry |
| **nstp021** | Wait a minute. | minute | 5.5 | 3.76 | 5.23 | DeathFuneral |
| **nstp022** | You know, acting. | act | 5.64 | 4.19 | 4.99 | Birdman |
| **nstp023** | Get to the point. | point | 5.45 | 3.86 | 5.60 | InNameFather |
| **nstp024** | Say what you want. | want | 6 | 5.29 | 6.22 | Birdman |
| **nstp025** | I will tell you. | tell | 5.27 | 3.86 | 5.82 | EasternPromises |
| **nstp026** | Look who's here. | look | 5.95 | 3.76 | 6.23 | Goodfellas |
| **nstp027** | We play by the rules. | rule | 4.5 | 3.67 | 4.67 | Snatch |
| **nstp028** | You said there was something else. | thing | 5.55 | 3.43 | 5.94 | InLoop |
| **nstp029** | I did the work. | work | 5.05 | 4.33 | 5.99 | Birdman |
| **nstp030** | I haven't seen you in a month. | month | 5.78 | 3.64 | 5.08 | InLoop |
| **nstp031** | She works with me. | work | 5.05 | 4.33 | 5.99 | Birdman |
| **nstp032** | I don't know how many times. | time | 5.6 | 3.41 | 6.35 | HotFuzz |
| **nstp033** | I'm telling you. | tell | 5.27 | 3.86 | 5.82 | HitmansBodyguard |
| **nstp034** | Let's get to work. | work | 5.05 | 4.33 | 5.99 | Birdman |
| **nstp035** | I made it happen. | happen | 5.5 | 3.43 | 5.32 | Birdman |
| **nstp036** | Let's go out. | let | 5.95 | 2.71 | 6.16 | InBruges |
| **nstp037** | Tell him what I said. | say | 5.91 | 4.43 | 6.13 | EasternPromises |
| **nstp038** | She'll be back in a minute. | minute | 5.5 | 3.76 | 5.23 | BigLebowski |
| **nstp039** | He's done nothing. | thing | 5.55 | 3.43 | 5.94 | InNameFather |
| **nstp040** | He's looking into it. | look | 5.95 | 3.76 | 6.23 | Goodfellas |
| **nstp041** | This is exactly my point | point | 5.45 | 3.86 | 5.60 | InBruges |
| **nstp042** | Look who's here. | look | 5.95 | 3.76 | 6.23 | Goodfellas |
| **nstp043** | Tell them I'm through the door. | tell | 5.27 | 3.86 | 5.82 | EasternPromises |

| | | | | | | |
|---|---|---|---|---|---|---|
| **nstp044** | Come and sit down. | sit | 5.82 | 3.19 | 5.14 | DeathFuneral |
| **nstp045** | Just a second here. | second | 5.23 | 3.48 | 5.61 | HitmansBodyguard |
| **nstp046** | You know the situation. | situation | 5.33 | 4.05 | 5.09 | HotFuzz |
| **nstp047** | I found it in her bag. | bag | 5.05 | 3.43 | 4.89 | EasternPromises |
| **nstp048** | I'll say it again. | say | 5.91 | 4.43 | 6.13 | BigLebowski |
| **nstp049** | Tell me your last name. | name | 5.62 | 3.04 | 5.61 | EasternPromises |
| **nstp050** | Man, don't say that. | say | 5.91 | 4.43 | 6.13 | BigLebowski |
| **nstp051** | That's your name. | name | 5.62 | 3.04 | 5.61 | BigLebowski |
| **nstp052** | Keep your seat. | seat | 5.22 | 3 | 4.78 | HotFuzz |
| **nstp053** | You and your buildings. | building | 5.47 | 3.35 | 5.24 | InBruges |
| **nstp054** | I'll be on that stage. | stage | 5.67 | 3.76 | 5.19 | Birdman |
| **tsnp001** | Blow the roof off this motherfucker. | motherfucker | 1.61 | 7.33 | 3.32 | WolfWallstreet |
| **tsnp002** | Found it, motherfucker. | motherfucker | 1.61 | 7.33 | 3.32 | SevenPsychos |
| **tsnp003** | You mad bastard. | mad | 2.47 | 5.59 | 4.73 | InNameFather |
| **tsnp004** | Don't be so bloody daft. | bloody | 2.86 | 5.76 | 4.66 | TheBigShort |
| **tsnp005** | Don't give me that bullshit. | bullshit | 2.33 | 7 | 3.68 | DeconstructingHarry |
| **tsnp006** | Don't bullshit me. | bullshit | 2.33 | 7 | 3.68 | TheBigShort |
| **tsnp007** | Where is my goddamn money. | damn | 4.32 | 5.1 | 4.28 | BigLebowski |
| **tsnp008** | You lying son of a bitch. | bitch | 2.55 | 5.95 | 4.09 | Goodfellas |
| **tsnp009** | You insolent bastard. | bastard | 3.31 | 5.1 | 4.00 | RocknRolla |
| **tsnp010** | This is bullshit, man. | bullshit | 2.33 | 7 | 3.68 | InsideMan |
| **tsnp011** | No more bullshit. | shit | 3.91 | 5.67 | 4.68 | Goodfellas |
| **tsnp012** | You little fucking asshole. | asshole | 2.11 | 6.38 | 3.34 | DeconstructingHarry |
| **tsnp013** | You nobody piece of shit. | shit | 3.91 | 5.67 | 4.68 | Birdman |
| **tsnp014** | You're a ruthless little cunt. | cunt | 3.7 | 6.1 | 3.03 | Snatch |
| **tsnp015** | What a fucking bitch. | bitch | 2.55 | 5.95 | 4.09 | PulpFiction |
| **tsnp016** | Fuck that motherfucker. | motherfucker | 1.61 | 7.33 | 3.32 | WolfWallstreet |
| **tsnp017** | Your father was a bitch. | bitch | 2.55 | 5.95 | 4.09 | EasternPromises |
| **tsnp018** | You are a boring fucking cunt. | cunt | 3.7 | 6.1 | 3.03 | InLoop |
| **tsnp019** | You fucking faggot. | faggot | 2.32 | 5.68 | 2.81 | Harry Brown |
| **tsnp020** | Shoot him in the face. | shoot | 3.5 | 6 | 4.53 | PulpFiction |
| **tsnp021** | Don't move, motherfucker. | motherfucker | 1.61 | 7.33 | 3.32 | Goodfellas |
| **tsnp022** | Just give me the gun. | gun | 3.66 | 7.74 | 4.60 | Birdman |
| **tsnp023** | Call that little shit. | shit | 3.91 | 5.67 | 4.68 | TheBigShort |
| **tsnp024** | You bastards murdered her. | murder | 1.48 | 6.24 | 4.70 | EasternPromises |
| **tsnp025** | I want to shoot the pigeons. | shoot | 3.5 | 6 | 4.53 | Harry Brown |
| **tsnp026** | You're such an asshole. | asshole | 2.11 | 6.38 | 3.34 | Birdman |
| **tsnp027** | The bitch shot my finger off. | shoot | 3.5 | 6 | 4.53 | Revolver |
| **tsnp028** | Get out of the way, asshole. | asshole | 2.11 | 6.38 | 3.34 | WolfWallstreet |
| **tsnp029** | Yes, I have killed people. | kill | 1.81 | 6.81 | 4.91 | InBruges |
| **tsnp030** | Because I kill the motherfucker. | kill | 1.81 | 6.81 | 4.91 | PulpFiction |
| **tsnp031** | You vicious fucking cunt. | cunt | 3.7 | 6.1 | 3.03 | WolfWallstreet |
| **tsnp032** | I start killing people. | kill | 1.81 | 6.81 | 4.91 | InsideMan |
| **tsnp033** | Tell me, you bastard. | bastard | 3.31 | 5.1 | 4.00 | EasternPromises |
| **tsnp034** | Fucking asshole dog-fucker. | asshole | 2.11 | 6.38 | 3.34 | SevenPsychos |
| **tsnp035** | Shoot him in the kneecap. | shoot | 3.5 | 6 | 4.53 | Revolver |

| | | | | | | |
|---|---|---|---|---|---|---|
| **tsnp036** | Don't give me that shit. | shit | 3.91 | 5.67 | 4.68 | Goodfellas |
| **tsnp037** | I shoot you, you go down. | shoot | 3.5 | 6 | 4.53 | Snatch |
| **tsnp038** | I'm gonna fucking kill you. | kill | 1.81 | 6.81 | 4.91 | BigLebowski |
| **tsnp039** | Screw you, motherfucker. | motherfucker | 1.61 | 7.33 | 3.32 | InBruges |
| **tsnp040** | Fucking shoot me. | shoot | 3.5 | 6 | 4.53 | NiceGuys |
| **tsnp041** | They should be fucking killed. | kill | 1.81 | 6.81 | 4.91 | PulpFiction |
| **tsnp042** | You've always been a cunt. | cunt | 3.7 | 6.1 | 3.03 | InBruges |
| **tsnp043** | Keep that motherfucker here. | motherfucker | 1.61 | 7.33 | 3.32 | Goodfellas |
| **tsnp044** | We batted them sons of bitches. | bitch | 2.55 | 5.95 | 4.09 | Goodfellas |
| **tsnp045** | I'm going to fucking stab you. | stab | 3.05 | 5.17 | 3.87 | InLoop |
| **tsnp046** | I shoot people for money. | shoot | 3.5 | 6 | 4.53 | InBruges |
| **tsnp047** | You poisonous little toad. | poisonous | 2.52 | 5.67 | 3.65 | RocknRolla |
| **tsnp048** | You hurt him, you die. | die | 1.67 | 6.9 | 4.90 | PulpFiction |
| **tsnp049** | It will end in violence. | violence | 2.71 | 5.95 | 4.60 | InNameFather |
| **tsnp050** | Shut up, bitch. | bitch | 2.55 | 5.95 | 4.09 | EasternPromises |
| **tsnp051** | You muggy cunt. | cunt | 3.7 | 6.1 | 3.03 | Harry Brown |
| **tsnp052** | Fuck you, you cunts. | cunt | 3.7 | 6.1 | 3.03 | SevenPsychos |
| **tsnp053** | I didn't kill anybody. | kill | 1.81 | 6.81 | 4.91 | InsideMan |
| **tsnp054** | Because you're a cunt. | cunt | 3.7 | 6.1 | 3.03 | SevenPsychos |
| **tstp001** | You'd be a dead man. | cunt | 3.7 | 6.1 | 3.03 | InBruges |
| **tstp002** | You know what this shit is. | shit | 3.91 | 5.67 | 4.68 | Harry Brown |
| **tstp003** | Get that shit up. | shit | 3.91 | 5.67 | 4.68 | Harry Brown |
| **tstp004** | You dirty, murdering bastard. | murder | 1.48 | 6.24 | 4.70 | InNameFather |
| **tstp005** | I'll kill your fucking car. | kill | 1.81 | 6.81 | 4.91 | BigLebowski |
| **tstp006** | I could've just killed you now. | kill | 1.81 | 6.81 | 4.91 | SevenPsychos |
| **tstp007** | They're gonna kill her. | kill | 1.81 | 6.81 | 4.91 | BigLebowski |
| **tstp008** | The guy's a fucking psychopath. | psychopath | 2.62 | 5.41 | 3.01 | SevenPsychos |
| **tstp009** | You nasty piece of shit. | shit | 3.91 | 5.67 | 4.68 | Harry Brown |
| **tstp010** | I'm fucking tired of this shit. | shit | 3.91 | 5.67 | 4.68 | InsideMan |
| **tstp011** | The bastard's gonna hit me. | bastard | 3.31 | 5.1 | 4.00 | RocknRolla |
| **tstp012** | Get off of me, bitch. | bitch | 2.55 | 5.95 | 4.09 | DeconstructingHarry |
| **tstp013** | I killed a little boy. | kill | 1.81 | 6.81 | 4.91 | InBruges |
| **tstp014** | Asshole, get out of here. | asshole | 2.11 | 6.38 | 3.34 | WolfWallstreet |
| **tstp015** | Shoot the bastards. | shoot | 3.5 | 6 | 4.53 | InNameFather |
| **tstp016** | Talk to me, you motherfucker. | motherfucker | 1.61 | 7.33 | 3.32 | Revolver |
| **tstp017** | You little shit. | shit | 3.91 | 5.67 | 4.68 | NiceGuys |
| **tstp018** | Those Communist motherfuckers. | motherfucker | 1.61 | 7.33 | 3.32 | SevenPsychos |
| **tstp019** | You're a fucking fraud. | fraud | 2.05 | 5.18 | 4.25 | Birdman |
| **tstp020** | There's a line, asshole | asshole | 2.11 | 6.38 | 3.34 | NiceGuys |
| **tstp021** | You drive around like an asshole. | asshole | 2.11 | 6.38 | 3.34 | NiceGuys |
| **tstp022** | I said hold that shit. | shit | 3.91 | 5.67 | 4.68 | Harry Brown |
| **tstp023** | You want to bullshit me. | bullshit | 2.33 | 7 | 3.68 | InsideMan |
| **tstp024** | Do it, goddammit. | damn | 4.32 | 5.1 | 4.28 | PulpFiction |
| **tstp025** | Dry me off, you cunt. | cunt | 3.7 | 6.1 | 3.03 | InBruges |
| **tstp026** | You stop talking shit. | shit | 3.91 | 5.67 | 4.68 | EasternPromises |
| **tstp027** | You've ruined my life. | ruin | 2.32 | 5.4 | 4.15 | DeconstructingHarry |

| | | | | | | |
|---|---|---|---|---|---|---|
| **tstp028** | Shut that motherfucker. | motherfucker | 1.61 | 7.33 | 3.32 | WolfWallstreet |
| **tstp029** | There's bombs everywhere. | bomb | 2.47 | 5.71 | 4.49 | InNameFather |
| **tstp030** | You drunk motherfucker. | motherfucker | 1.61 | 7.33 | 3.32 | NiceGuys |
| **tstp031** | Shoot the fucker. | shoot | 3.5 | 6 | 4.53 | Revolver |
| **tstp032** | You little fucking cunt. | cunt | 3.7 | 6.1 | 3.03 | InBruges |
| **tstp033** | He killed your friend. | kill | 1.81 | 6.81 | 4.91 | Revolver |
| **tstp034** | Let him go, you bastards. | bastard | 3.31 | 5.1 | 4.00 | InNameFather |
| **tstp035** | You're a real tight fucker. | fucker | 3.52 | 6.32 | 3.14 | Snatch |
| **tstp036** | I am shitting you. | shit | 3.91 | 5.67 | 4.68 | InLoop |
| **tstp037** | He puts a gun to my head. | gun | 3.66 | 7.74 | 4.60 | InNameFather |
| **tstp038** | I fucking shot her. | shoot | 3.5 | 6 | 4.53 | Harry Brown |
| **tstp039** | I'm gonna rip your eyes out. | rip | 3.28 | 5.26 | 4.18 | TheBigShort |
| **tstp040** | It really makes me hate you. | hate | 1.96 | 6.26 | 4.81 | TheBigShort |
| **tstp041** | I wanted to kill this little fuck. | kill | 1.81 | 6.81 | 4.91 | Goodfellas |
| **tstp042** | Stop that shit. | shit | 3.91 | 5.67 | 4.68 | Birdman |
| **tstp043** | You son of a bitch. | bitch | 2.55 | 5.95 | 4.09 | BigLebowski |
| **tstp044** | I'll shoot you in the cock. | shoot | 3.5 | 6 | 4.53 | NiceGuys |
| **tstp045** | You're a fucking lunatic. | lunatic | 3.47 | 5.64 | 3.38 | TheBigShort |
| **tstp046** | You've had your friend killed. | kill | 1.81 | 6.81 | 4.91 | SevenPsychos |
| **tstp047** | I'm gonna bash his fucking head. | bash | 3.68 | 5.14 | 3.71 | TheBigShort |
| **tstp048** | Wake up, you piece of shit. | shit | 3.91 | 5.67 | 4.68 | WolfWallstreet |
| **tstp049** | You people are full of shit. | shit | 3.91 | 5.67 | 4.68 | Birdman |
| **tstp050** | I will kill you very slowly. | kill | 1.81 | 6.81 | 4.91 | RocknRolla |
| **tstp051** | Die, you motherfucker. | die | 1.67 | 6.9 | 4.90 | Goodfellas |
| **tstp052** | You fucking bullshitter. | bullshit | 2.33 | 7 | 3.68 | Goodfellas |
| **tstp053** | I'm shooting the dog | shoot | 3.5 | 6 | 4.53 | Snatch |
| **tstp054** | You suicidal self-loathing and shit. | shit | 3.91 | 5.67 | 4.68 | SevenPsychos |

Supplementary Table 1.5. List of neutral sentences used for main experiment's dichotic pair

| **Pair#** | **Neutral Sentence** | **Word** | **Val** | **Aro** | **Freq** | **Movie** |
|---|---|---|---|---|---|---|
| **nstp001** | Let's try that. | try | 5.64 | 4 | 5.65 | CaptainFantastic |
| **nstp002** | I'll get my things together. | thing | 5.55 | 3.43 | 5.94 | DeathFuneral |
| **nstp003** | We went to school together. | school | 5.41 | 4.57 | 5.45 | RocknRolla |
| **nstp004** | I'm rebranding him. | brand | 5.05 | 2.82 | 4.68 | InLoop |
| **nstp005** | You just tell him. | tell | 5.27 | 3.86 | 5.82 | BigLebowski |
| **nstp006** | You're packed already. | pack | 5.55 | 4.1 | 4.62 | HotFuzz |
| **nstp007** | They got the metric system. | system | 5.5 | 3.38 | 5.26 | PulpFiction |
| **nstp008** | That's how things really are. | thing | 5.55 | 3.43 | 5.94 | AmericanBeauty |
| **nstp009** | Technically I'm not a doctor yet. | doctor | 5.93 | 4.05 | 5.02 | 50/50 |
| **nstp010** | It was on the dresser. | dresser | 5.28 | 2.58 | 3.40 | EasternPromises |
| **nstp011** | It's legal to own it. | legal | 5.24 | 3.35 | 4.75 | PulpFiction |
| **nstp012** | I want to go with him | want | 6 | 5.29 | 6.22 | InNameFather |
| **nstp013** | He can read the funny papers. | paper | 5.42 | 3.52 | 4.94 | InNameFather |
| **nstp014** | We stay here for two weeks. | week | 5.27 | 3.33 | 5.66 | InBruges |

| | | | | | | |
|---|---|---|---|---|---|---|
| **nstp015** | It's moving around in there. | move | 4.95 | 4.24 | 5.49 | DeathFuneral |
| **nstp016** | He was a doctor. | doctor | 5.93 | 4.05 | 5.02 | EasternPromises |
| **nstp017** | I didn't get your name. | name | 5.62 | 3.04 | 5.61 | PulpFiction |
| **nstp018** | No, we have one room booked. | room | 5.55 | 3.1 | 5.60 | InBruges |
| **nstp019** | Get a hold of those minutes. | minute | 5.5 | 3.76 | 5.23 | InLoop |
| **nstp020** | He fixes the cable. | cable | 5.9 | 3.52 | 4.30 | BigLebowski |
| **nstp021** | I'll be back for you. | back | 4.76 | 2.59 | 6.25 | SevenPsychos |
| **nstp022** | Those aren't leather seats. | seat | 5.22 | 3 | 4.78 | HitmansBodyguard |
| **nstp023** | You sound different. | different | 5.91 | 3.95 | 5.66 | SevenPsychos |
| **nstp024** | You could meet her next time. | time | 5.6 | 3.41 | 6.35 | 50/50 |
| **nstp025** | We didn't do nothing. | thing | 5.55 | 3.43 | 5.94 | InNameFather |
| **nstp026** | I'm used to the bus. | bus | 5.12 | 3.83 | 4.74 | 50/50 |
| **nstp027** | You always say that. | say | 5.91 | 4.43 | 6.13 | PulpFiction |
| **nstp028** | I have clients to see tomorrow morning. | client | 5.22 | 2.95 | 4.03 | WhateverWorks |
| **nstp029** | I'll give you a clue. | clue | 5.38 | 3.52 | 4.62 | SevenPsychos |
| **nstp030** | He didn't actually say anything. | thing | 5.55 | 3.43 | 5.94 | InBruges |
| **nstp031** | I would remember her name. | name | 5.62 | 3.04 | 5.61 | EasternPromises |
| **nstp032** | I know we have a standard routine. | routine | 4.95 | 3.15 | 4.39 | WhateverWorks |
| **nstp033** | You should get moving. | move | 4.95 | 4.24 | 5.49 | HitmansBodyguard |
| **nstp034** | She asked around. | ask | 5.95 | 3.48 | 5.41 | SevenPsychos |
| **nstp035** | I'll leave that with you. | leave | 4.68 | 4.48 | 5.49 | RocknRolla |
| **nstp036** | It's in your drawer. | drawer | 4.67 | 3 | 3.98 | 50/50 |
| **nstp037** | I just need your signature. | signature | 5.57 | 3.3 | 4.08 | FullMonty |
| **nstp038** | Tell us a little bit about it. | tell | 5.27 | 3.86 | 5.82 | NiceGuys |
| **nstp039** | It's got your name on it. | name | 5.62 | 3.04 | 5.61 | RocknRolla |
| **nstp040** | I suppose it's cheaper. | cheap | 5.24 | 4.47 | 4.65 | InBruges |
| **nstp041** | No, she just liked the name. | name | 5.62 | 3.04 | 5.61 | InNameFather |
| **nstp042** | You dropped something. | thing | 5.55 | 3.43 | 5.94 | InNameFather |
| **nstp043** | I see other guys my age. | age | 5.78 | 3.71 | 5.18 | DeconstructingHarry |
| **nstp044** | You can use my straw. | straw | 5.89 | 2.35 | 4.17 | PulpFiction |
| **nstp045** | I've been transferred. | transfer | 5.29 | 3.22 | 4.26 | HotFuzz |
| **nstp046** | It's under my mother's name. | name | 5.62 | 3.04 | 5.61 | Goodfellas |
| **nstp047** | That's all I wanted to do. | want | 6 | 5.29 | 6.22 | Goodfellas |
| **nstp048** | I'll see you in a sec. | second | 5.23 | 3.48 | 5.61 | NottingHill |
| **nstp049** | You might have told somebody. | tell | 5.27 | 3.86 | 5.82 | SevenPsychos |
| **nstp050** | Make it work with one line. | line | 4.82 | 3.24 | 5.45 | Birdman |
| **nstp051** | It's under her name. | name | 5.62 | 3.04 | 5.61 | Goodfellas |
| **nstp052** | Don't get attached to it. | attach | 5.26 | 3.81 | 3.65 | Snatch |
| **nstp053** | He thinks it's good for business. | business | 5.48 | 3.71 | 5.47 | Snatch |
| **nstp054** | The whole office is going. | office | 4.54 | 3.05 | 5.13 | 500Summer |
| **tsnp001** | Look at that, we could make a team. | team | 5.91 | 3.38 | 5.67 | RocknRolla |
| **tsnp002** | That's a novel too. | novel | 5.74 | 3.41 | 4.43 | NottingHill |
| **tsnp003** | It counts for something. | thing | 5.55 | 3.43 | 5.94 | FullMonty |
| **tsnp004** | You just can't switch off. | switch | 5.29 | 3.9 | 4.37 | HotFuzz |
| **tsnp005** | I'll have a pint of lager. | lager | 5.4 | 3.11 | 3.56 | HotFuzz |
| **tsnp006** | I'll leave my number. | number | 5.59 | 3.5 | 5.56 | AmericanBeauty |

| | | | | | | |
|---|---|---|---|---|---|---|
| **tsnp007** | We're not going to school today. | school | 5.41 | 4.57 | 5.45 | DeconstructingHarry |
| **tsnp008** | We're business associates. | associate | 5.2 | 3.11 | 3.91 | NiceGuys |
| **tsnp009** | Remember your training. | training | 5.19 | 4.1 | 4.99 | CaptainFantastic |
| **tsnp010** | Boys, get to work. | work | 5.05 | 4.33 | 5.99 | PulpFiction |
| **tsnp011** | You need to tell us. | tell | 5.27 | 3.86 | 5.82 | HitmansBodyguard |
| **tsnp012** | It's been six weeks. | week | 5.27 | 3.33 | 5.66 | InNameFather |
| **tsnp013** | I don't need the script. | script | 5.42 | 3.65 | 4.07 | Birdman |
| **tsnp014** | We like to get things done. | thing | 5.55 | 3.43 | 5.94 | RocknRolla |
| **tsnp015** | This is what happens. | happen | 5.5 | 3.43 | 5.32 | BigLebowski |
| **tsnp016** | She finally did it. | final | 4.56 | 4.4 | 5.46 | CaptainFantastic |
| **tsnp017** | The lads can't lift him. | lift | 5.32 | 4.9 | 4.75 | Snatch |
| **tsnp018** | I'm just trying to do something. | try | 5.64 | 4 | 5.65 | Adaptation |
| **tsnp019** | You'll never guess what. | guess | 5.18 | 3.86 | 5.16 | InBruges |
| **tsnp020** | Here, let me do that. | let | 5.95 | 2.71 | 6.16 | InNameFather |
| **tsnp021** | I came prepared. | prepare | 5.32 | 4.15 | 4.48 | HitmansBodyguard |
| **tsnp022** | He was nominated. | nominate | 5.86 | 3.48 | 3.89 | Birdman |
| **tsnp023** | Give me your left arm. | arm | 5.44 | 3.44 | 4.70 | NiceGuys |
| **tsnp024** | I don't mind telling you. | tell | 5.27 | 3.86 | 5.82 | Snatch |
| **tsnp025** | We could just listen to the radio. | radio | 6 | 3.84 | 4.82 | 50/50 |
| **tsnp026** | Give me one minute. | minute | 5.5 | 3.76 | 5.23 | WolfWallstreet |
| **tsnp027** | You two really look alike. | look | 5.95 | 3.76 | 6.23 | Adaptation |
| **tsnp028** | You know where the lavatory is. | lavatory | 5.18 | 4.21 | 3.27 | Juno |
| **tsnp029** | Come over here for a second. | second | 5.23 | 3.48 | 5.61 | WolfWallstreet |
| **tsnp030** | I'll text you the details. | detail | 5.67 | 3.81 | 4.63 | InLoop |
| **tsnp031** | We do different projects. | project | 5 | 4.55 | 4.94 | TheBigShort |
| **tsnp032** | He will ask for it. | ask | 5.95 | 3.48 | 5.41 | SevenPsychos |
| **tsnp033** | I'm at my place. | place | 5.86 | 3.52 | 5.78 | BigLebowski |
| **tsnp034** | It's not a public area up here. | public | 5.33 | 3.35 | 5.44 | NiceGuys |
| **tsnp035** | My feet are freezing. | freeze | 4.64 | 4 | 4.30 | FullMonty |
| **tsnp036** | I'm gonna ask you again. | ask | 5.95 | 3.48 | 5.41 | NiceGuys |
| **tsnp037** | They always have an excuse. | excuse | 4.52 | 3.61 | 4.86 | DeconstructingHarry |
| **tsnp038** | You're dealing with numbers. | number | 5.59 | 3.5 | 5.56 | WolfWallstreet |
| **tsnp039** | You just got to pick it up. | pick | 5.91 | 3.62 | 5.16 | Snatch |
| **tsnp040** | You've got a visitor. | visitor | 5.27 | 4 | 3.97 | WolfWallstreet |
| **tsnp041** | Don't use your real name. | name | 5.62 | 3.04 | 5.61 | DeconstructingHarry |
| **tsnp042** | I think I missed something. | miss | 4.1 | 3.9 | 5.21 | 500Summer |
| **tsnp043** | I don't use pans. | pan | 5.15 | 3.05 | 4.60 | SevenPsychos |
| **tsnp044** | Here's the time and here's the place. | place | 5.86 | 3.52 | 5.78 | RocknRolla |
| **tsnp045** | Well, I was buying this shirt. | shirt | 5.56 | 2.3 | 4.47 | WhateverWorks |
| **tsnp046** | You can get a little drunk. | drunk | 4.06 | 5.05 | 4.46 | HotFuzz |
| **tsnp047** | Think about it for a second. | second | 5.23 | 3.48 | 5.61 | WolfWallstreet |
| **tsnp048** | Let me have your bag. | bag | 5.05 | 3.43 | 4.89 | CaptainFantastic |
| **tsnp049** | I didn't make that line up. | line | 4.82 | 3.24 | 5.45 | Adaptation |
| **tsnp050** | He's not coming. | come | 5.64 | 3.57 | 6.26 | FullMonty |
| **tsnp051** | It was in his jacket. | jacket | 5.86 | 3.35 | 4.29 | EasternPromises |
| **tsnp052** | It's not a motorcycle. | motorcycle | 5.8 | 5.8 | 3.55 | PulpFiction |

| | | | | | | |
|---|---|---|---|---|---|---|
| **tsnp053** | I'm in the elevator with you. | elevator | 5.95 | 3.65 | 3.26 | DeconstructingHarry |
| **tsnp054** | Open your mouth. | mouth | 5.59 | 4.14 | 4.78 | InNameFather |
| **tstp001** | Nothing's the matter with me. | matter | 5.82 | 4.05 | 5.24 | SevenPsychos |
| **tstp002** | That's what we wanted to ask you. | ask | 5.95 | 3.48 | 5.41 | 500Summer |
| **tstp003** | She looks like you. | look | 5.95 | 3.76 | 6.23 | Goodfellas |
| **tstp004** | It can really only go two ways. | way | 5.91 | 2.9 | 6.12 | Juno |
| **tstp005** | You copy my work every week. | work | 5.05 | 4.33 | 5.99 | Juno |
| **tstp006** | It's changed a lot over the years. | year | 5.75 | 3.33 | 5.92 | DeathFuneral |
| **tstp007** | I have to tell you something. | tell | 5.27 | 3.86 | 5.82 | WhateverWorks |
| **tstp008** | Nah, it was a premeditated act. | act | 5.64 | 4.19 | 4.99 | Juno |
| **tstp009** | I can't wait to tell the guys. | tell | 5.27 | 3.86 | 5.82 | Adaptation |
| **tstp010** | Lift your elbows above your ears. | lift | 5.32 | 4.9 | 4.75 | 50/50 |
| **tstp011** | Looks like a chauffeur to me. | look | 5.95 | 3.76 | 6.23 | NottingHill |
| **tstp012** | It will come back to me. | back | 4.76 | 2.59 | 6.25 | Snatch |
| **tstp013** | Tell me about yourself. | tell | 5.27 | 3.86 | 5.82 | BigLebowski |
| **tstp014** | You can use onions, too. | onion | 5.37 | 4.95 | 4.29 | DeconstructingHarry |
| **tstp015** | I'm going to chat with her. | chat | 5.75 | 4.27 | 4.60 | DeathFuneral |
| **tstp016** | Go get your jacket on. | jacket | 5.86 | 3.35 | 4.29 | FullMonty |
| **tstp017** | We're going up to the glacier. | glacier | 5.5 | 3.88 | 3.46 | CaptainFantastic |
| **tstp018** | You either have it or you don't. | have | 5.86 | 3.52 | 6.90 | DeathFuneral |
| **tstp019** | I go through doors first. | door | 5.43 | 3.19 | 5.26 | HitmansBodyguard |
| **tstp020** | You made your point. | point | 5.45 | 3.86 | 5.60 | CaptainFantastic |
| **tstp021** | This is an important business function. | function | 5.55 | 4.1 | 4.06 | AmericanBeauty |
| **tstp022** | I left my phone in the office. | office | 4.54 | 3.05 | 5.13 | TheBigShort |
| **tstp023** | I mentioned one committee. | committee | 5.52 | 3.56 | 4.77 | InLoop |
| **tstp024** | Let's hang back. | back | 4.76 | 2.59 | 6.25 | PulpFiction |
| **tstp025** | I'm the senior broker here. | broker | 4.15 | 3.77 | 3.38 | WolfWallstreet |
| **tstp026** | This is your third test today. | test | 4.44 | 4.3 | 5.06 | Juno |
| **tstp027** | We'll go back and get him. | back | 4.76 | 2.59 | 6.25 | DeathFuneral |
| **tstp028** | I'm going to the supply room. | room | 5.55 | 3.1 | 5.60 | 500Summer |
| **tstp029** | The evidence did not support him. | evidence | 4.72 | 3.9 | 5.06 | 500Summer |
| **tstp030** | Have to make an appointment. | appointment | 4.43 | 4.8 | 4.19 | FullMonty |
| **tstp031** | I found her will. | will | 5.32 | 2.9 | 6.55 | CaptainFantastic |
| **tstp032** | My story is, whatever works. | work | 5.05 | 4.33 | 5.99 | WhateverWorks |
| **tstp033** | We're moving in together. | move | 4.95 | 4.24 | 5.49 | DeathFuneral |
| **tstp034** | It's all about the details. | detail | 5.67 | 3.81 | 4.75 | RocknRolla |
| **tstp035** | I thought that's what you wanted. | want | 6 | 5.29 | 6.22 | NiceGuys |
| **tstp036** | I saw your watch. | watch | 5.45 | 3.37 | 5.30 | PulpFiction |
| **tstp037** | They paid good money for tickets. | ticket | 5.28 | 3.95 | 4.49 | WhateverWorks |
| **tstp038** | I'm trying to level with you. | level | 5.72 | 2.15 | 5.13 | AmericanBeauty |
| **tstp039** | I'm so sorry I kept you waiting. | wait | 4.55 | 3.62 | 5.37 | AmericanBeauty |
| **tstp040** | You throw your notepad away. | throw | 5.55 | 4.52 | 4.88 | SevenPsychos |
| **tstp041** | I'm supposed to be buying a flat. | flat | 4.43 | 3.04 | 4.99 | DeathFuneral |
| **tstp042** | You should grow a moustache. | moustache | 5.35 | 3.23 | 3.80 | Juno |
| **tstp043** | I'm trying to work out here. | work | 5.05 | 4.33 | 5.99 | AmericanBeauty |
| **tstp044** | It's only that big on the map. | map | 5.81 | 3.95 | 4.52 | InNameFather |

| | | | | | | |
|---|---|---|---|---|---|---|
| **tstp045** | Those were his exact words. | word | 5.77 | 4 | 5.29 | SevenPsychos |
| **tstp046** | You know, a long time ago. | time | 5.6 | 3.41 | 6.35 | Adaptation |
| **tstp047** | No stops till the exit point. | point | 5.45 | 3.86 | 5.60 | HitmansBodyguard |
| **tstp048** | I'm not gonna ask her about this. | ask | 5.95 | 3.48 | 5.41 | 500Summer |
| **tstp049** | I used to watch it every week. | week | 5.27 | 3.33 | 5.66 | 500Summer |
| **tstp050** | It's not building a model airplane. | airplane | 5.25 | 5.62 | 3.15 | Adaptation |
| **tstp051** | I went to the doctor today. | doctor | 5.93 | 4.05 | 5.02 | Juno |
| **tstp052** | I'm doing it right this time. | time | 5.6 | 3.41 | 6.35 | Adaptation |
| **tstp053** | You've got five minutes. | minute | 5.5 | 3.76 | 5.23 | NottingHill |
| **tstp054** | My mom uses color-safe bleach. | bleach | 4.62 | 5.3 | 3.22 | Juno |
