## Supplement 2 for "Trait Anxiety Effects on Late Phase Threatening Speech Processing: Evidence from Electroencephalography"

Supplementary figures below show examples for all relevant electrodes at Window4 from direct-threat only: amplitude as a function of BIS score, ear and stimulus type. Figures from all other time-windows from direct- and indirect-threat can be found in our OSF repository (https://osf.io/n5b6h/).

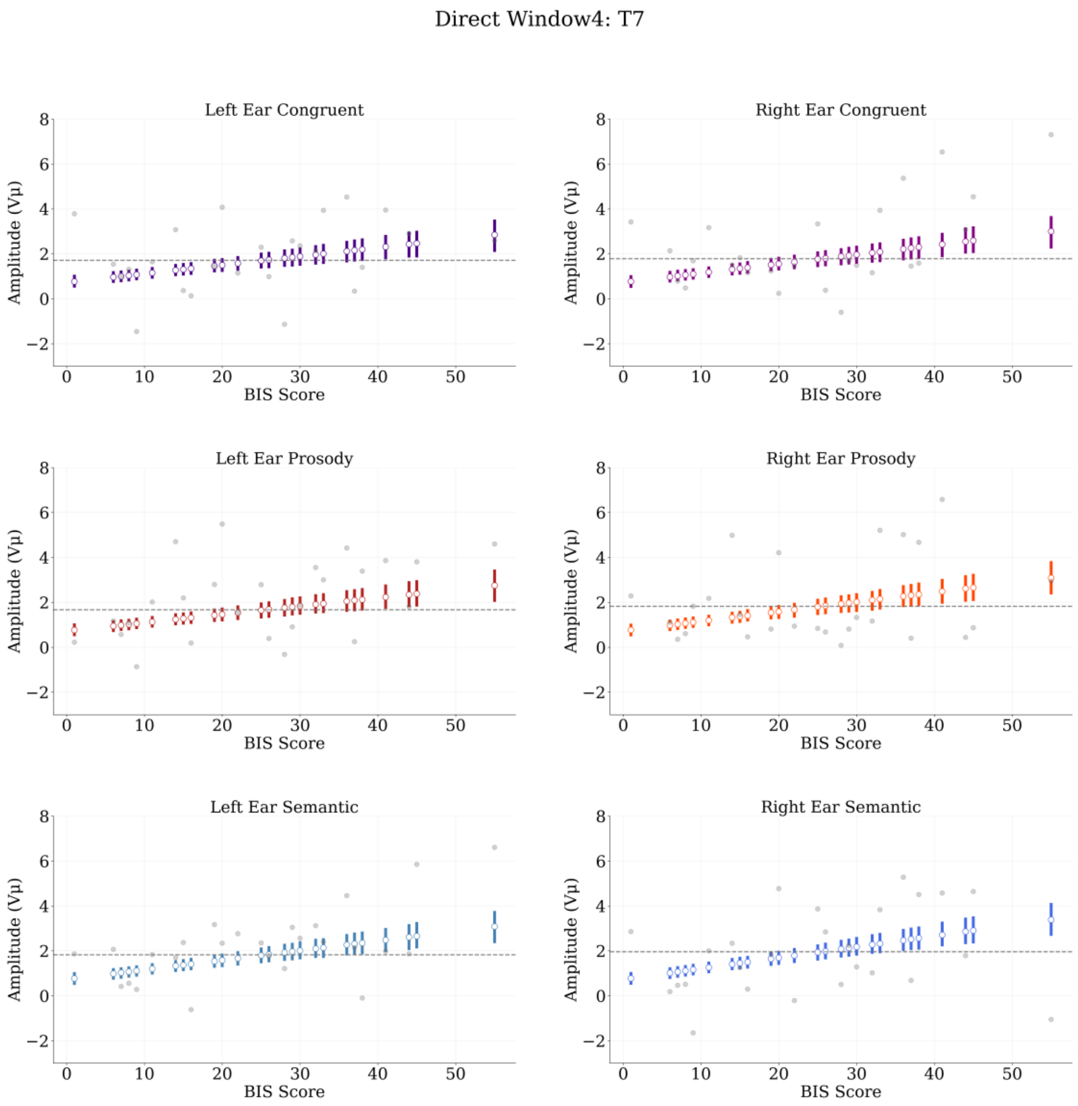


**Supplementary Figure 2.1**. Direct-threat regressions at electrode T7, Window4 (500-750ms). White circles represent posterior means by BIS score. Bars represent highest density intervals (HDIs). Grey dots are mean amplitudes by BIS score. Dashed grey line indicates posterior median BIS.

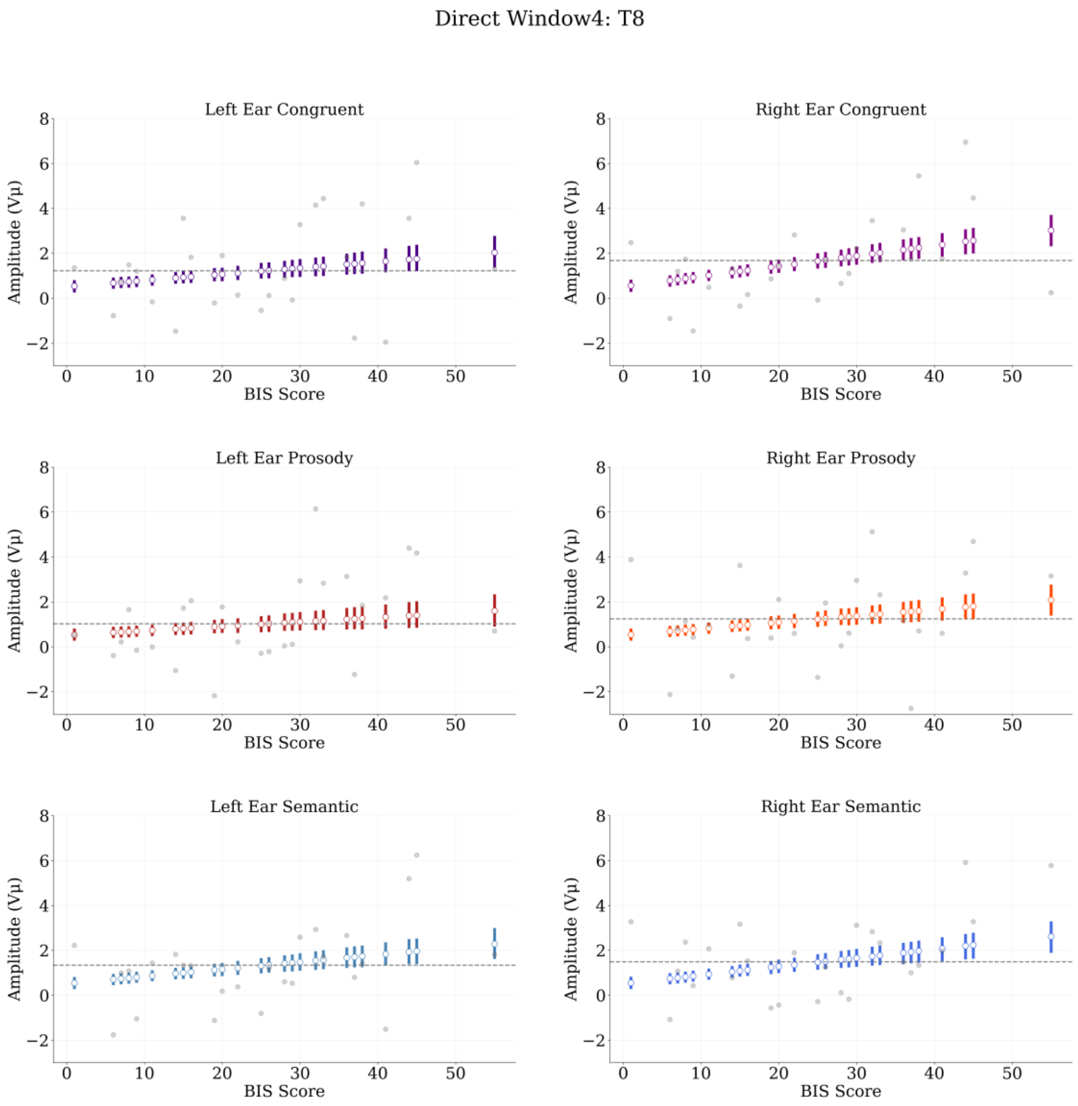


**Supplementary Figure 2.2**. Direct-threat regressions at electrode T8, Window4 (500-750ms). White circles represent posterior means by BIS score. Bars represent highest density intervals (HDIs). Grey dots are mean amplitudes by BIS score. Dashed grey line indicates posterior median BIS.

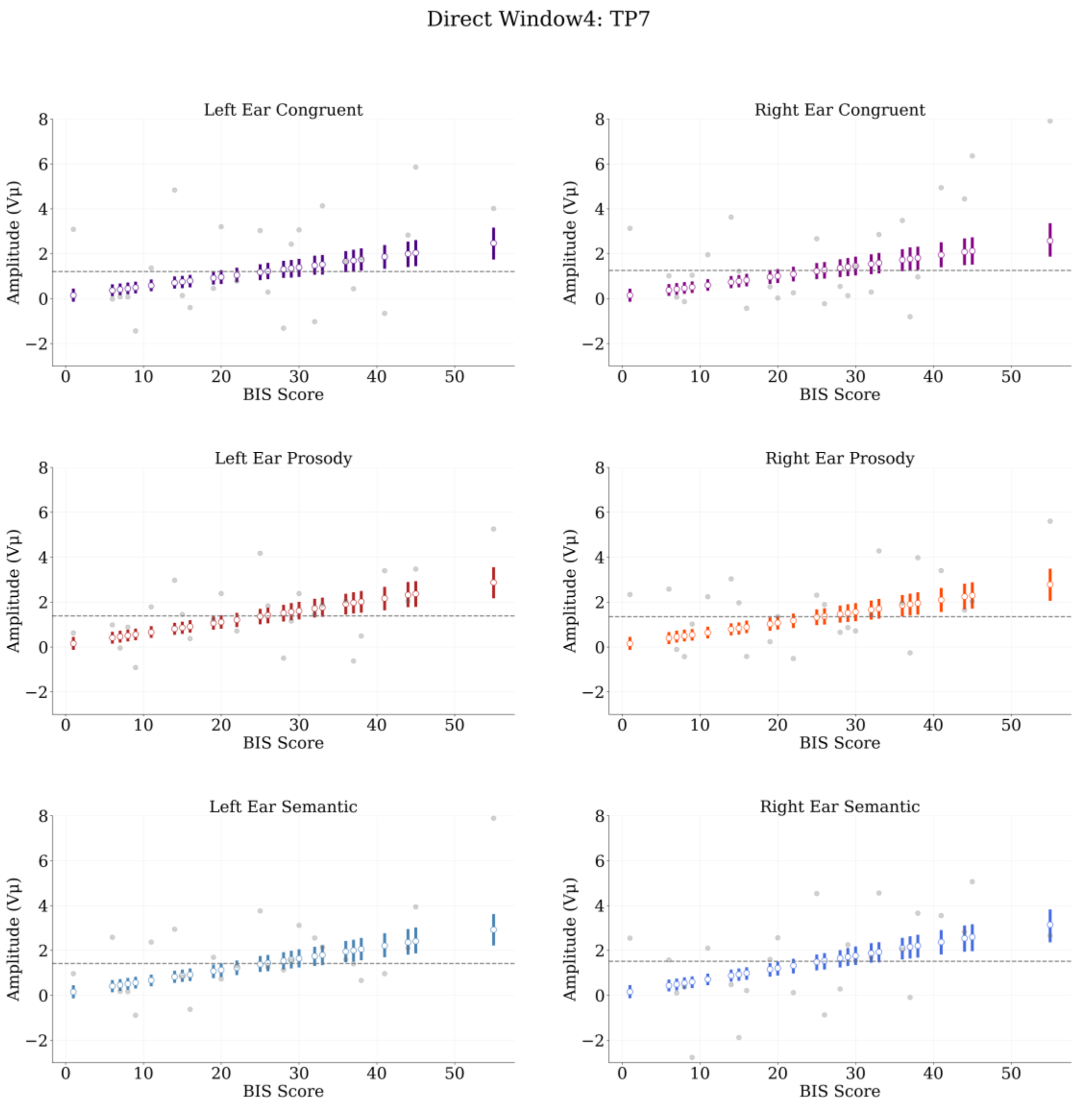


**Supplementary Figure 2.3**. Direct-threat regressions at electrode TP7, Window4 (500-750ms). White circles represent posterior means by BIS score. Bars represent highest density intervals (HDIs). Grey dots are mean amplitudes by BIS score. Dashed grey line indicates posterior median BIS.

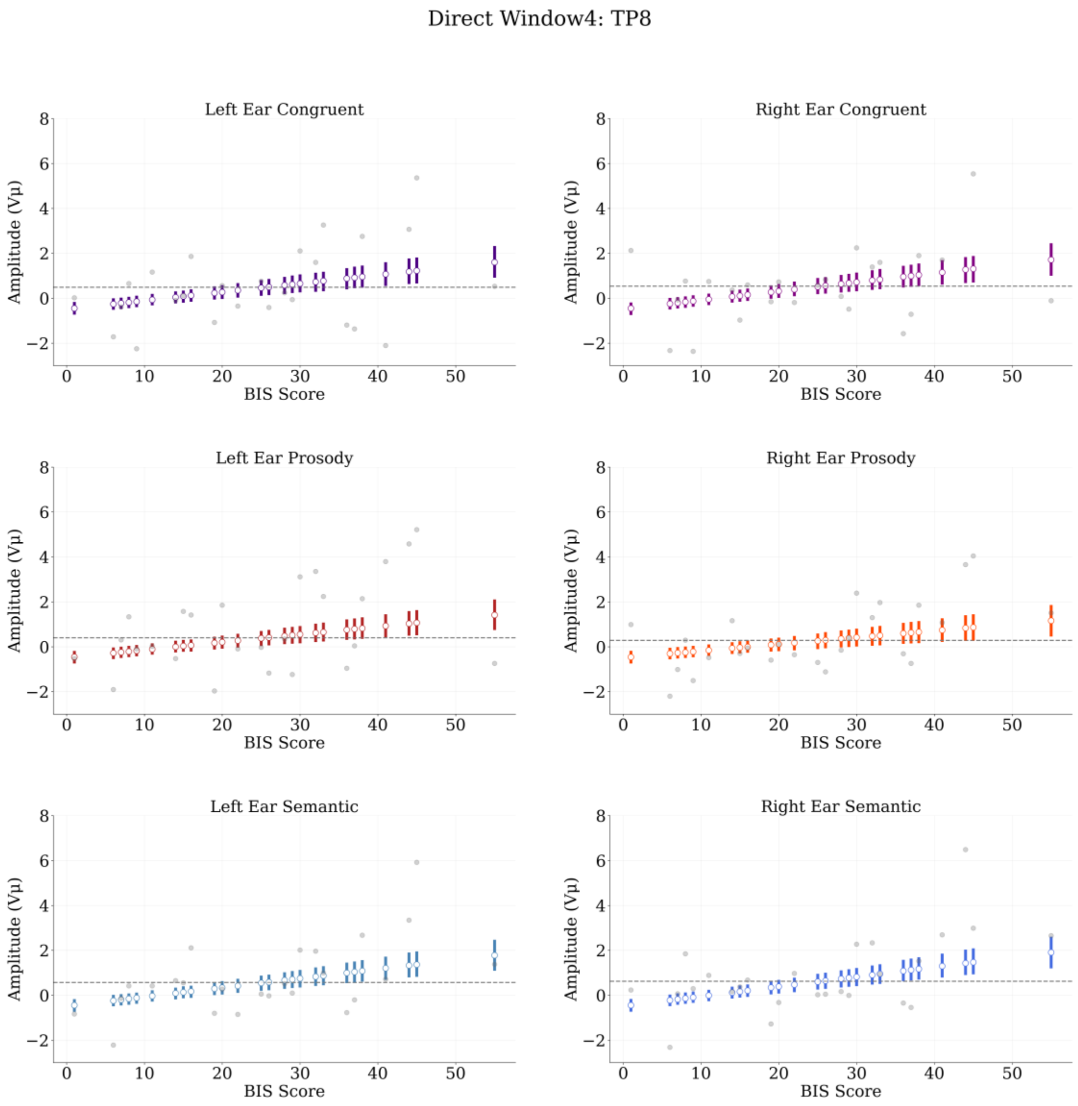


**Suppementary Figure 2.4**. Direct-threat regressions at electrode TP8, Window4 (500-750ms). White circles represent posterior means by BIS score. Bars represent highest density intervals (HDIs). Grey dots are mean amplitudes by BIS score. Dashed grey line indicates posterior median BIS.

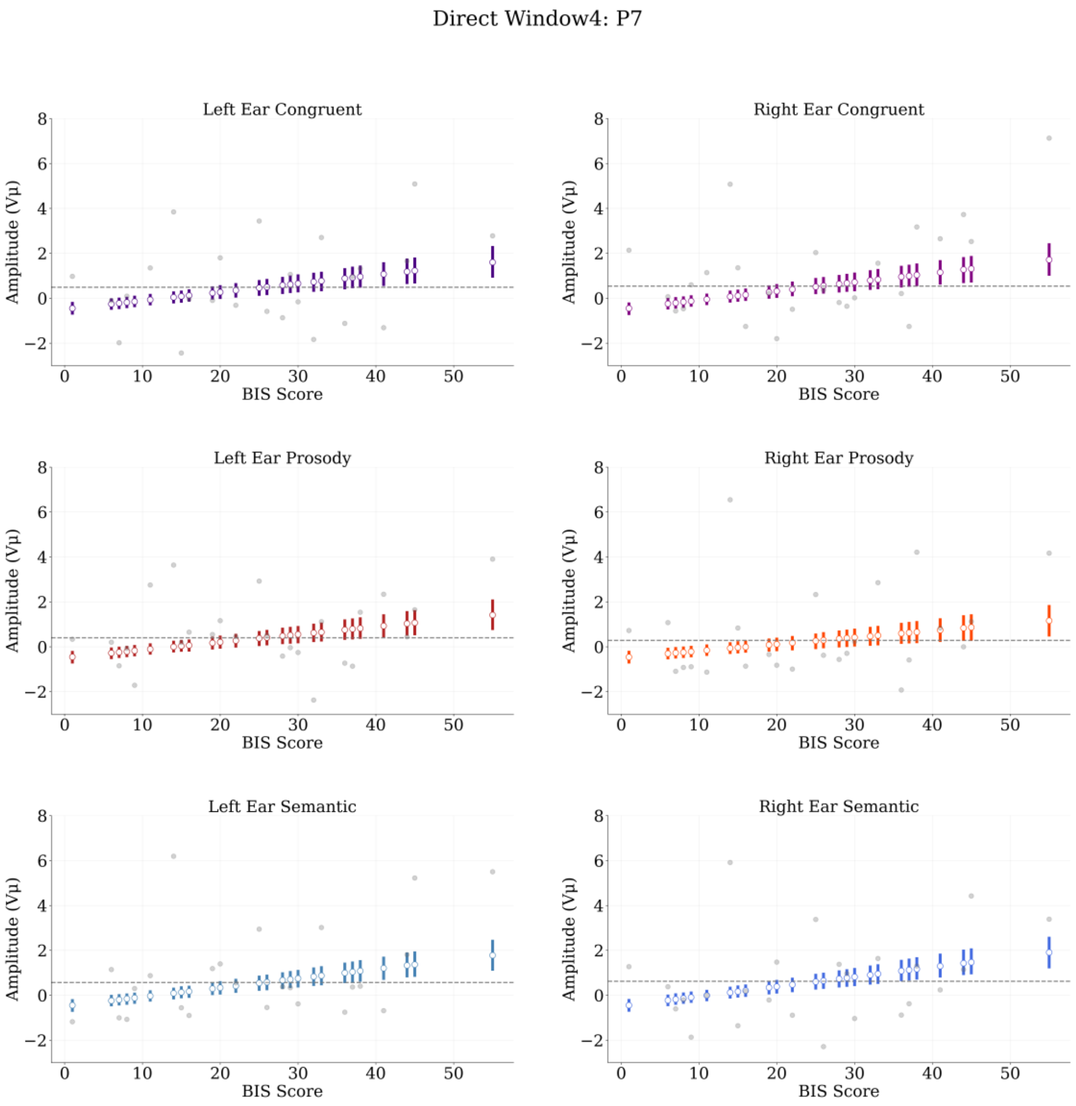


**Supplementary Figure 2.5**. Direct-threat regressions at electrode P7, Window4 (500-750ms). White circles represent posterior means by BIS score. Bars represent highest density intervals (HDIs). Grey dots are mean amplitudes by BIS score. Dashed grey line indicates posterior median BIS.

Direct Window4: P8

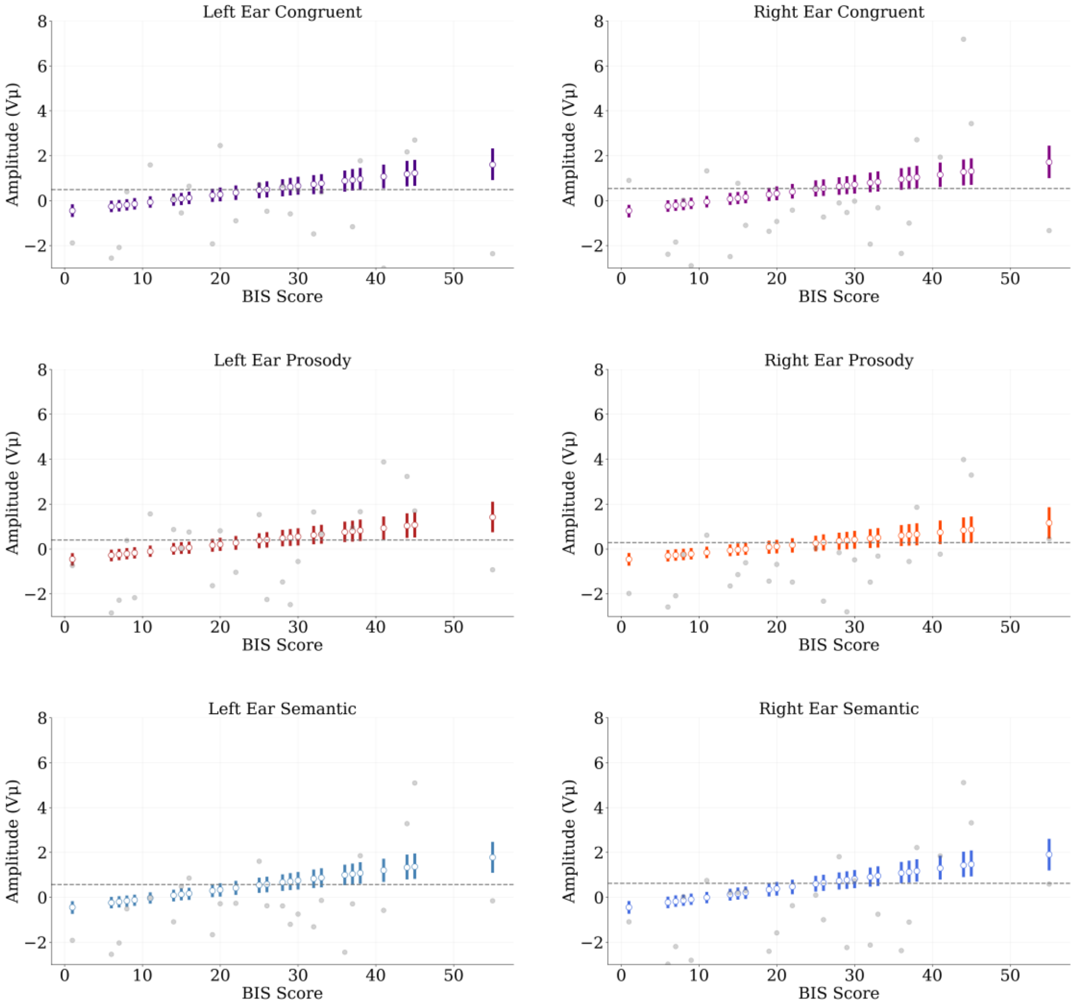


**Supplementary Figure 2.6**. Direct-threat regressions at electrode P8, Window4 (500-750ms). White circles represent posterior means by BIS score. Bars represent highest density intervals (HDIs). Grey dots are mean amplitudes by BIS score. Dashed grey line indicates posterior median BIS.

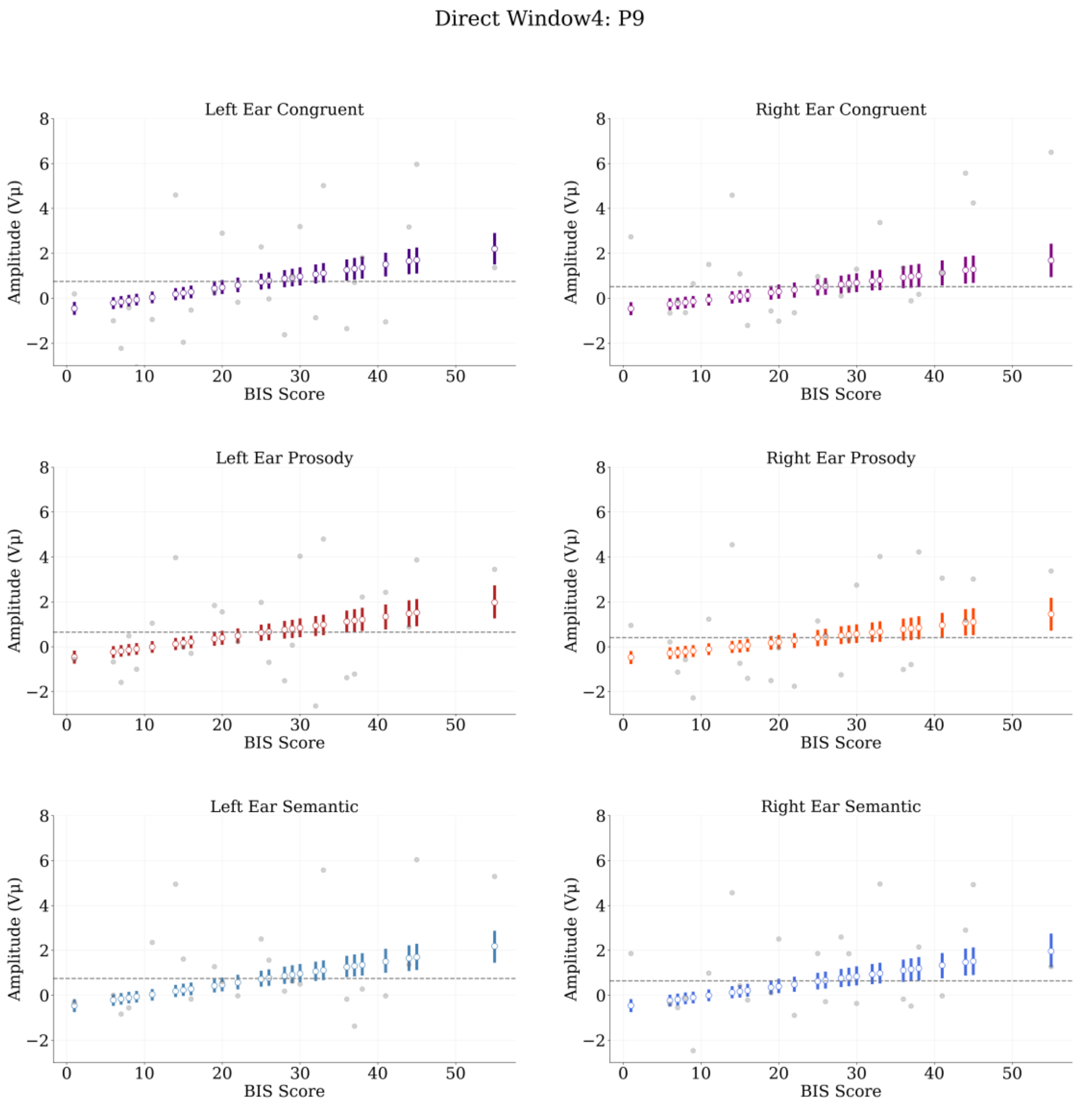


**Supplementary Figure 2.7**. Direct-threat regressions at electrode P9, Window4 (500-750ms). White circles represent posterior means by BIS score. Bars represent highest density intervals (HDIs). Grey dots are mean amplitudes by BIS score. Dashed grey line indicates posterior median BIS.

Direct Window4: P10

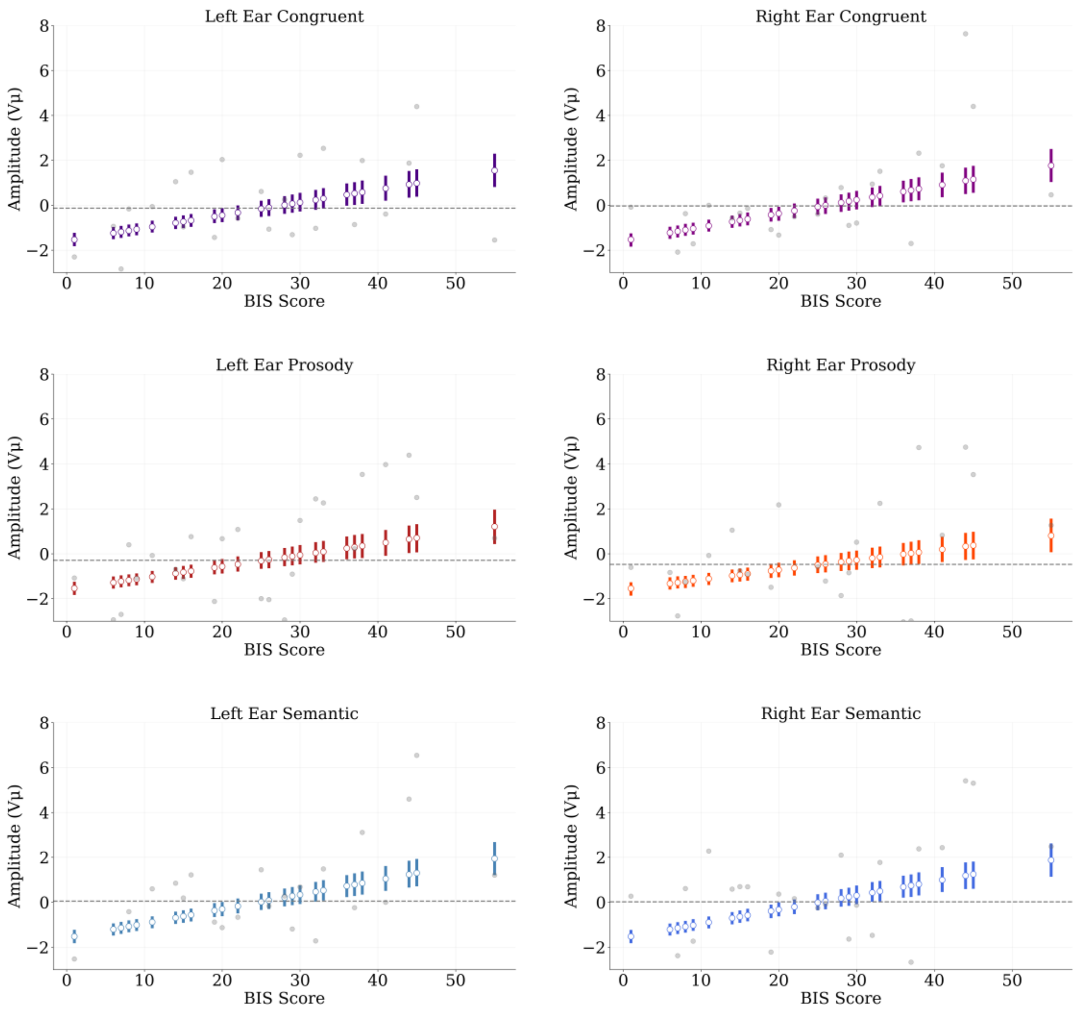


**Supplemenary Figure 2.8**. Direct-threat regressions at electrode P10, Window4 (500-750ms). White circles represent posterior means by BIS score. Bars represent highest density intervals (HDIs). Grey dots are mean amplitudes by BIS score. Dashed grey line indicates posterior median BIS.
