## Supplement 3 for "Trait Anxiety Effects on Late Phase Threatening Speech Processing: Evidence from Electroencephalography"

Supplementary figures below show examples for all four time-windows using electrode TP7 as an example: aimplitide as a function of BIS score, ear and sentence type. Figures from all other relevant electrodes can be found in our OSF repository (https://osf.io/n5b6h/). Supplementary figures 3.1 to 3.4 reflect the direct threat condition and 3.5 to 3.8, indirect threat.

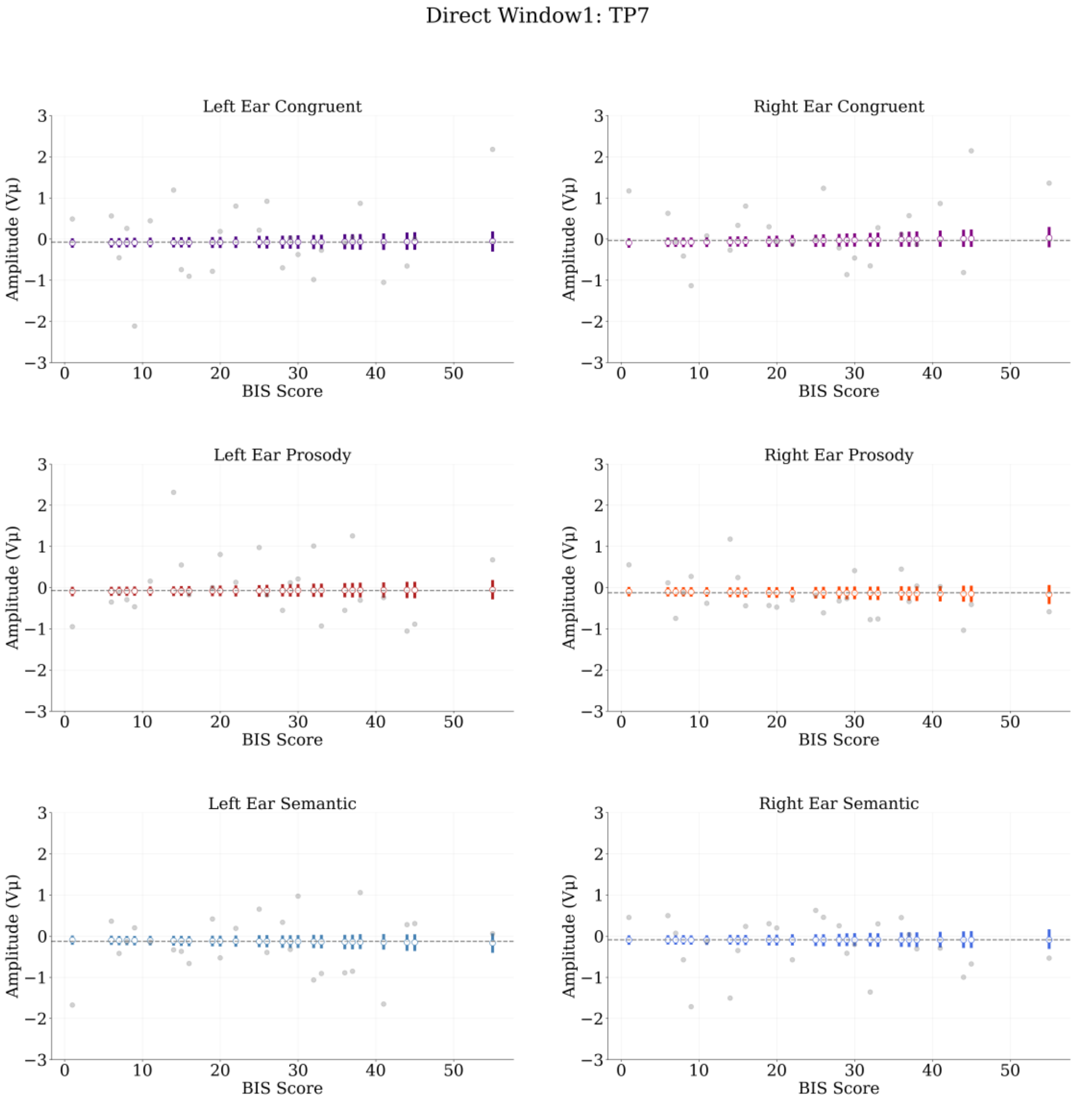


**Supplementary Figure 3.1**. Direct-threat regressions at electrode TP7, Window1 (50-150ms). White circles represent posterior means by BIS score. Bars represent highest density intervals (HDIs). Grey dots are mean amplitudes by BIS score. Dashed grey line indicates posterior median BIS.

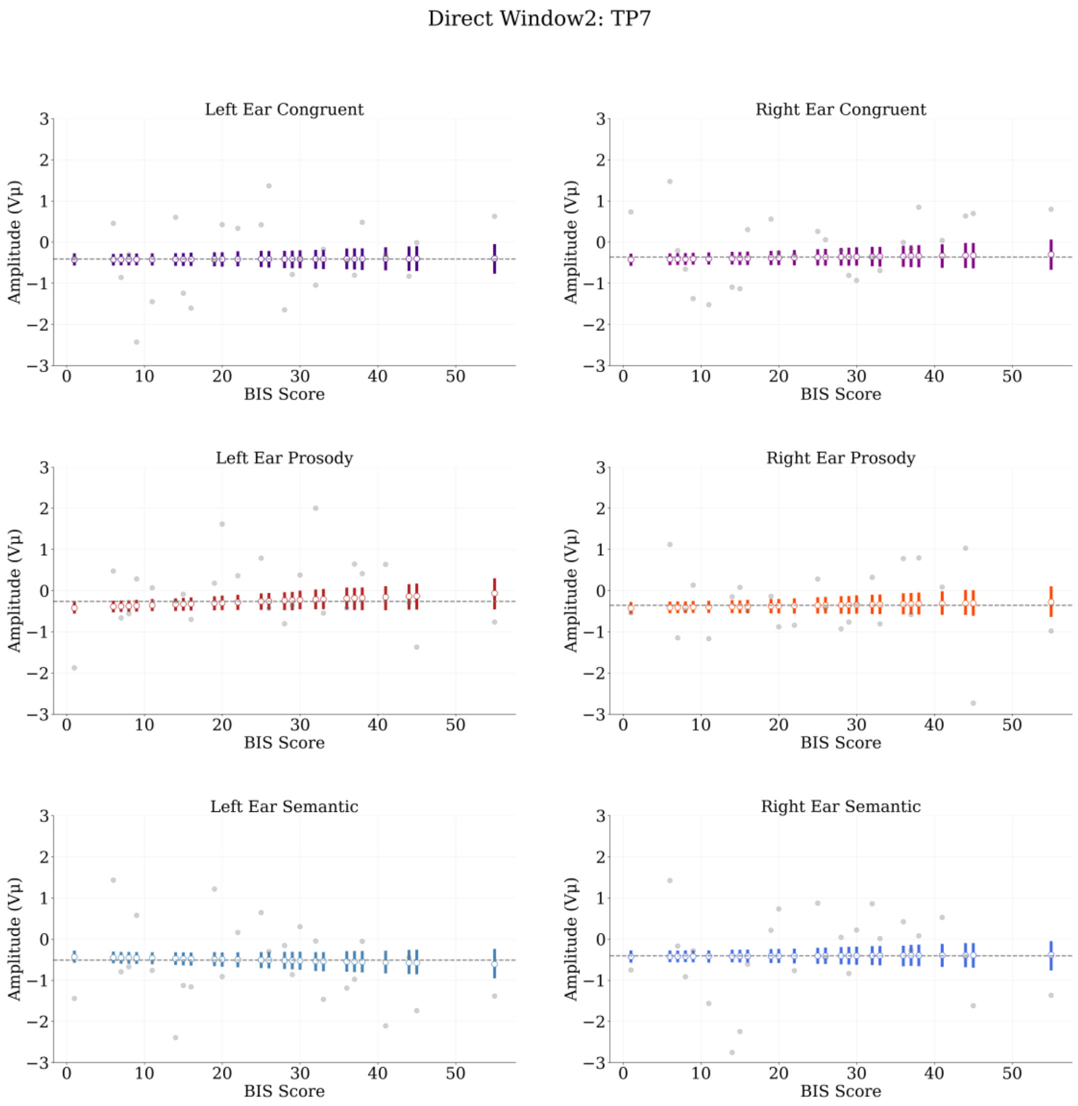


**Supplementary Figure 3.2**. Direct-threat regressions at electrode TP7, Window2 (150-250ms). White circles represent posterior means by BIS score. Bars represent highest density intervals (HDIs). Grey dots are mean amplitudes by BIS score. Dashed grey line indicates posterior median BIS.

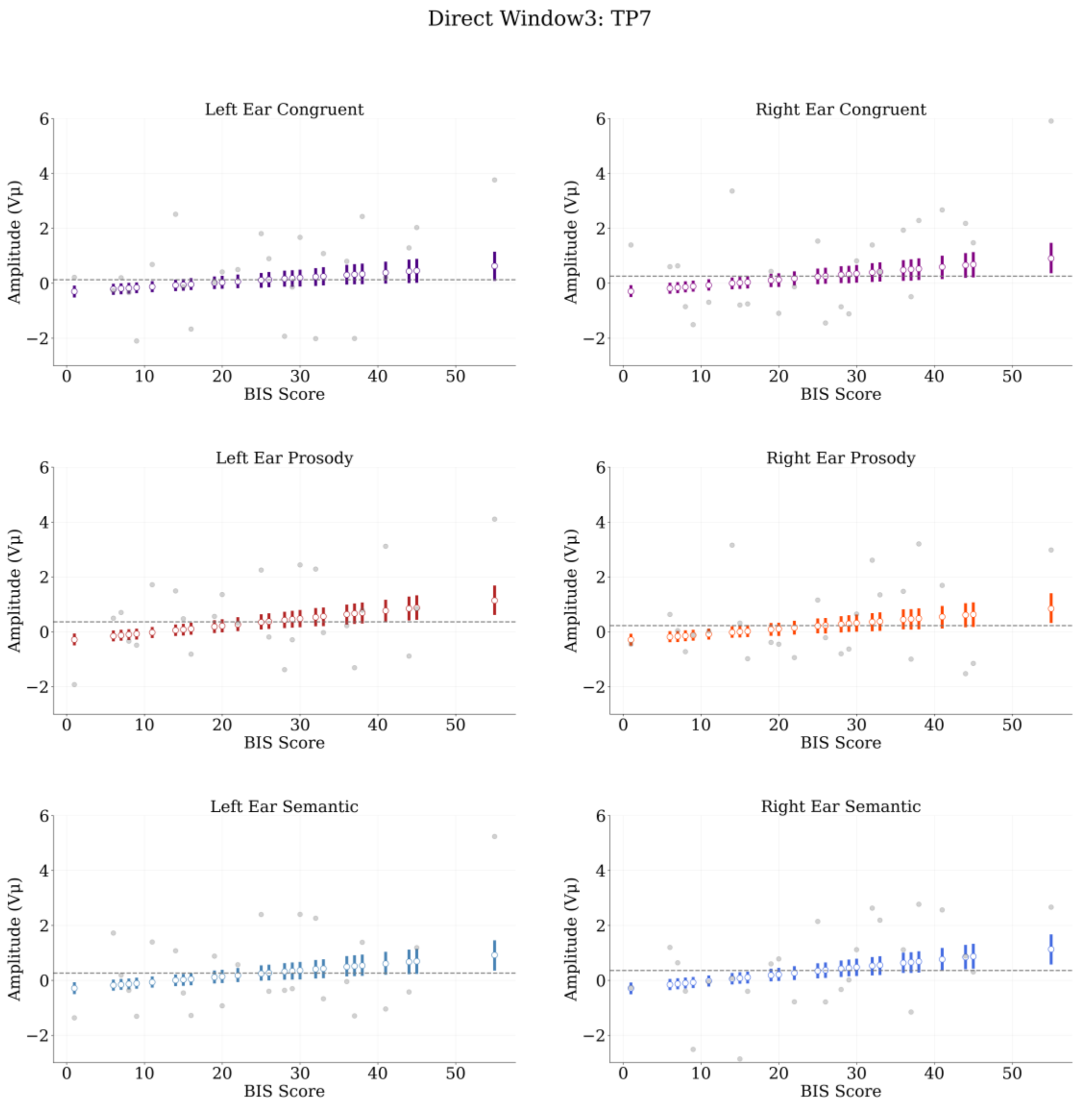


**Supplementary Figure 3.3**. Direct-threat regressions at electrode TP7, Window3 (250-500ms). White circles represent posterior means by BIS score. Bars represent highest density intervals (HDIs). Grey dots are mean amplitudes by BIS score. Dashed grey line indicates posterior median BIS.

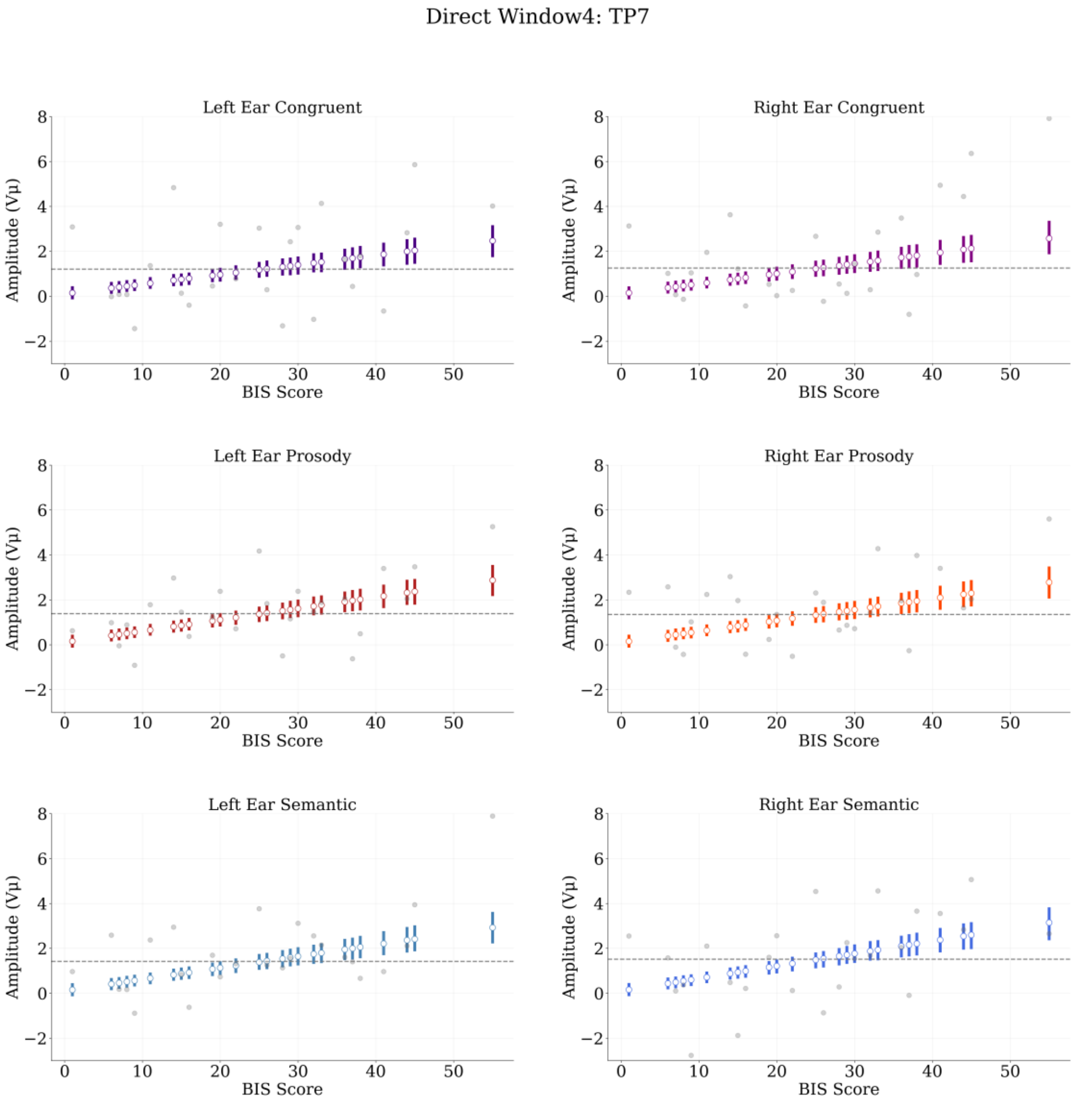


**Supplementary Figure 3.4**. Direct-threat regressions at electrode TP7, Window4 (500-750ms). White circles represent posterior means by BIS score. Bars represent highest density intervals (HDIs). Grey dots are mean amplitudes by BIS score. Dashed grey line indicates posterior median BIS.

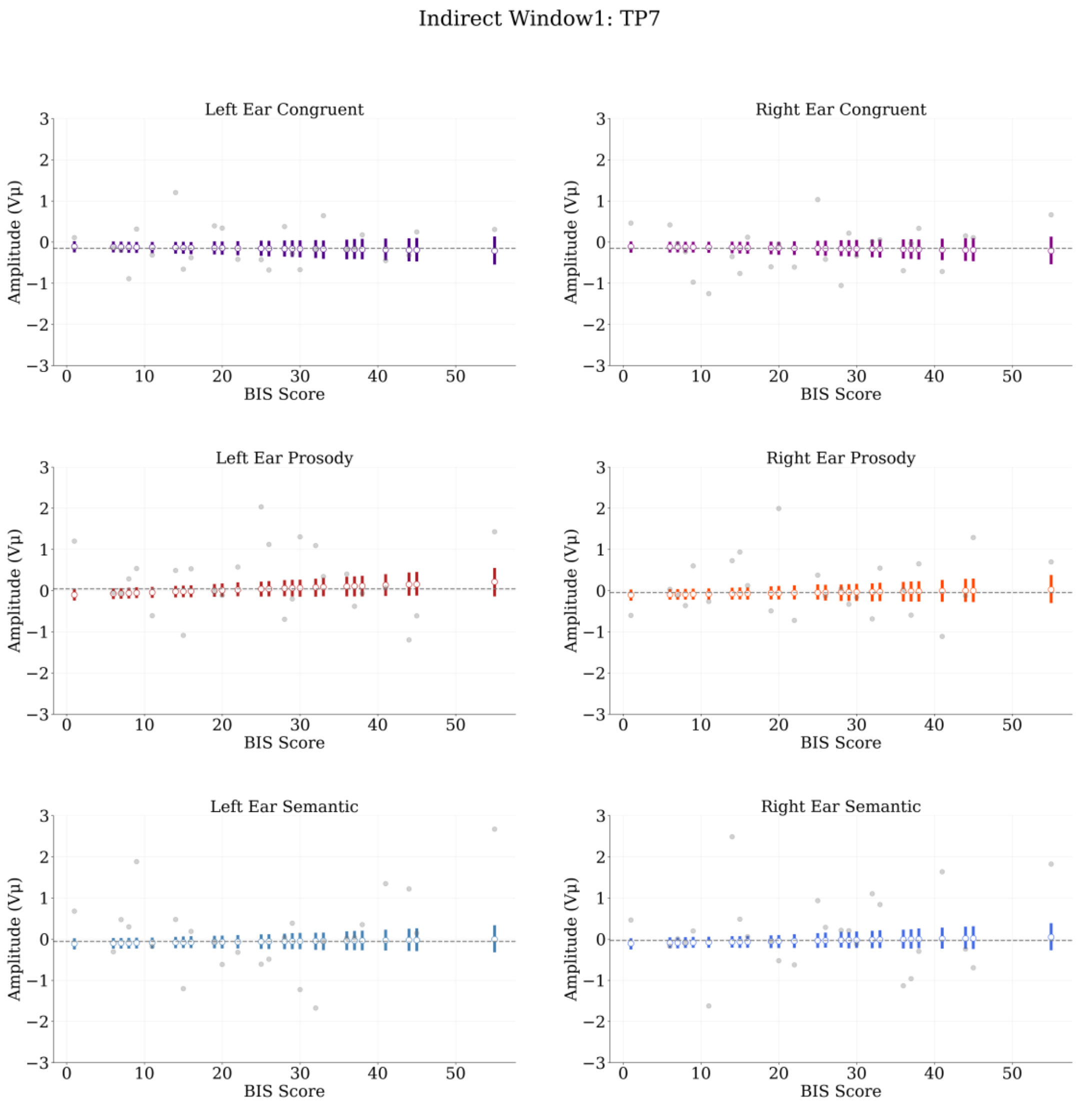


**Supplementary Figure 3.5**. Indirect-threat regressions at electrode TP7, Window1 (50-150ms). White circles represent posterior means by BIS score. Bars represent highest density intervals (HDIs). Grey dots are mean amplitudes by BIS score. Dashed grey line indicates posterior median BIS.

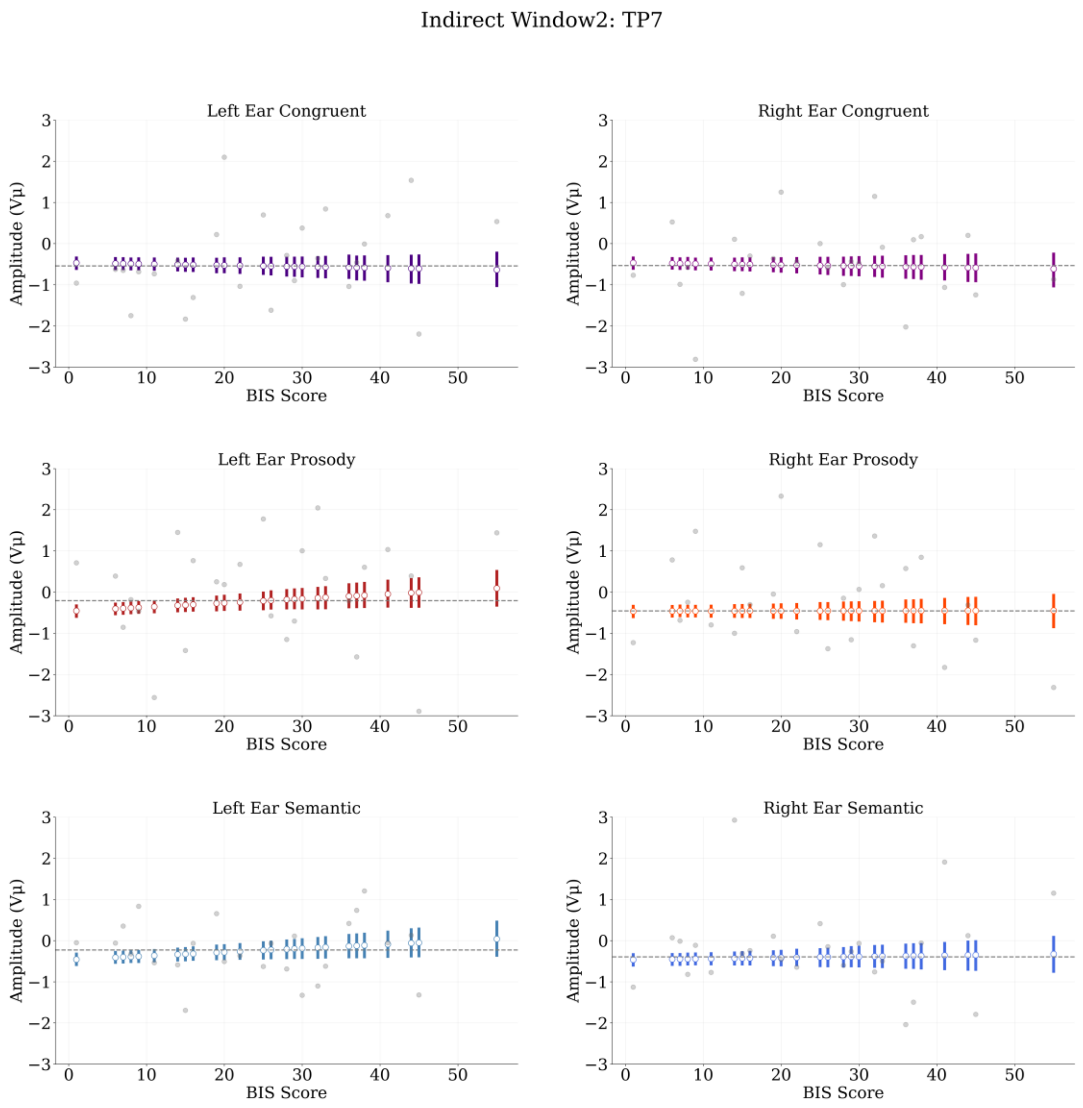


**Supplementary Figure 3.6**. Indirect-threat regressions at electrode TP7, Window2 (150-250ms). White circles represent posterior means by BIS score. Bars represent highest density intervals (HDIs). Grey dots are mean amplitudes by BIS score. Dashed grey line indicates posterior median BIS.

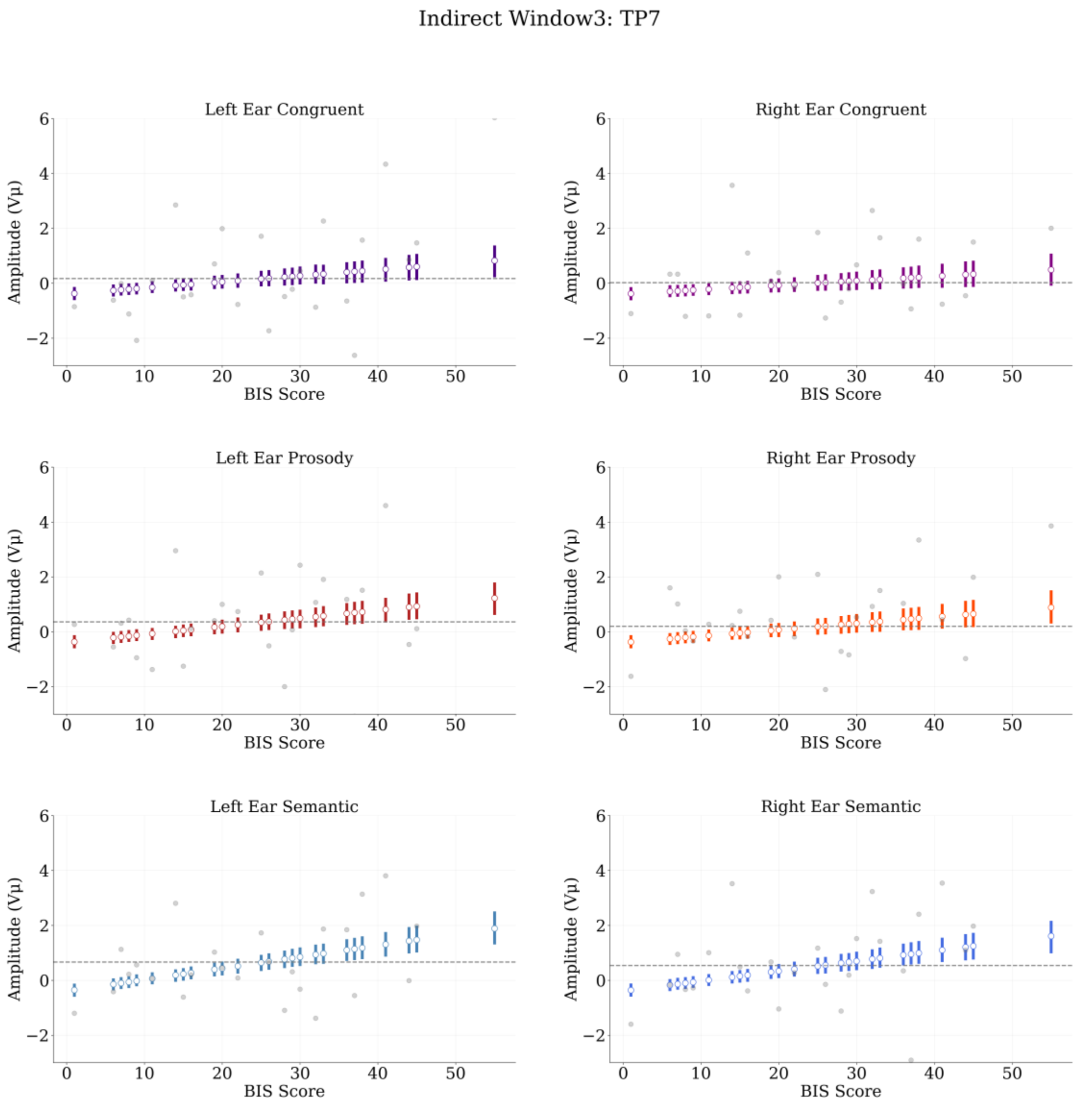


**Supplementary Figure 3.7**. Indirect-threat regressions at electrode TP7, Window3 (250-500ms). White circles represent posterior means by BIS score. Bars represent highest density intervals (HDIs). Grey dots are mean amplitudes by BIS score. Dashed grey line indicates posterior median BIS.

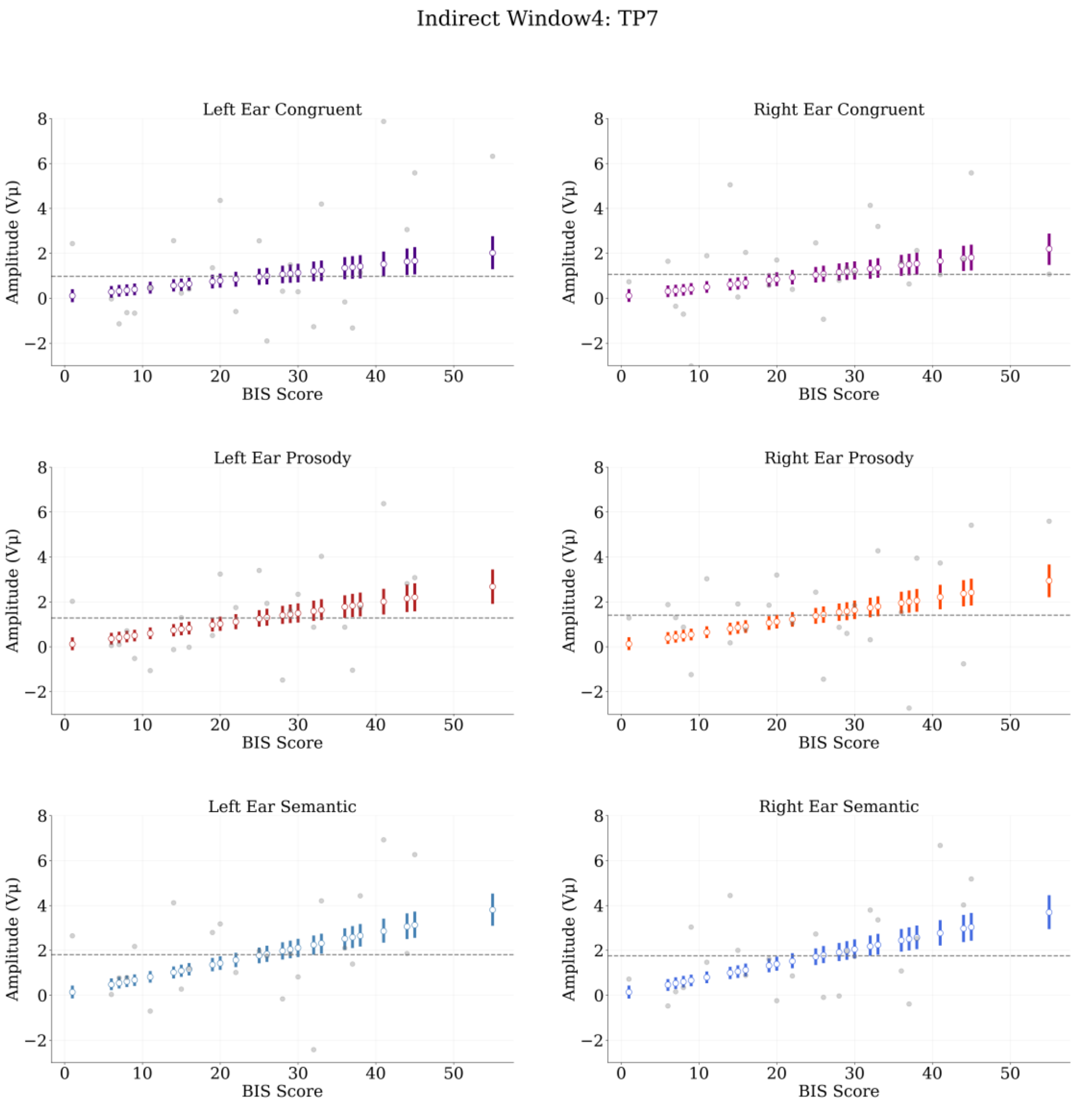


**Supplementary Figure 3.8**. Indirect-threat regressions at electrode TP7, Window4 (500-750ms). White circles represent posterior means by BIS score. Bars represent highest density intervals (HDIs). Grey dots are mean amplitudes by BIS score. Dashed grey line indicates posterior median BIS.
