## Supplement 4 for "Trait Anxiety Effects on Late Phase Threatening Speech Processing: Evidence from Electroencephalography"

**Supplement 4: List Effects**

A minor coding error occurred in one of the experimental lists during randomisation of stimuli. Here, we report analyses to check whether this issue systematically affected the results of the study. As described in the main text, we constructed two lists, reflecting the counterbalanced order of the direct-threat and indirect-threat tasks. Participants for List A received the direct-threat task first, and participants for List B received the indirect-threat task first. We intended that each threatening stimulus would be presented to each ear exactly once per participant for each of these two tasks, thus exactly balancing laterality of presentation at the item level. This was the case for List A (both direct-threat and indirect-threat tasks) and for direct-threat task in List B. However, for the indirect-threat task in List B half of the participants received the same sentence at a given ear twice for some sentences (and none for the other ear). This led to some fluctuation in the number of trials per condition × ear × subject in the indirect threat task, for list B only: (e.g. 59 Prosody instead of 54 Prosody). For example, Subject 1b had 59 Prosody in left ear and 49 Prosody in right ear, instead of the 54 intended. If this deviation from our intended strict balance in laterality of stimulus presentation undermined our ability to detect systematic effects of interest, this should be reflected in systematically reduced magnitude of these effects in indirect-threat condition, for list B only.

To address this issue we conducted additional analyses, as described in the main text but also including list as a factor (List A and List B). Posterior distributions indicate that some variability occurs between lists, but it does not reflect a systematic effect. Rather, this variability is consistent with the un-pooling of participants by list. Supplementary Figures 4.1 and 4.2 summarise effects of direct-threat, where lists were presented as intended and thus should reflect just the consequences of un-pooling by list. The general pattern of increased amplitude as a function of BIS remains, although slightly diminished in some conditions, such as Congruent at right ear (TP7 List A) or Congruent at left ear (TP8 ListA, TP7 ListB), and strongly diminished only on Prosody right ear at TP8 List A. As this follows no specific pattern, the most straightforward interpretation is that it corresponds to normal variability by list.

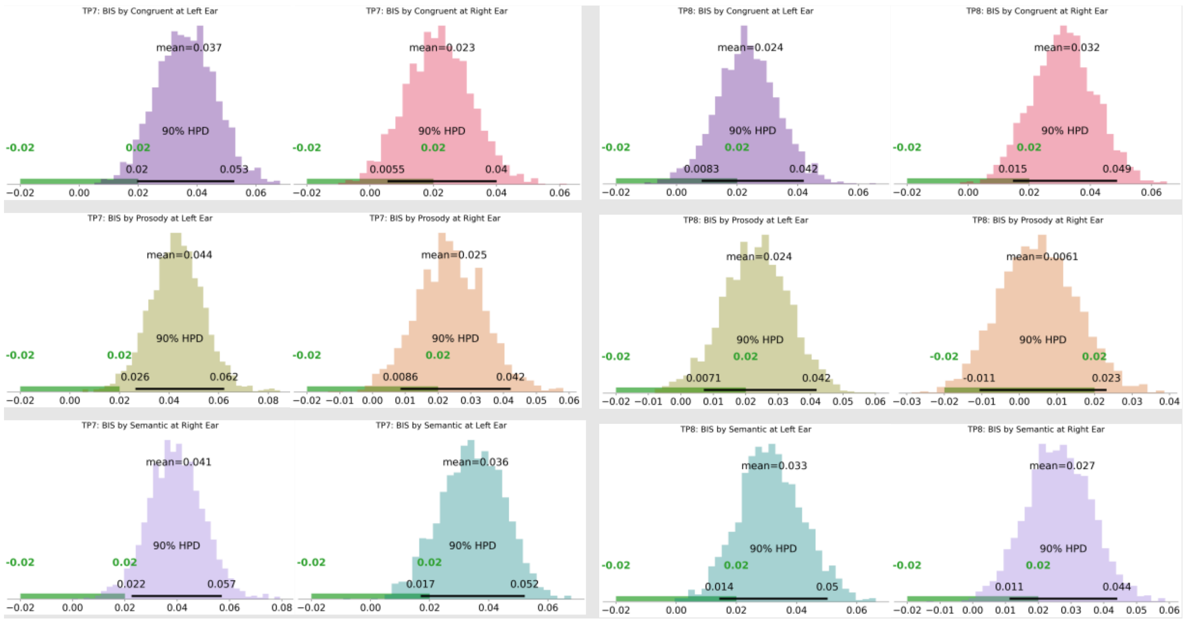


**Supplementary Figure 4.1**. Direct-threat List A estimates. Plots shows summaries of posterior distributions from TP7 (left panels) and TP8 (right panels) electrodes. Black band: 90% Highest posterior density (HDI). Green bands: 2SD regions of practical equivalence (ROPE).

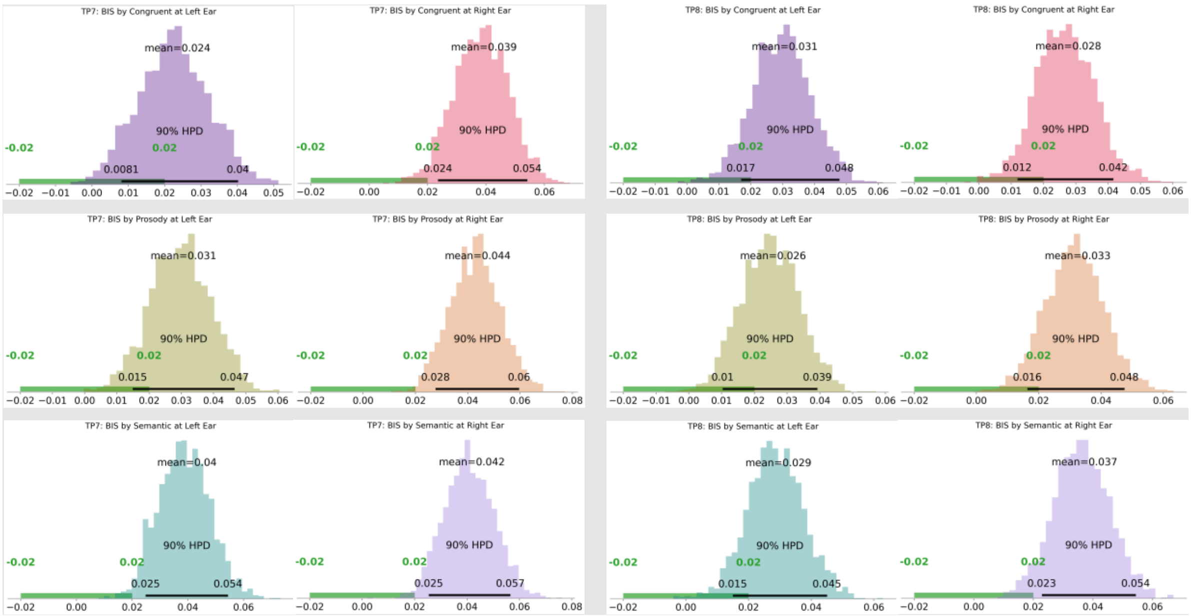


**Supplementary Figure 4.2**. Direct-threat List B estimates. Plots shows summaries of posterior distributions from TP7 (left panels) and TP8 (right panels) electrodes. Black band: 90% Highest posterior density (HDI). Green bands: 2SD regions of practical equivalence (ROPE).

Supplementary Figures 4.3 and 4.4 summarise results from indirect-threat, where List B was not precisely balanced as intended. Note that the same pattern remains, with the difference that the effect is somewhat reduced at TP8 for Prosody at both left and right ears. This, however, does not represent a serious variation as the effect did not greatly decrease in other right-lateral electrodes (e.g. T8) near TP8 (posterior distributions from all electrodes can be found in our OSF repository (https://osf.io/n5b6h/). As this pattern does not systematically differ for list B, we can conclude that this minor deviation from exact ear balance at the item level does not undermine our findings.

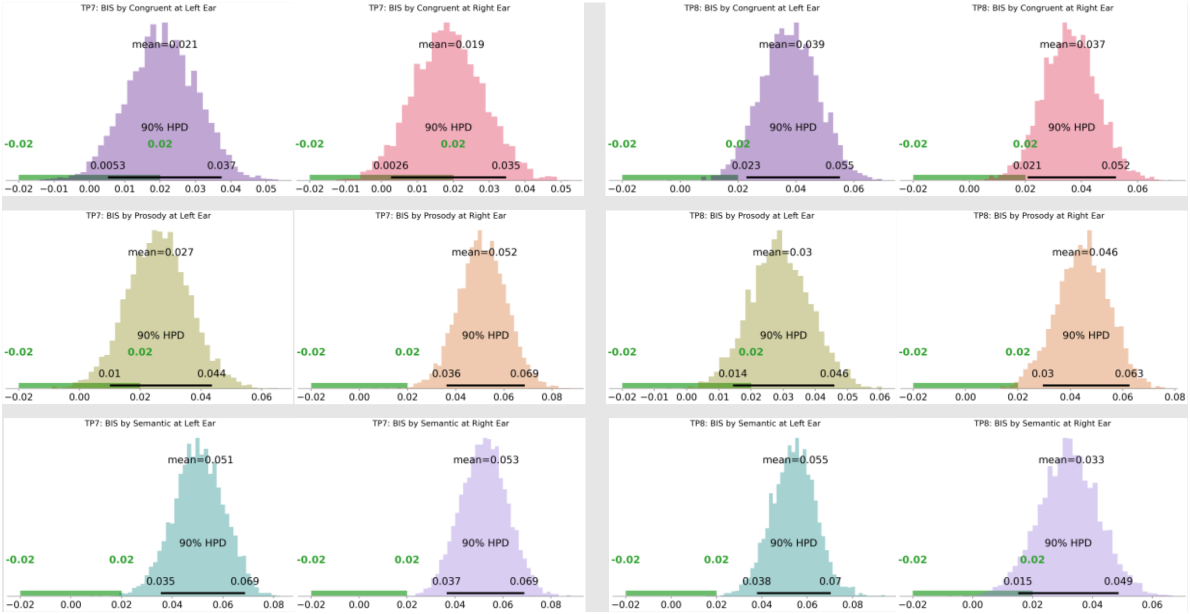


**Supplementary Figure 4.3.** Indirect-threat List B estimates. Plots shows summaries of posterior distributions from TP7 (left panels) and TP8 (right panels) electrodes. Black band: 90% Highest posterior density (HDI). Green bands: 2SD regions of practical equivalence (ROPE).

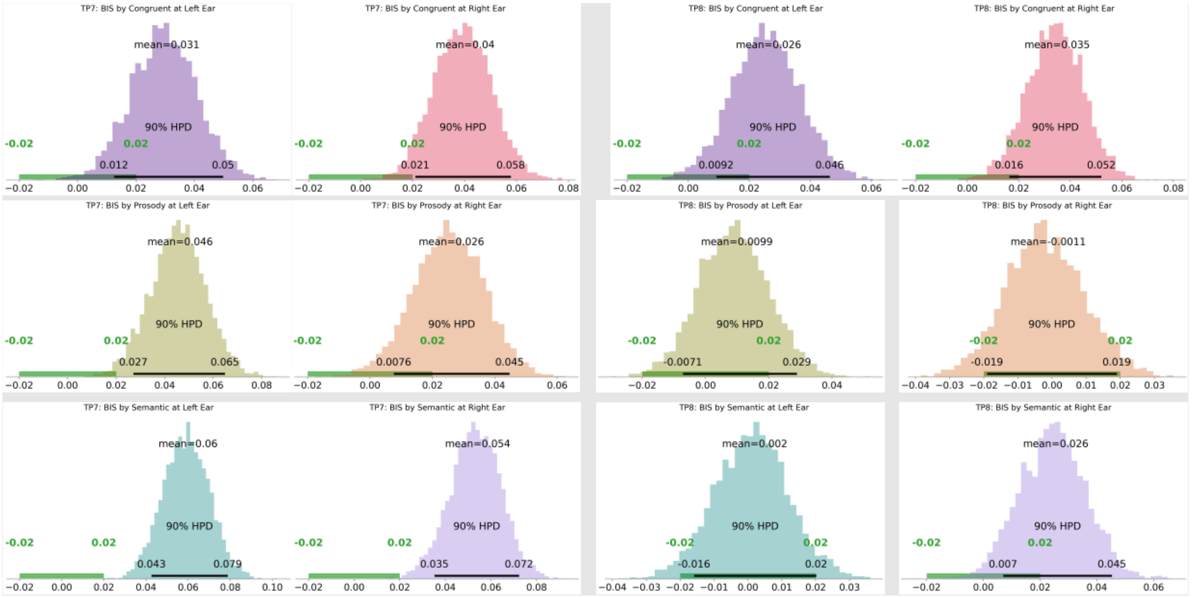


**Supplementary Figure 4.4**. Indirect-threat List A estimates. Plots shows summaries of posterior distributions from TP7 (left panels) and TP8 (right panels) electrodes. Black band: 90% Highest posterior density (HDI). Green bands: 2SD regions of practical equivalence (ROPE).
